# Denisovan ancestry and population history of early East Asians

**DOI:** 10.1101/2020.06.03.131995

**Authors:** Diyendo Massilani, Laurits Skov, Mateja Hajdinjak, Byambaa Gunchinsuren, Damdinsuren Tseveendorj, Seonbok Yi, Jungeun Lee, Sarah Nagel, Birgit Nickel, Thibaut Devièse, Tom Higham, Matthias Meyer, Janet Kelso, Benjamin M. Peter, Svante Pääbo

## Abstract

We present analyses of the genome of a ~34,000-year-old hominin skull cap discovered in the Salkhit Valley in North East Mongolia. We show that this individual was a female member of a modern human population that, following the split between East and West Eurasians, experienced substantial gene flow from West Eurasians. Both she and a 40,000-year-old individual from Tianyuan outside Beijing carried genomic segments of Denisovan ancestry. These segments derive from the same Denisovan admixture event(s) that contributed to present-day mainland Asians but are distinct from the Denisovan DNA segments in present-day Papuans and Aboriginal Australians.

## Introduction

Modern humans may have been present in East Asia as early as 80,000 years before present (BP) (*1*, *2*) but how they eventually settled in the region remains largely unknown (*3*–*5*). To date, genomic data in East Asia exist from only a single Paleolithic individual, a ~40,000-year-old individual from Tianyuan Cave in the Beijing area in China (*6*). This individual was more closely related to present-day East Asians than to ancient Europeans, but surprisingly shared more alleles with a ~35,000-year-old individual from Belgium (Goyet Q116-1) than with other ancient Europeans (*7*). In Siberia, that neighbors East Asia to the north, four modern human individuals older than 20,000 years BP have been studied: a ~45,000-year-old individual from Ust’-Ishim in West Siberia who did not contribute ancestry to present-day populations (*8*); a ~24,000-year-old individual from Mal’ta in South-Central Siberia who was more related to Western Europeans than to East Asians and was part of a population that contributed approximatively one third of the ancestry of present-day Native Americans (*9*); and two ~31,000-year-old individuals from the Yana Rhinoceros Horn Site (RHS) in North East Siberia who show affinities to early modern humans in both West and East Eurasia (*10*).

In 2006 a hominin skull cap was discovered during mining operations in the Salkhit Valley in Northeastern Mongolia (48°16’17.9”N, 112°21’37.9”E) (*11*) (Fig. 1a). Its unusual morphology led to that it was referred to as *Mongolanthropus* (*11*) and later to suggestions that it was affiliated with Neandertals or *Homo erectus* (*12*–*14*). Recently, it was radiocarbon dated to 33,900-34,950 calibrated years BP (95% probability interval) and its mitochondrial (mt) DNA was shown to belong to a group of mtDNAs that are widespread in Eurasia today (*15*).

**Fig. 1.**
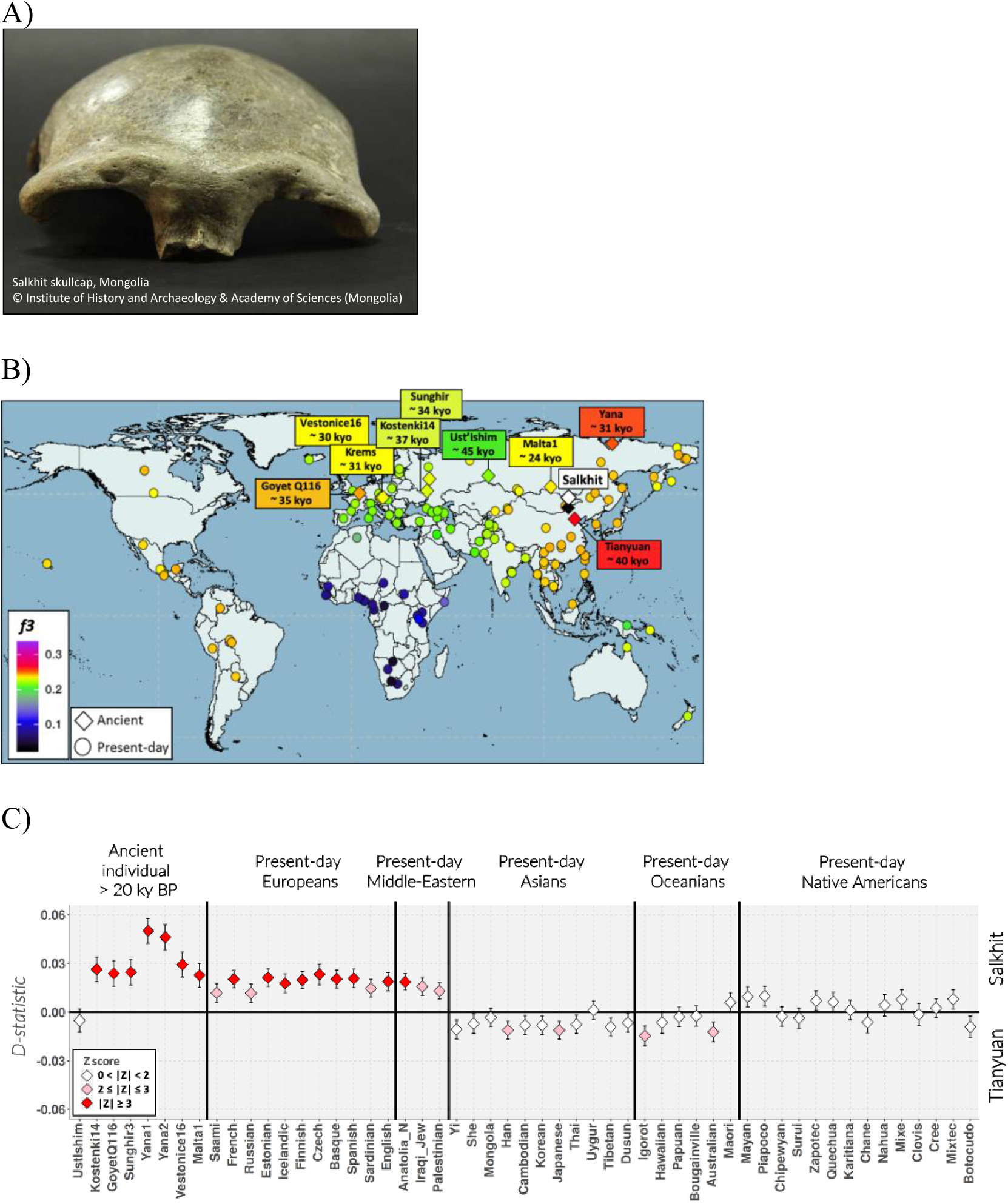
The Salkhit individual and her relationship to ancient and present-day humans. **A)** The Salkhit skull cap. **B)** Heat map illustrating the genetic similarity between the Salkhit individual and modern humans from Eurasia older than 20,000 years (diamonds) as well as present-day human populations (circles) determined by *f3-statistic* of the form *f3(Salkhit, X; Mbuti)*. The warmer the color, the higher is the genetic similarity between the Salkhit individual and a population/individual. **C)** Relative amounts of allele sharing between the Salkhit and Tianyuan genomes and ancient and present-day humans determined by *D-statistics* of the form *D(Salkhit, Tianyuan, X, Mbuti).* The *D-*statistic is positive when the individual/population shares more alleles with the Salkhit individual than with the Tianyuan individual. The colors of the diamonds indicate whether the Z-score is significant (red), weakly significant (pink), or not significant (white).

## Results and Discussion

### Data quality

To study the Salkhit individual’s nuclear genome, we generated shotgun sequence data from six DNA libraries prepared from bone powder from the Salkhit skull cap (*16*). Between 0.6% and 5.6% of the DNA fragments in the libraries mapped uniquely to the human reference genome (hg19) (Table S2). Apparent cytosine (C) to thymine (T) substitutions, that are common at the ends of ancient DNA molecules as a result of deamination of cytosine residues (*17*), affect between 23% and 40% of the 5’-ends, and between 13% and 25% of the 3’-ends in the six libraries, indicating the presence of ancient hominin DNA (Table S3, Fig S1-2). Using C to T substitution patterns (*18*), we estimate the extent of contamination by present-day human DNA to vary between ~5% and ~50% among the libraries (Table S3). Because of the high level of human contamination subsequent analyses (unless specified otherwise) were performed using only DNA sequences showing evidence of cytosine deamination at their first or last base (referred as “deaminated fragments”), among which contamination is estimated to 1-3% (Table S4).

### Sex determination and lineage assignment

The average coverage of the autosomes is not significantly different from that of the X chromosome, indicating that, despite the robust morphology of the skull, the Salkhit individual was female (Table S8, Fig. S6). To determine to which major group of hominins she belonged, we estimated the percentage of derived alleles shared with the genomes of a present-day human (Mbuti, HGDP00982) (*19*), a Neandertal (*Denisova 5*) (*20*), and a Denisovan (*Denisova 3*) (*21*). We find that 32% of informative positions covered by deaminated fragments carry alleles seen in the present-day human, while 5% and 7% carry alleles seen in the Neandertal and Denisovan genomes, respectively. This falls in the range seen for present-day Eurasian individuals (Table S9, Fig. S7) indicating that the Salkhit individual was a modern human, in agreement with more recent morphological analyses (*13*, *14*).

### Salkhit and ancient and present-day modern human

To investigate the relationship of the Salkhit individual to ancient and present-day modern humans, we enriched the libraries for human DNA fragments by hybridization capture using oligonucleotide probes targeting ~2.2 million single-nucleotide polymorphisms (SNPs) selected to be informative about modern human population history (*22*–*24*). Of these, 28% were covered by deaminated DNA fragments in the Salkhit libraries.

We inferred the extent of genetic similarity (using ‘outgroup’ *f3*-*statistics* and *D-statistics* (*25*)) between the Salkhit individual, modern human individuals older than 20,000 years (Table S1) and 131 present-day populations (*19*). The Salkhit individual, similar to the ~40,000-year-old Tianyuan individual from China, is more related to present-day East Eurasians and Native Americans than to West Eurasians (Fig. 1B, Table S10, Fig. S9). Both early East Asians are equally related to most present-day East Eurasians and Native Americans (Fig. 1C, Table S11) but differ in their affinity to West Eurasians. Present-day West Eurasians share more alleles with the Salkhit individual than with the Tianyuan individual (Fig. 1C). Additionally, the Salkhit individual shares as many alleles with the Tianyuan individual as with the ~31,000-year-old Yana individuals from Northeast Siberia (Table S12 and S14), yet the Tianyuan and the Yana individuals share fewer alleles with each other than with the Salkhit individual (Fig. 1C, Tables S12-S14). Those observations suggest that gene flow occurred between populations ancestral to the Salkhit individual and the Yana individuals before ~34,000 years BP, *i.e*. between early populations in East Asia and in Siberia following the divergence of East and West Eurasians. The ~35,000-year-old Goyet Q116 individual from Belgium shares more alleles with the Salkhit individual as well as with the Tianyuan individual (*7*) than do other Europeans analyzed to date (Tables S15-S16). The fact that the Salkhit individual shares even more alleles with the Goyet Q116 individual than does the Tianyuan individual (Fig. 1C) is probably due to gene flow bringing western Eurasian ancestry into the ancestors of the Salkhit individual.

### Admixture model

Population admixture models that are compatible with genomic data from modern human individuals older than 20,000 years were evaluated using qpGraph (*25*) (Fig. 2, Fig. S13). Our models suggest that the Tianyuan individual is an unadmixed representative of an early East Eurasian population and the ancient European individuals *Kostenki 14*, *Sunghir 3* and *Vestonice 16* are representatives of early West Eurasian populations. The Salkhit individual, who lived in Mongolia about 6,000 years after the Tianyuan individual, carries ~75% of its ancestry from the Tianyuan-related East Eurasian population and the remaining ~25% from a Siberian population related to the Yana individuals, who lived some 3,000 years later than the Salkhit individual. In agreement with previous results (*10*), the Yana individuals are estimated to have about a third of their ancestry from early East Eurasians and the remaining two thirds from the early West Eurasians. Their relationship to the Salkhit individual is complex in that models without bidirectional gene flow between an East Asian population (ancestral to the Salkhit and the Tianyuan individuals or only to the Salkhit individual) and a population related to the Yana individuals do not fit the data. Thus, sometime before 34,000 years ago, gene flow from West to East Eurasia occurred, perhaps mediated by ancestors of the early colonizers of Siberia as represented by the Yana individuals.

**Fig. 2.**
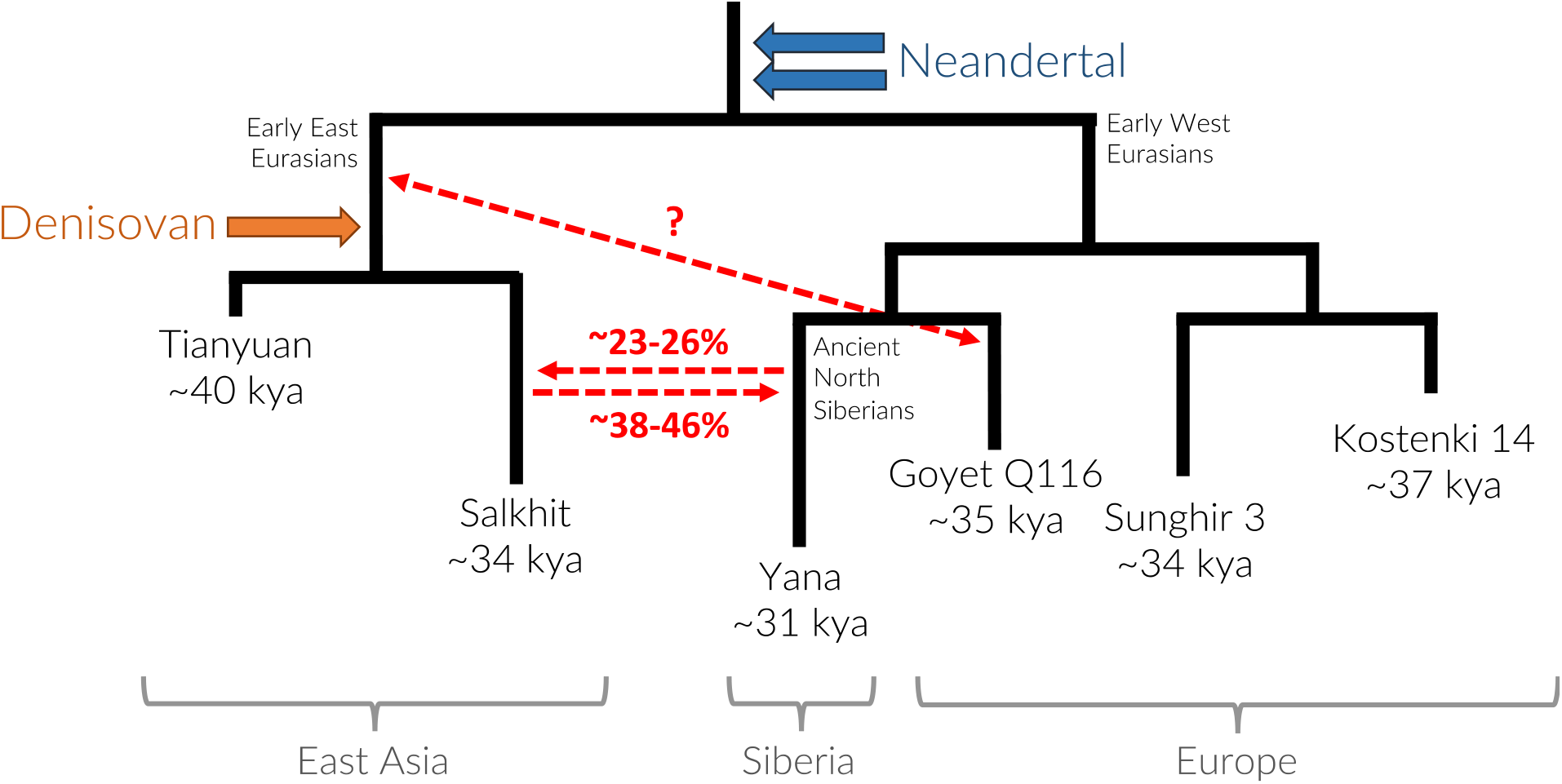
Simplified demographic model including the Salkhit individual and Eurasians older than 20,000 years. The model depicted is the combinations of fit admixture graph models. Admixture between East and West Eurasians are represented by red arrows. The bidirectional gene flow affecting ancestors of the Yana individual may have involved ancestors of the Salkhit individual as indicated or the common ancestors of Tianyuan and Salkhit individuals (not indicated; see Fig. S13). Neandertal and Denisovan admixtures are indicated by blue and orange arrows, respectively.

### Archaic ancestry

We estimate the proportion of Neandertal ancestry in the Salkhit genome to ~1.7% (Table S17, Fig. S14), similar to other early Eurasians. As for all other Eurasian individuals, the Neandertal ancestry in the Salkhit individual is more similar to the Vindija Neandertal from Croatia than to the “Altai” Neandertal from Denisova Cave (*20*, *26*) (Fig.S15).

In addition to the Neandertal ancestry, present-day individuals in East Asia carry ancestry from Denisovans, although in mainland Asia the amount of Denisovan ancestry in present-day populations is more than ten times lower than the amount of Neandertal ancestry (*27*–*29*). This has hitherto made it impossible to determine if ancient genomes from Asia, which are of lower quality than present-day genomes, carry Denisovan ancestry. We applied a novel hidden Markov approach (*30*) that is able to identify introgressed Neandertal and Denisovan genomic segments in low coverage ancient genomes, using a genotype likelihood approach that incorporates contamination, so that all fragments can be used for this analysis. Using data from ~1.7 million SNPs where Neandertal and/or Denisovan genomes differ from present-day African genomes, we detect 18 segments of Denisovan ancestry longer than 0.2 cM in the Salkhit genome (Fig. 3, Table S18, Fig. S16 and S27) and 20 such segments in the Tianyuan genome (Table S18, Fig. S17 and S27). We detect about a third as many segments of Denisovan DNA in the genomes of the ancient Siberians *Yana 1* and *Yana 2*, and *Mal’ta 1* (Table S18, Fig. S18-20 and S27), consistent with that they carry lower proportions of East Asian ancestry. In contrast, no Denisovan ancestry is detected in the genome of the ~45,000-year-old Siberian individual from Ust’Ishim in West Siberia, nor in any European individual older than 20,000 years (Table S18, Fig. S21-24 and S27). Thus, the Salkhit and Tianyuan genomes provide direct evidence that ancestors of modern humans who lived in East Asia 40,000 years ago had met and mixed with Denisovans. The low number of these segments do not provide enough power to date the introgression event. But given their relatively short length (≤ 1.3 cM), the Denisovan introgression is likely to have happened at least ten thousand years before these individuals lived.

**Fig. 3.**
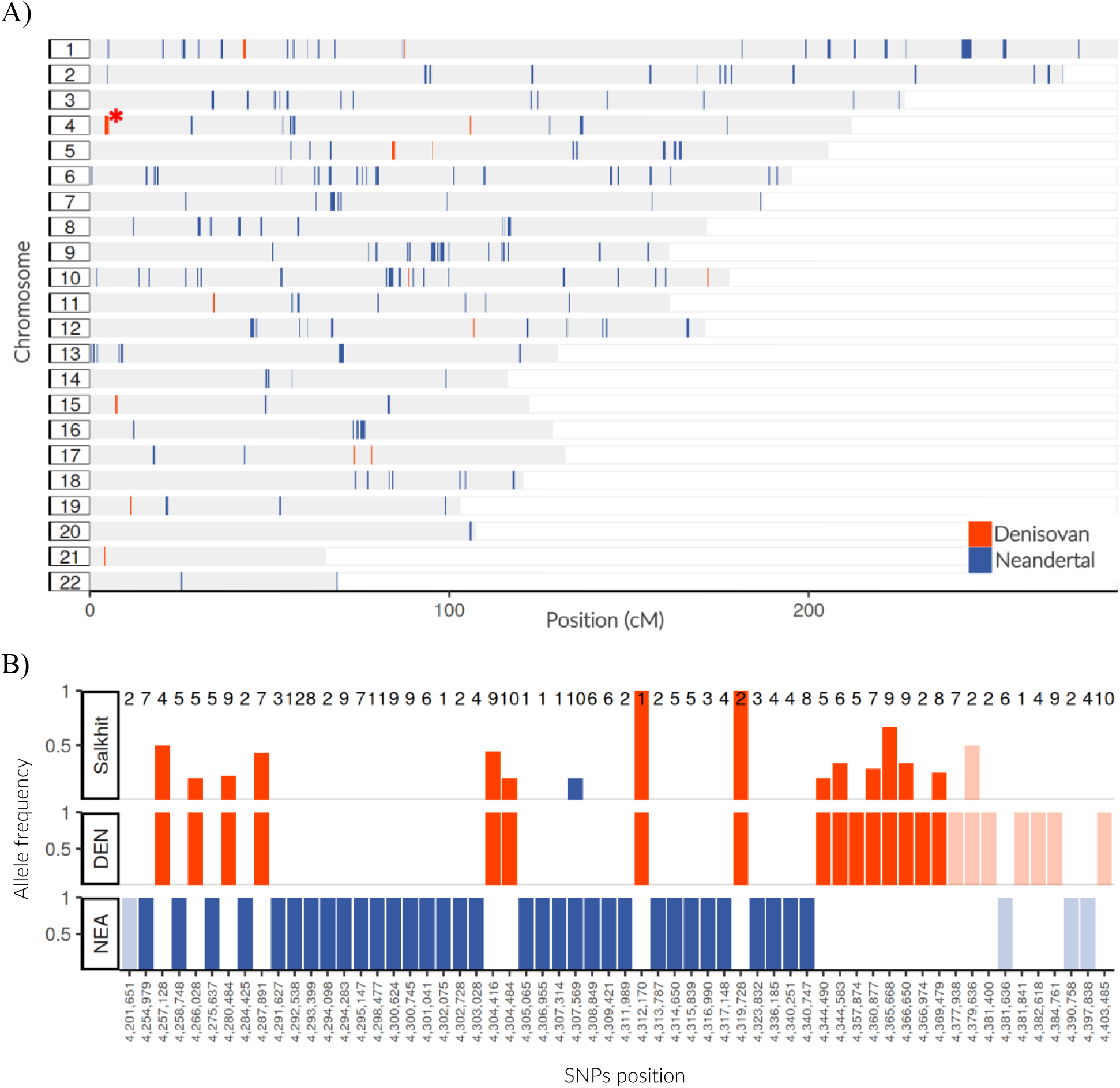
Archaic ancestry in the Salkhit genome. **A)** Genomic distribution of Denisovan (orange) and Neandertal (blue) DNA segments in the Salkhit genome. **B)** Allele frequencies in the longest Denisovan ancestry segment (chr4:4.16-5.31cM, marked by an asterisk in A). The bars in the top panel give the proportions of Salkhit DNA fragments carrying archaic alleles at sites where alleles are fixed between Africans and the Denisovan genome (red), or between Africans and two Neandertal genomes (blue). The total number of fragments are given on top. SNP bars outside the inferred Denisovan segment are faded. Note that in the called region, all Denisovan-like alleles except two occur in the Salkhit genome.

One of the risks of inferring ancestry fragments from ancient genomes is that genome quality may affect the ability to detect introgressed segments. Under the assumption that many of the factors that affect the detection of Denisovan DNA will similarly affect the detection of Neandertal DNA, the ratio of Denisovan to Neandertal ancestry segments detected by AdmixFrog may be a reasonably robust metric of the relative amount of Denisovan ancestry. In the Salkhit and Tianyuan genomes, these ratios are about 7.5% and 8.1%, respectively. For the North Siberians *Yana 1* and *Yana 2* genomes, the ratios are about 3.9% and 4.7%. Because there is no substantial difference in the amounts of Neandertal DNA in the two early East Asian and the two Yana genomes (Fig. S14), this observation indicates that the ancient Siberians individuals carry less Denisovan DNA than the Salkhit and Tianyuan individuals.

### Denisovan ancestry continuity

We compared the Denisovan segments in the Salkhit and Tianyuan genomes to those in present-day people to estimate whether introgressed segments between genomes overlap more often than expected by chance. Significance is assessed using 500 bootstrap reshuffles, where segments are randomly relocated across the analyzed genomes (Supplementary Text 8). The Denisovan DNA segments in the ancient East Asian genomes overlap more than expected with Denisovan segments detected in the genomes of several present-day populations in Asia and in populations with some Asian ancestry such as Hawaiians (Fig. 4, Table S19, Fig. S28-30). In contrast, we find no significant overlap with Denisovan segments detected in Papuans or Aboriginal Australians, although these groups carry on the order of twenty times more Denisovan DNA than mainland Asians.

**Fig.4.**
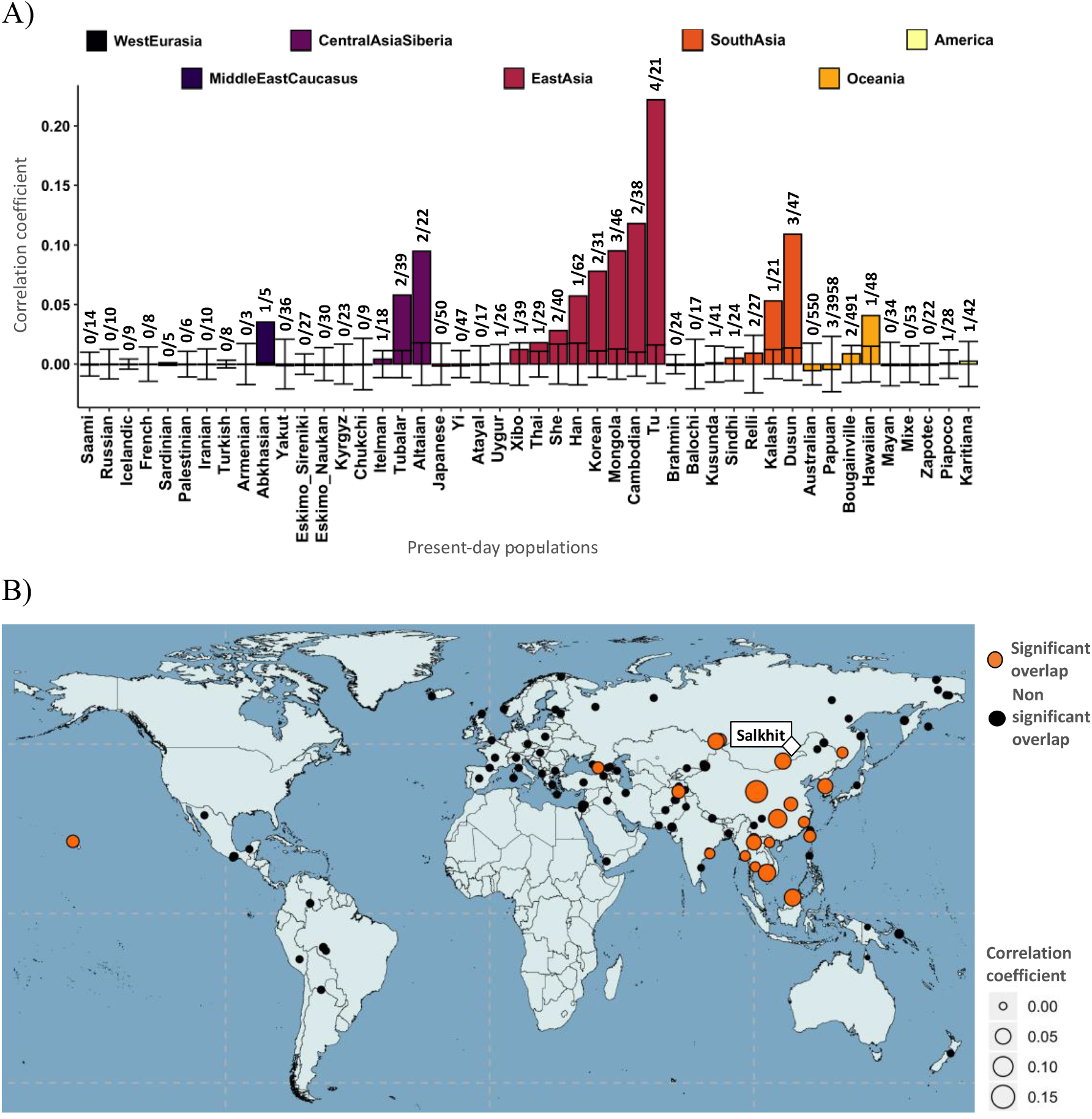
Overlap of Denisovan segments in the Salkhit genome and present-day non-African populations. A) Correlation coefficient of the overlap between the Denisovan segments > 0.2 cM in the Salkhit genome and Denisovan segments > 0.05 cM in 45 present-day Eurasian population (see Fig. S28 for the same with 111 present-day populations). Numbers above the bars give the number of overlapping segments and the number of segments in the present-day population. The range of correlation coefficients generated by 500 bootstraps are indicated. B) Geographic locations of present-day populations for which Denisovan ancestry segments overlap significantly with the Salkhit individual (orange circles). The size of circles is proportional to the correlation coefficients.

It has been shown that at least two Denisovan populations contributed ancestry to present-day East Asian populations, and that Denisovan ancestry in populations in Oceania derived from only one of these sources (*28*). The overlap of Denisovan DNA segments (Fig. 4, Table S19 Fig. S28-30) is in agreement with this and suggests that the ancestral population of the Tianyuan and the Salkhit individuals that mixed with Denisovans contributed ancestry to populations in large parts of Asia today. The lack of any significant overlap with Aboriginal Australian and Papuans suggests that these Oceanian populations received most of their Denisovan ancestry from a different source.

## Conclusion

In conclusion, we show that the 34,000-year-old Salkhit individual carried more West Eurasian ancestry than the 40,000-year-old Tianyuan individual, indicating that after the major West/East Eurasians split, gene flow from West Eurasia to East Asia occurred prior to 34,000 years ago, perhaps mediated by populations related to the Siberian Yana individuals. We also show that these early East Asians carried segments of Denisovan DNA that come from admixture events that also contributed Denisovan DNA to populations across mainland Asia today but not to Papuans and Aboriginal Australians.

## Materials and Methods

### Specimen sampling

The Salkhit skullcap was discovered in 2006 outside archaeological context during mining operations in the Salkhit Valley of the Norovlin county in the Khentii province, eastern Mongolia (48°16′17.9” N and 112°21′37.9” E). The specimen which consists of a mostly complete frontal as well as two partially complete parietal and nasal bones was brought to the attention of the Mongolian Academy of Sciences (*11*). We sampled the specimen at the Institute of Archaeology, Mongolian Academy of Sciences in Ulaanbaatar, Mongolia. After the removal of approximately 1 mm of surface material using a sterile dentistry drill to reduce present-day human DNA contamination, bone powder was drilled from three adjacent spots (A, B and C) at the internal surface of the skull.

### Data

The present-day data used include 288 individual genomes from 136 populations from the Simons Genome Diversity Panel (SGDP) (*19*). The lineage assignment (Supplementary Material 3) was done using 6 present-day Eurasians reported by Prüfer et al., 2014 (*20*) (as Panel B) and the high coverage archaic genomes from a Neandertal (*26*) and a Denisovan individual (*21*). Ancient modern human data used, and their original sources are presented in Table S1.

### DNA extraction

DNA extracts and libraries were made in a clean room facility dedicated to the analysis of ancient DNA at the Max Planck Institute for Evolutionary Anthropology in Leipzig. DNA was extracted from 33.1 mg, 30.9 mg and 27 mg of drilled powder from each of the 3 sampling spots A, B and C, respectively, using a silica-based protocol optimized for the recovery of short molecules from ancient material (*31*, *32*) and eluted in 50μL of TET buffer. Additionally, 51.9 mg, 53.2 mg and 54.9 mg of drilled powder from each of the sampling spot A, B and C, respectively, were incubated with 1 mL of 0.5% of sodium hypochlorite at room temperature for 15 minutes. After centrifugation and removal of the bleach supernatant, the pellet was washed 3 times with 1 mL of water for 3 min to remove residual bleach (*33*, *34*). DNA were extracted from the bleached treated pellet following the same silica-based protocol as for the non-bleached extracts. Extraction negative controls were carried through each step of DNA extraction for both the bleach treated and the non-treated DNA extracts.

### Library preparation

10μL of each extract were converted into single stranded DNA libraries, developed for highly degraded ancient DNA, as previously described (*35*) using an automated liquid handling platform (*36*). One positive control and one negative control were included with each set of library preparation. No uracil-DNA glycosylase treatment was performed to preserve the C to T substitution patterns to authenticate DNA sequences.

### Nuclear SNPs capture

We applied in solution hybridization capture to enrich our libraries for DNA fragments containing single nucleotide polymorphism (SNPs) of interest (*22–24*). After quantification using a NanoDrop ND-1000 (NanoDrop Technologies) photospectrometer, 1 μg of each of the 6 libraries were hybridized to 3 panels of 52-oligonucleotides probes targeting a total of ~2.2 million SNPs ascertained to investigate modern human population history and a 4th panel of ~1.7 million SNPs to study relationship with Neandertals and Denisovans (*22–24*).

**Panel 1 “390k”:** 394,577 SNPs, about 90% of which are on the Affymetrix Human Origins arrays (*23*, *25*).
**Panel 2 “840k”:** 842,630 SNPs, included the rest of the SNPs on the Affymetrix Human Origins array, all SNPs from the Illumina 610-Quad array, all SNPs on the Affimetrix 50k array (*22–24*).
**Panel 3 “1000k”:** 997,780 SNPs constituting all transversion polymorphisms seen in two Yoruba males from Nigeria and transversion polymorphisms seen in the Altai Neanderthal genome (*22–24*).
**Panel 4 “Archaic”:** 1,749,385 where the West-African Yoruba population carry a high frequency of one allele while at least one archaic individual carries an alternative allele (*22–24*).

### Sequencing, mapping and processing

We generated between 2.31 and 5.18 million shotgun sequencing reads from the 6 libraries and between 32.2 and 68.8 million reads for the 24 enriched libraries (6 libraries × 4 oligonucleotide probe panels) (Table S2) using a paired-end configuration of 2 × 76 bp + 7 cycles for each insert and index read on an Illumina HiSeq2500. Base calling was performed using *Bustard* (Illumina) and sequences that did not exactly match one of the expected index combinations were discarded. Adapter sequences were removed and overlapping paired-end reads were merged using *leeHom* with the parameter « --ancientdna » (*37*). The raw data were mapped to the revised human reference genome sequence *hg19* available from the UCSC genome browser using Burrows-Wheeler Aligner (BWA) (*38*) with parameter « -n 0.01 −o 2 −l 16500 » (*21*). Mapped sequences with identical alignment start and end were removed by collapsing into single sequences by consensus calling using *bam-rmdup* (https://github.com/mpieva/biohazard-tools). Sequences longer than 35 bases with a mapping quality higher than 25 were retained for subsequent analyses. In order to assess the impact of present-day human contamination, sequences showing evidence for cytosine (C) to thymine (T) mismatch at their first or last terminal position were filtered and analyzed. For each library dataset, summary statistics including number of reads generated, endogenous DNA content and complexity, proportion of sequences carrying apparent C to T mismatch at their extremities as well as the contribution of each library to the final coverage were obtained using in house perl scripts as previously described (*24*, *35*) (Table S2). For each library, we retained all sequences covering the targeted SNPs from which we filtered the sequences showing evidence of C to T mismatch at their extremities. The genotyping was made by randomly sampling one sequence for each SNP covered.

## Acknowledgments

We thank Montgomery Slatkin (UC Berkeley, USA), Pontus Skoglund (Crick Institute, London, UK), Qiaomei Fu (IVPP, Chinese Academy of sciences, China), David Reich (Harvard Medical School, Boston, USA) and Bence Viola (University of Toronto, Canada) for helpful discussions.

## Author contributions

D.M., S.N. and B.N. performed the laboratory work. D.M., L.S., M.H., M.M., J.K., B.P. and S.P. generated, analyzed and interpreted the data. D.M, L.S, M.H. and B.P performed the computational work. D.T., B.G., S.Y. carried out morphological analyses of the fossil. T.D, T.H carried out radiocarbon dating. S.Y., D.T., B.G. and J.L. analyzed all archaeological data. D.M. wrote the manuscript with input from all authors.

## Data and materials availability

The sequence data generated for the current study are available in the European Nucleotide Archive (ENA) under accession XXXX. All data needed to evaluate the conclusions in the paper are present in the paper and/or the Supplementary Materials. Additional data related to this paper may be requested from the authors.

## Supplementary Informations

S1 Data quality

S2 Human DNA contamination estimates

S3 Sex determination and hominin group assignment

S4 The Salkhit individual and present-day populations

S5 The Salkhit individual and ancient modern humans

S6 Neandertal ancestry in the Salkhit individual

S7 Tracks of Neandertal and Denisovan ancestry in the Salkhit and other modern human genomes

S8 Continuity of the Denisovan ancestry between early East Asians and present-day populations

Figures S1-S30

Tables S1-S19

