## Supplemental Information for "Denisovan ancestry and population history of early East Asians"

S1 Data quality
S2 Present-day human DNA contamination estimates
S3 Sex determination and hominin group assignment
S4 The Salkhit individual and present-day populations
S5 The Salkhit individual and ancient modern humans
S6 Neandertal ancestry in the Salkhit individual
S7 Tracks of Neandertal and Denisovan ancestry in the Salkhit and other modern human genomes
S8 Continuity of the Denisovan ancestry between early East Asians and present-day populations

Figures S1-S30

Tables S1-S19

### Supplementary Information S1 - Data quality

Using two positions where the Salkhit mitochondrial DNA (mtDNA) genome sequence differs from a panel of 311 present-day human mtDNA genomes, the contamination of each extract A, B and C were previously estimated to 42.6%, 35.2% and 19.6%, respectively (*15*). In order to reduce the rate of exogenous contamination of the samples, we treated additional aliquot of drilled powder from samples A, B and C with 0.5% bleach (*33, 34*) and prepared new extracts and libraries.

From shotgun datasets, the overall proportion of DNA in the libraries A, B and C mapping to the human reference genome *hg19* is 1.5%, 0.6% and 0.8% for non-bleached libraries and 5.6%, 2.0% and 2.3% for bleached libraries, respectively (Table S2). The proportion of human DNA in the libraries increased after bleach treatment by a factor of three on average. We estimated the complexity of the human DNA in the libraries by dividing the number of unique sequences mapped to the human genome with the total number of sequences generated and multiplying that proportion by the total number of molecules in the library before amplification as estimated by quantitative qPCR (*35*). If each library were sequenced to exhaustion, the average coverage of the human genome in extracts A, B and C would be 1.2x, 0.4x and 0.4x for non-bleached libraries, and 0.4x, 0.1x and 0.2x for bleached libraries, respectively (Table S2).

The human DNA fragments in the libraries are of an average length of 60, 55 and 56 for the non-bleached treated libraries and 56, 56 and 55 from bleached libraries from extracts A, B and C, respectively. Contaminating DNA molecules may be expected to be on average longer than ancient DNA molecules. Except for extract A, which is most contaminated based on the mtDNA (*15*), we do not observe a decrease of average size of mapped sequences in the bleached extracts (Table S2).

#### Cytosine deamination

Cytosine (C) to thymine (T) mismatches relative to the reference sequence often result from deamination of C residues which accumulate in ancient DNA molecules over time, particularly towards the ends of molecules (*17*). At the 5' end of the sequences, the C to T mismatch frequencies vary from 22.8% to 38.7% for the non-bleached libraries, and from 36.4% to 40.0% for the bleached libraries. At the 3' end of the sequences, the C to T mismatch frequencies vary from 13.1% to 23.9% for the non-bleached libraries and from 24.4% to 25.7% for the bleached libraries (Table S2 and S3 and Fig. S1). The libraries from extract A, showing the highest mtDNA contamination level, have lower C to T mismatch frequencies (Table S2 and Fig. S1) and they increase the most after bleach treatment (Table S2 and Fig. S1).

The presence of contaminating present-day human DNA molecules in the libraries is supported by the fact that sequences that carry C to T mismatches at one end have higher frequencies of C to T mismatch at the other end (conditional C to T frequency) than when all DNA fragments are considered (Table S3 and Fig. S1 and S2). Under the assumption that all deaminated fragments are of ancient origin and that deamination at the two ends of DNA molecules are independent of each other, C to T mismatch frequencies at one end conditioned by a mismatch at the other end reflect the cytosine deamination frequencies of the ancient DNA in the libraries (*18, 39*). An increase of the conditional C to T frequency in comparison to the overall C to T frequency is observed for all 6 libraries at different scales. For extract A, the increase is lower for the bleached extract than for the non-bleached extract. Estimates of contamination levels are presented in Supplementary Material 2.

**Targeted SNPs coverage in Salkhit datasets**

Out of the ~2.2 million SNPs targeted with the three probe panels, we captured 1,914,361 (85.7%) SNPs from the six libraries, and 884,795 (39.6%) if using only sequences showing evidence of C to T mismatch to the reference genome at their first or last positions (Table S2). Those figures fall in the range of genomic information available for other ancient modern humans when intercept for the same set of SNPs (Table S1). Enrichment with the Panel 4 probes (archaic introgression), yielded 1,318,653 SNPs (75.4%), and 762,476 SNPs (43.6%) when considering sequences with C to T mismatches at their first or last position (Table S1 and S2).

### Supplementary Information S2 - Present-day human DNA contamination estimates

We estimated the level of human contamination of the nuclear DNA in the libraries using two approaches. The first approach is based on the conditional C to T mismatch frequencies and the second by using the Hidden Markov Model based *admixfrog* program (*30*).

#### Conditional C to T mismatch frequencies

We use the C to T mismatch frequency at one end of library DNA fragments showing evidence for C to T mismatches at the opposing end as a proxy for the deamination frequency of the endogenous DNA fragments (*39*). For non-contaminated samples, these "conditional mismatch frequencies" should be similar to the mismatch frequencies of all mapped fragments. A higher frequency for the conditional mismatches compared to the overall frequency indicates modern DNA contamination under the assumption that deamination at the two ends of DNA molecules is independent.

The number of shotgun fragments available to estimate the C to T mismatch is relatively low and thus yield large 95% confidence interval (Table S3). For the 6 libraries enriched by capture with the four panels of probes, we estimate contamination levels by subtracting from 1 the quotient of the overall mismatch frequency over the "conditional" one. By combining the data from all probe panels we estimate the contamination level of the non-bleached libraries from extracts A, B and C to ~50%, ~34% and ~19%, respectively, similar to the mtDNA contamination estimates (~43%, ~35% and ~20% for extract A, B and C, respectively (15)) (Tables S3 and S4). For the bleached libraries, the contamination estimates are ~5%, ~15% and ~20%, respectively (Tables S3 and S4).

#### Admixfrog

We estimated the nuclear contamination by present-day human DNA of our libraries using the program Admixfrog (*30*) and obtain contamination level of ~47%, ~25% and ~15% for non-bleached libraries, and ~2%, ~20% and ~23% for bleached libraries A, B and C, respectively (Table S4). Similarly we estimated the contamination in the proportion of fragment showing evidence for C to T mismatch at their first and last base to ~3%, ~3% and ~2% for non-bleached libraries, and ~1%, ~1% and ~2% for bleached libraries A, B and C, respectively (Table S4).

Sets of Salkhit data to investigate contamination effect on population genetic analyses

To investigate how much different contamination levels by present-day human DNA might affect population genetic tests, we organized the data obtained from all Salkhit libraries enriched using Panels 1, 2 and 3 in 6 different sets with different complexity and levels of contamination (Table S5):

- **Salkhit_all:** all sequences from bleached and non-bleached libraries.
- **Salkhit_all_deam:** sequences with terminal C to T mismatch from Salkhit_all.
- **Salkhit_bl_all:** all sequences from the bleached libraries only.
- **Salkhit_bl_all_deam:** sequences with terminal C to T mismatch from Salkhit_bl_all.
- **Salkhit_bl_A:** all sequences from the bleached libraries from sample A.
- **Salkhit_bl_A_deam:** sequences with terminal C to T mismatches from Salkhit_bl_A.

**Present-day human population affinities to different Salkhit datasets**

Using the ADMIXTOOLS package 5.1 (*25*) we computed *D-statistic* tests of the form *D(Salkhit_setX, Salkhit_setY; present-day population, Outgroup)* to compare the relative number of shared alleles between present-day populations and different sets of Salkhit data. The "**Salkhit_all**" and "**Salkhit_all_deam**" sets share similar number of alleles with most present-day human populations around the world ($-2 < Z < 2$ ) with the exception of Western Eurasians populations and some Siberians populations, indicating that the source of present-day human DNA contamination is of European origin (Table S6, Fig. S3). Other combinations of data sets do not show significant differences in allele sharing to the different present-day populations tested (Table S6, Fig. S3).

**Present-day European component in different Salkhit datasets**

Using the ADMIXTOOLS package 5.1(*25*), we computed *f4-ratio* statistic shown in the equation below (*22, 27*) to estimate the proportion of present-day European content in the different sets of Salkhit data using Tianyuan as an outgroup for the excess of Europeans component in the Salkhit genome. The test allows to estimate the proportion of shared alleles between Salkhit and present-day Europeans (Population1 and Population2), represented here by French and Spanish populations, at the exclusion of the Tianyuan individual (Population3).

$$f4-ratio = \frac{f4(French, Mbuti; Salkhit\_set, Tianyuan)}{f4(French, Mbuti; Spanish, Tianyuan)}$$

The set of Salkhit data composed by all sequences has significantly more present-day European component than all the other sets with an estimate of ~32% (Table S7 and Fig. S4). All the other Salkhit datasets are overlapping in their estimates of present-day European component (Table S7 and Fig. S4) with slight variations due to the difference in contamination level or complexity. Despite having less complexity both sets **Salkhit_all_deam** and **Salkhit_bl_all_deam** give significant results unlike the **Salkhit_bl_A_deam** composed of deaminated fragment from one library only.

**Salkhit Dataset for population genetics analyses**

The complexity of both less contaminated sets, **Salkhit_all_deam** and **Salkhit_bl_deam**, composed by deaminated sequences provide enough power for further population genetic analyses. We investigated the genetic similarity of each set to present-day populations using outgroup *f3-statistic* of the form *f3(Salkhit,X;Mbuti)* when X is a present-day population. The plot presented in Figure S5 shows the correlation of genetic similarity between both sets and the different present-day populations. If both sets were totally equivalent, all the dots would be on the diagonal line. However, we observe a slight tendency of increase of affinity from the present-day populations toward the **Salkhit_all_deam** set (Fig. S5) which has more complexity than **Salkhit_bl_all_deam**. The difference in genetic affinity observed is the reflect of the difference of complexity between both sets. To infer the population history of the Salkhit individual we are using the set of deaminated fragment with most complexity "**Salkhit_deam**".

### Supplementary Information S3 – Sex determination and hominin group assignment

#### Sex determination

The prominent supraorbital region has been invoked to suggest that the Salkhit individual is a male (*13*). We determined the biological sex of the specimen by comparing the coverage of the X chromosome to the coverage of the autosomal chromosomes using 218,000 DNA fragments longer than 34 bases generated by shotgun sequencing of the 6 Salkhit libraries, mapped uniquely to the human genome *hg19*. They cover 0.39% of the X chromosome and between 0.37% and 0.43% of each of the 22 autosomes (Table S8 and Fig. S6). Using 48,900 sequences with C to T mismatch to the reference at their first or last base, we cover 0.049% of the X chromosome while the autosomes coverage ranges from 0.049% to 0.061 (Table S8 and Fig. S6). Thus, the coverage of the X chromosome falls in the range of the coverage of the autosomes, indicating that the Salkhit individual was a female.

#### Hominin group assignment

To determine whether the Salkhit individual is more related to modern humans, Neandertals or Denisovans, we used the high quality genome sequence of the Altai Neandertal (*20*), the Denisovan (*21*), and a present-day human individual from Africa (Mbuti, HGDP00982) (*20*) to identify derived states at positions where one or more of these three genomes differ from those of the chimpanzee and other primates (bonobo, gorilla, orangutan, rhesus macaque) (*39*). We then estimate the percentages of those informative positions covered in the Salkhit individual that share the derived state for each branch in the tree relating the three genome sequences (Table S9, Fig. S7). For comparison, we estimate the percentage of derived allele sharing for 6 non-African individuals (Dai-HGDP01308, French-HGDP00533, Han-HGDP00775, Papuan-HGDP00546, Sardinian-HGDP01076, Karitiana-HGDP01015) (Table S9) (*20*) using the same approach. We find that 33.1% of all Salkhit shotgun sequences covering the informative positions, and 32.4% of sequences showing C to T mismatch at their first or last bases, carry the modern human derived alleles. For the 6 present-day non-African individuals, that proportions range from 30.1% to 32.6% (Table S9, Fig. S7). The proportion of Neandertal derived alleles are 4.3% and 5.4% for all Salkhit sequences and for sequences showing evidence for C to T mismatch, respectively, while for the 6 present-day Eurasian individuals it ranges from 4.1% to 4.5% (Table S9, fig. S7). The proportion of shared Denisovan derived alleles are 8% and 6.9% for all Salkhit sequences and for sequences showing evidence for C to T mismatch, respectively, while for the 6 present-day Eurasian individuals it ranges from 6.5% to 8.3% (Table S9). Overall, the results of the Salkhit dataset overlap with these of the 6 present-day Eurasian individuals, which indicate that the Salkhit individual was a modern human.

The advantage of this test is that it works well on limited amount of data. Because it is based on diagnostic position identified on one single individual for each hominin group, it only shows globally whether one individual is more related to one or another hominin group and cannot be used to infer further relationship between the individual tested and the different hominin groups used.

### Supplementary Information S4 – The Salkhit individual and present-day populations

#### Principal component analysis

To get an overview of the genetic relationships of the Salkhit and other ancient individuals in the context of present-day human genetic variation, we performed a principal component analysis (PCA) using 105 non-African present-day populations (211 individuals) from the Simon Genome Diversity Project (SGDP) (*19*) to compute the principal components and projected 15 modern humans older than 20,000 years (Table S1, Fig. S8). The Salkhit dataset used is composed of deaminated fragments only from the libraries enriched with probe panels 1, 2 and 3. Figure S8a and S8b presents the genetic clustering of the dataset by plotting each individual according to the first and second principal component PC1 and PC2 which explain respectively 5.3% and 3.2% and to the first and third component PC1 and PC3 which explain respectively 5.3% and 2.5% of our dataset variability. The Oase (*22*) and Ust'Ishim (*8*) individuals, who diverged from the main Eurasian lineage before the major West/East split, are around the center of the plot (coordinate ~0,~0), implying equal genetic distance to the populations used to compute the principal components. All other ancient West Eurasians individuals cluster closer to present-day Europeans than to other present-day populations. The North Eastern Siberian Yana individuals (*10*) are also closer to Europeans than to East Asians along PC1. The Salkhit and Tianyuan individuals are closer to present-day Asian populations, but they do not cluster together, suggesting that they have different affinities to some of the present-day populations used in the analyses.

#### Genetic similarity between ancient Eurasians and present-day populations

We estimated genetic similarity between different early modern human individuals including the Salkhit and present-day human populations using Outgroup *f3*-statistics (*25*) of the form *f3(Y,X;Mbuti),* where Y is an ancient individual and X is a present-day Eurasian population. The test shows that the Salkhit individual shares more genetic similarity with present-day East Asians and Native Americans than with West Eurasians (Table S10 and Fig. S9). In agreement with published results, *Kostenki14, Sunghir3* and *Goyet-Q116* share more genetic similarity with present-day West Eurasians than with East Eurasians (Fig. S10) (*24, 40, 41*). The Siberian individuals *Yana1* and *Malta 1* share the most genetic similarity with Native Americans and with West Eurasians than with East Eurasians (*9, 10*). And, as the Salkhit individual, the Tianyuan individual from China shares the most genetic affinity with present-day East-Eurasians and Native Americans (7). However, the Salkhit individual shares more drift with present-day Western Eurasians than does the Tianyuan individual (p=0.0008) (Fig. S11).

The difference in genetic affinity between both early East Asians and present-day Europeans populations computed using *f3-statistic* are consistent with the symmetry test *D-statistic* of the form *D(Salkhit, Tianyuan; X, Mbuti)*, in that when X is a present-day European population, the *D*-statistic is always significantly positive, showing that the Salkhit individual share more alleles with present-day West Eurasians than does the Tianyuan individual (Table S11).

### Supplementary Information S5 – The Salkhit individual and ancient modern humans

#### Salkhit individual is equally related to the 40,000-year-old Tianyuan individual from China and to the 31,000-year-old Yana individuals from North East Siberia.

We investigated the relationship of the Salkhit individual to other modern human older than 20,000 years by comparing their genetic similarity using outgroup *f3-statistic* and *D-statistic*. *D(Salkhit, X, Tianyuan, Mbuti)* is always significantly positive when X is a modern human showing that the 40,000-year old Tianyuan individual from China is genetically closer to the Salkhit individual than to any other modern human (Table S12 and 13 and Fig. S12). However, *D(Tianyuan, X, Salkhit, Mbuti)* is significantly positive when X is a modern human with the exception of the two Yana individuals, indicating that the Salkhit individual share an equivalent amount of alleles with the Tianyuan and the Yana individuals (Table S14).

#### Ancient Europeans are more related to the Salkhit individual than to the Tianyuan individual

*f3(X, Y, Mbuti),* where X and Y are modern humans older than 20,000 years (Fig. S12), shows that the Salkhit individual is more genetically similar to ancient modern human older than 20,000 years analyzed to date than is the Tianyuan individual (Table S12, Fig. S12), including the 35,000-year-old *Goyet-Q116* from Belgium, previously shown to have an unexpected connection to the Tianyuan individual (*7*). Both Salkhit and Tianyuan share more genetic similarity with *Goyet-Q116* than they share with other ancient Europeans older than 20,000 years analyzed to date (Fig. S12). *D(Goyet-Q116, X; Salkhit, Mbuti)* and *D(Goyet-Q116, X; Tianyuan, Mbuti)* are always significantly positive (Z>2) when X is an ancient Europeans older than 20,000 years (Tables S15-16). However, *D(Salkhit, Tianyuan, Goyet-Q116, Mbuti)* is positive (Z=2.99) (Table S11) showing that as other Europeans, *Goyet-Q116* shares more alleles with the Salkhit individual than with the Tianyuan individual.

The tests of genetic similarity show that the Salkhit individual has an excess of allele sharing with Europeans in comparison to the Tianyuan individual. The Salkhit individual shares an equivalent amount of allele with the Tianyuan and the Yana individuals. Considering that the Yana individuals have affinities to both West Europeans and East Asians (*10*), the genetic affinity of the Salkhit individual to Europeans may be the consequence of gene flow between the Salkhit and Yana ancestors. Thus, while the 40,000-year-old Tianyuan individual seems be a relatively unadmixed instantiation of early East Asians, the 34,000-year-old Salkhit individual contain an additional early West Eurasian genetic component.

**Admixture graph modeling**

We used Admixture Graph model (qpGraph) (*25*) from ADMIXTOOLS package 5.1 with default parameters to structure the allele sharing between the 40,000-year-old Tianyuan individual, the 37,000-year-old Kostenki14 individual, the 35,000-year-old Goyet-Q116 individual, the 34,000-year-old Salkhit individual, the 34,000-year-old Sunghir3 individual, the 31,000-year-old Yana individuals and the 30,000-year-old Vestonice16 individual. The program evaluated whether each tested model is a fit to the data by testing whether the predicted values of all the *f2-, f3-, f4-statistic* among all possible pairs, triples and quadruples of individuals matched the observed values, and assessing the significance of the difference using an empirical standard error computed with a Block Jackknife (*23-25*). For each model, the program estimates also the admixture proportions.

Models which fit the data with -3 < Zscore < 3 show early split between East and West Eurasian with the Tianyuan individual as an unadmixed representative of early East Eurasians and Kostenki14, Goyet-Q116, Sunghir3 and Vestonice16 as representative of early West Eurasian. The West Eurasian component in the Salkhit individual and the East Eurasian component in the North-East Siberian Yana individuals are explained by bidirectional gene flow between early East Eurasians (either ancestor of the Salkhit after the split from Tianyuan (Fig. S13A, B, C and D) or common ancestor of both the Salkhit and Tianyuan (Fig. S13E, F, G and H) and Yana ancestors (Fig. S13). That bidirectional gene flow can be modeled either sequentially with an admixture event between West and East Eurasians leading to an intermediate admixed population which contributed ancestry to the Salkhit and Yana individuals (Fig. S13A, B, E anf F), or in parallel with gene flows from the ancestral population of the Yana individuals into the Salkhit ancestors and vice versa (Fig. S13C, D, G anf H). The Salkhit individual is estimated to have around quarter of ancestry from a West Eurasian population related to the Yana individuals (Fig. S13).

Additionaly, the connection between the *Goyet-Q116* and the Tianyuan and the Salkhit individuals is as well supported when the fitted models involve gene flow from East Asians to a population ancestral to *Goyet-Q116* (Fig. S13A, C, E and G) or in the opposite direction (Fig.

S13B, D, F and H). Additional genomes from early West ad East Eurasians are needed to resolve the relationship of *Goyet-Q116* to early East Asians.

**Supplementary Information S6 – Neandertal ancestry in the Salkhit individual**

To estimate the proportion of Neandertal ancestry in the Salkhit genome and in other early and present-day modern human, we used *f4-ratio* statistic introduced by Reich et al., 2010 (*27*) making use of making use of the two high quality Neandertal genomes available (*20, 26*) as previously described by Petr et al., 2019 (*42*). We used ADMIXTOOLS package (*25*) version 5.1 to process the statistic according to the following equation:

$$f4-ratio = \frac{f4(Alta\ddot{i}\ neandertal, Chimp; X, Mbuti)}{f4(Alta\ddot{i}\ neandertal, Chimp; Vindija\ neandertal, Mbuti)}$$

where X is a modern human. We estimated the proportion of Neandertal ancestry in the Salkhit genome to 1.7% (1.2% - 2.2% CI95%), similar to most early modern humans available, including the Tianyuan individual 1.7% (1.3-2.1 CI95%) (Table S17, Fig. S14), and with the sole exception of *Oase 1* (*22*).

*D*-statistic of the form *D(Altaï Neandertal, Vindija Neandertal; Salkhit, Mbuti)* shows that the Salkhit individual shared significantly more alleles with the Vindija Neandertal than with the Altaï Neandertal, as is the case for other modern humans (*34*) (Fig. S15)

**Supplementary Information S7 – Tracks of Neandertal and Denisovan ancestry in the Salkhit and other modern human genomes**

In addition to the Neandertal ancestry, present-day individuals in East and Southeast Asia as well as in Oceania carry ancestry from Denisovans (*20, 27*). In present-day East Asians, the proportion of Denisovan ancestry is estimated to ~0.2% (*28, 43*). However, due to insufficient power of the current methods to detect very low archaic component in low coverage genomes, it has not been possible to detect Denisovan ancestry signal in ancient East Asians such as the Tianyuan individual (*7*). Using the *admixfrog* program, a novel Hidden Markov Model developed for local ancestry inference (*30*) we investigated the Denisova and Neandertal components in the Salkhit genome and how these are related to other ancient and present-day individuals.

**Identifying archaic introgressed tracts**

We detect archaic introgressed tracts from low-coverage, contaminated capture data using the *admixfrog* method (*30*). The method is run on 1,749,385 sites from the "archaic admixture array" (Panel 4) (*22*), which are designed to be informative on archaic admixture in modern human. We restrict the analysis to 1,349,087 sites that are biallelic in a combined data set of SGDP and 1000G.

First we create a reference file that contains allele frequencies form the following populations:

- Africans (AFR): 44 Africans from SGDP (*19*)
- Neanderthal (NEA): The Altai (*Denisova5*) (*20*) and Vindija Neandertal (*26*)
- Denisovan (DEN): *Denisova3* (*21*)
- Primates: Chimpanzee (*pantro4*)

by using the commmand:

admixfrog-ref --out stats/admixfrog_panels/new/ref_archaicadmixture.csv.xz --state-file config/data.yaml --rec-file recs/maps_b37/maps_chr.{CHROM} --vcf-ref vcfs/vcf_other/chrom_asc_giant/archaicadmixture/{CHROM}.vcf.gz --force-ref --states NEA=AltaiNeandertal,Vindija33.19 DEN=Denisova ALT=AltaiNeandertal CHA=Chagyrskaya-Phalanx VIN=Vindija33.19 PAN=panTro4 AFR=sgdpafr.

All analyses use the African American genetic map (44) from https://www.well.ox.ac.uk/~anjali/Aamap/, (build hg19), linearly interpolated, when necessary.

For each ancient individual we prepare the infiles from bam-files using the commands:

admixfrog-bam --bamfile {bamfile}  --out {name}.in.xz –ref stats/admixfrog_panels/new/ref_archaicadmixture.csv.xz

where name is the name of the individual in the analysis, *e.g*. Salkhit and bamfile is the path to the bamfile.

To identify archaic introgressed regions in the genome we run the command:

admixfrog --ref {reffile} --infile {name}.in.xz --out {outputprefix} --female -b {winsize} –states AFR NEA DEN –cont AFR –ancestral PAN

This command runs admixfrog with AFR NEA and DEN as potential sources, using AFR as a potential contaminant source, and the chimpanzee genotype as putative ancestral allele. We set the window size to 0.005 cM.

**Samples used**

We ran *admixfrog* on 11 ancient individuals (where coverage was above 1X) and 235 non-African present-day individuals from the Simons Diversity Project (*19*). Figure S16 to S24 show the distribution of Neandertal and Denisovan tracks called with *admixfrog* from the genomes of the early modern humans older than 20,000 years Tianyuan, Salkhit, *Yana1, Yana2, Malta, Ust'Ishim, Oase, Kostenki14* and *Sunghir3*.

As short fragments are difficult to identify particularly in low-coverage genomes, we only consider fragments that are longer than 0.2 cM for ancient samples, and longer than 0.05 cM for present-day samples, as above these cutoffs the power and precision of *admixfrog* is very high. We called 20 and 18 fragments of Denisovan ancestry in the Tianyuan and the Salkhit genomes, respectively, and 3, 6 and 4 such fragments in the genomes of *Yana1*, *Yana2* and *Malta1* individuals. Figure S25 and S26 show the allele frequency of the largest Denisovan introgressed fragment in the Salkhit and the Tianyuan genomes, respectively.

A list of all ancient samples analyzed and the number of Denisovan ancestry tracts > 0.2 cM in their genome is shown in table S18 as well as in Figure S26.

**Supplementary Information S8 – Continuity of the Denisovan ancestry between early East Asians and present-day populations**

We wanted to investigate if the Denisovan ancestry fragments detected in the Salkhit and the Tianyuan genomes overlap with those present in present-day humans. If the archaic fragments overlap between two individuals more than one would expect by chance, their genomic location correlate and they are likely to share common ancestry and originate from the same admixture event.

First, we calculate the number of Denisovan fragments that overlap between any pair of present-day and ancient individuals. For each of 109 Simons Diversity populations (*19*), we group the archaic fragments from all individuals to increase our power. To summarize the overlap, we use the Pearson correlation coefficient as a summary statistic. This statistic is equal to 0 if there is no correlation and equates to 1 if all fragments overlap. In order to compute confidence intervals for the amount of overlap we perform 500 bootstrap iterations where we randomly place archaic fragments of the same number and sizes than the observed ones along the genome (with a length of 2.8 Gb to account for uncallable regions) for both individuals (Table S19).

The Denisovan ancestry in both Salkhit and Tianyuan genomes correlate significantly with the Denisovan ancestry in populations in Central and East Asia (Fig. S28 to S30). In Oceania we do not observe a significant overlap between Salkhit and Papuan Denisovan fragments, despite Papuans having the highest amount of Denisova ancestry. This suggests that the Denisovan component in Salkhit is not originating from the same event as the Denisova component in Papuans.

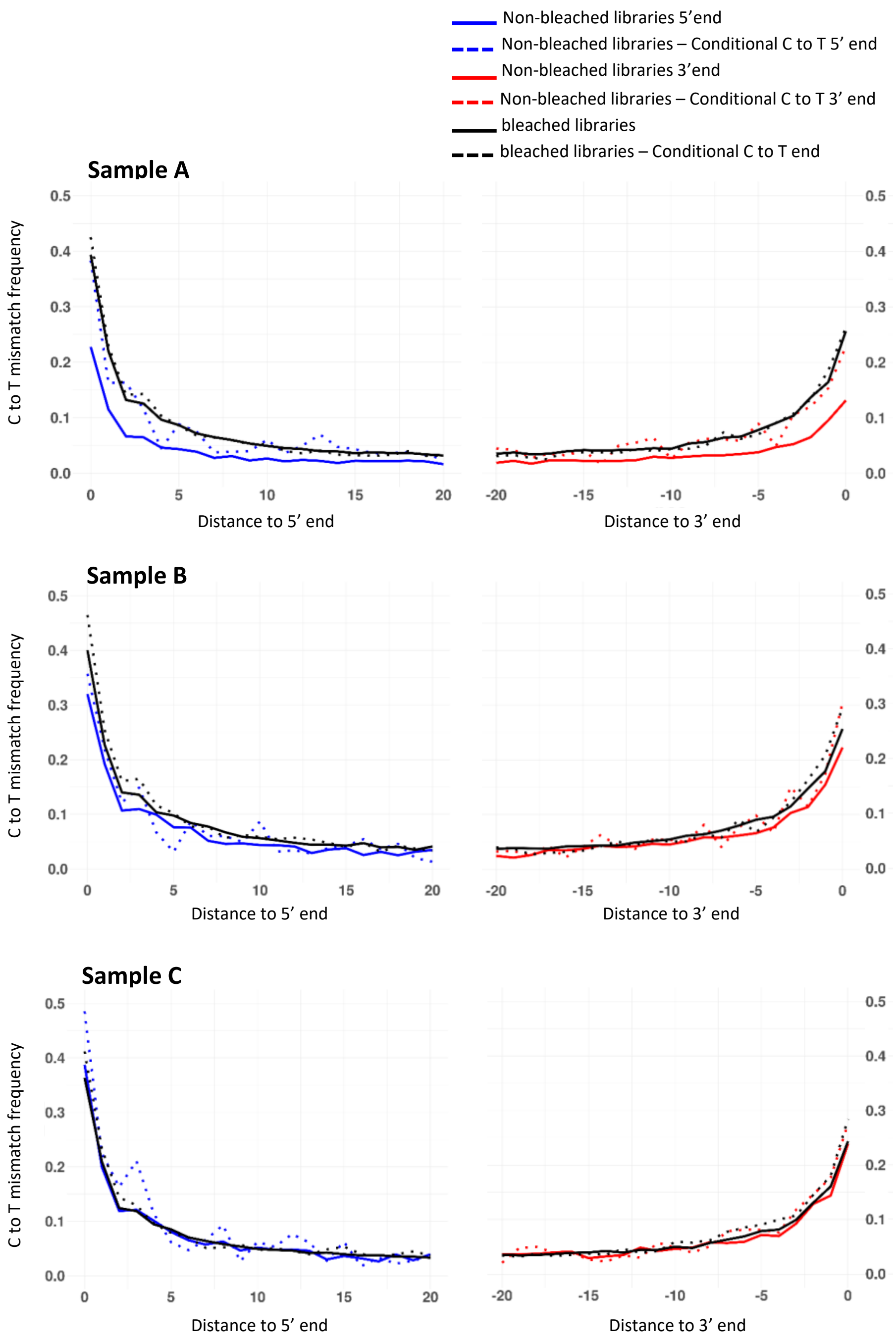


**Fig. S1 C to T mismatch frequencies in Salkhit libraries.** The frequencies of C to T mismatches of the 20 terminal positions of the Salkhit DNA shotgun reads relative to the reference sequence *hg19* are shown for both non-bleached and bleached libraries made from the three samples A, B and C. Conditional C to T mismatches corresponding to the C to T mismatch frequency at one end for sequences carrying a C to T mismatch at the other end, are shown in dotted lines.

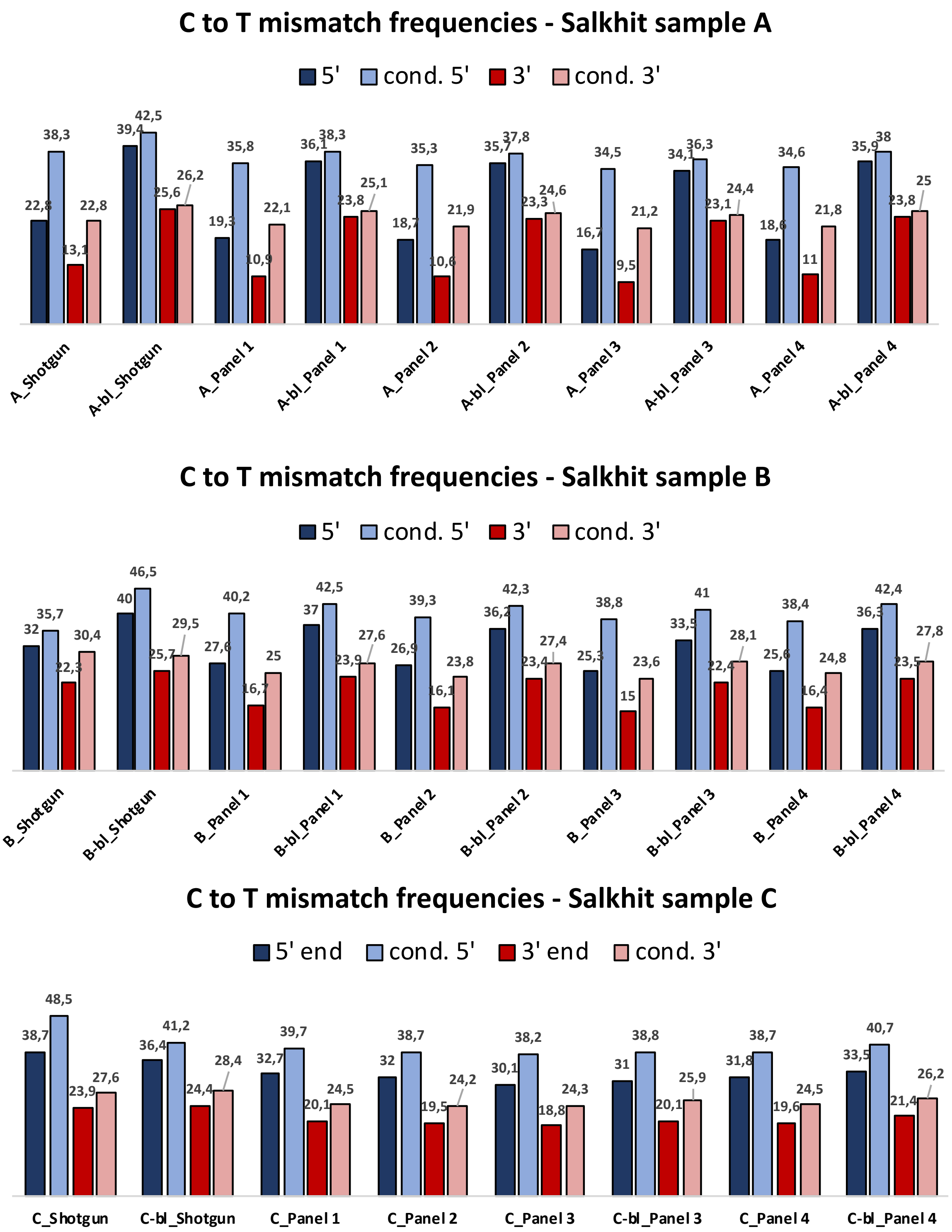


**Fig. S2 C to T mismatch frequencies of Salkhit libraries.** The frequencies of C to T mismatches for the terminal positions at 5'- and 3'-ends relative to the reference sequence *hg19* before and after enrichment with the four SNPs panels are shown for non-bleached and bleached libraries made from samples A, B and C. Conditional C to T mismatches are the C to T mismatches at one end of DNA fragments carrying a C to T mismatch at the other end.

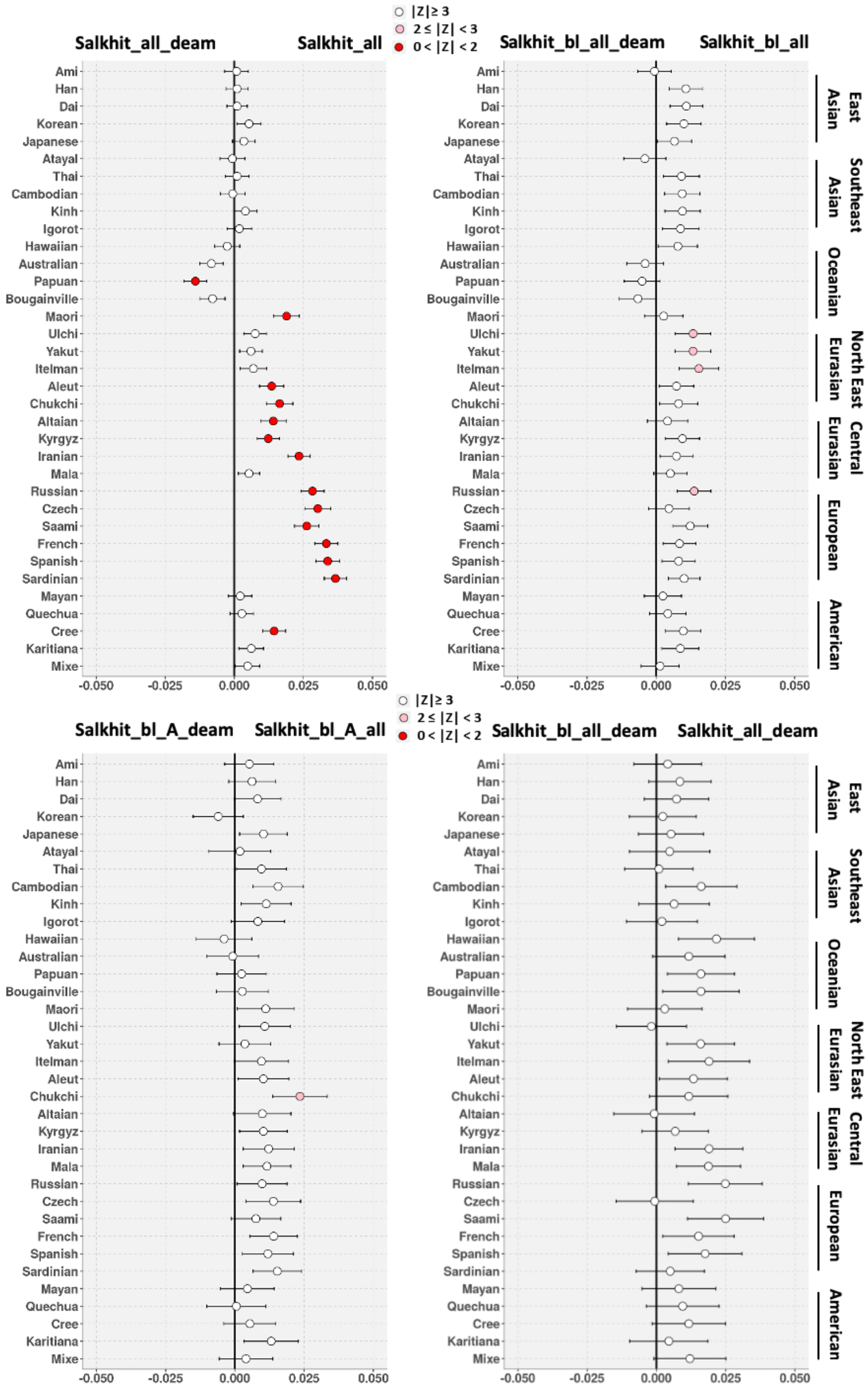


**Fig. S3 Difference of genetic affinity between present-day population and different Salkhit dataset with different level of contamination**. *D-Satistic* of the form *D(Salkhit-set1, Salkhit-set2, present-day_population, Mbuti)* comparing the number of shared alleles between a present-day population and different Salkhit datasets.

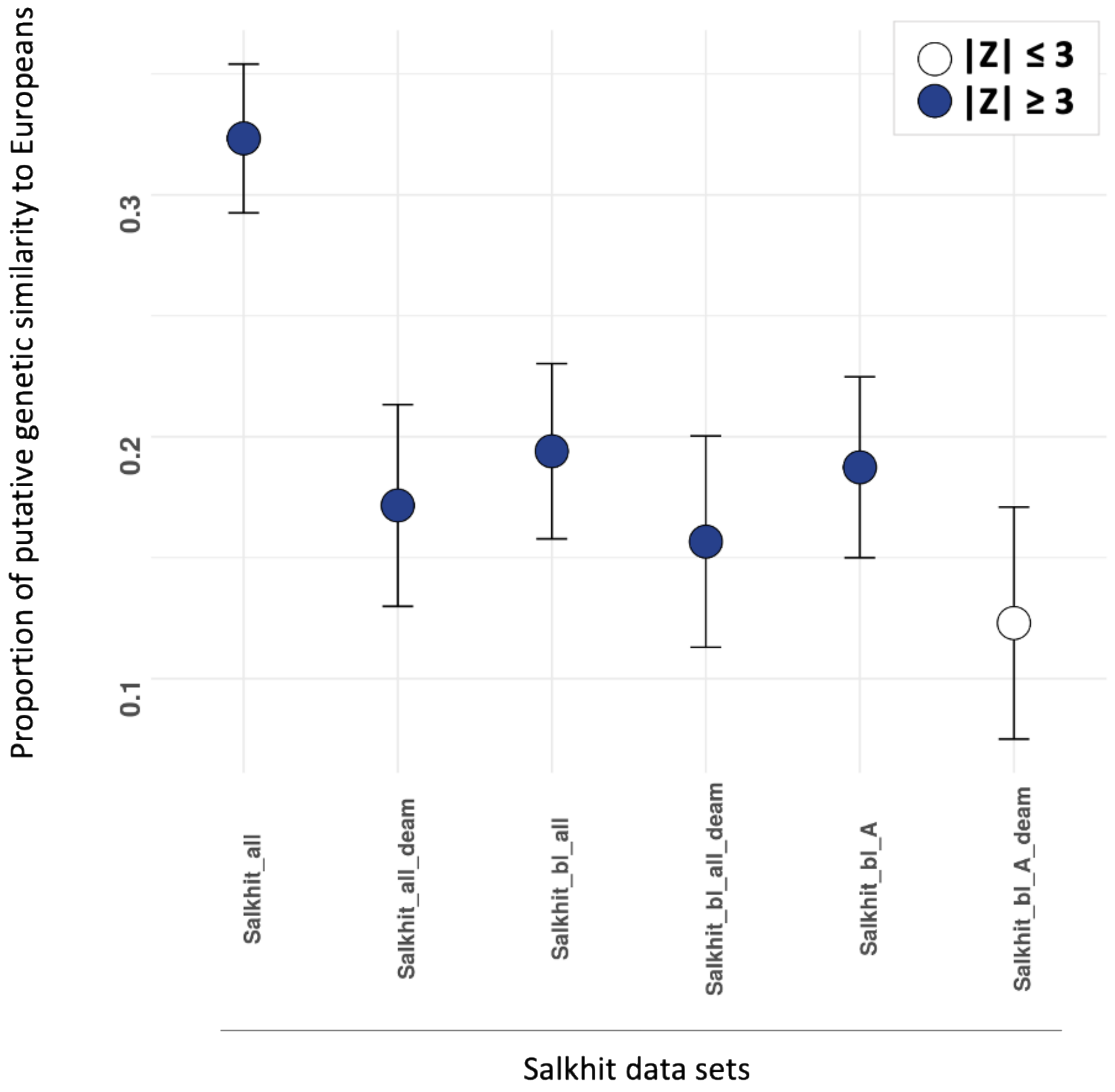


**Fig. S4. Relative proportion of shared allele between present-day European and different Salkhit datasets.** The proportion is estimate by f4-ratio of the form *f4(French, Mbuti; Salkhit_set, Tianyuan/French, Mbuti; Spanish, Tianyuan)*.

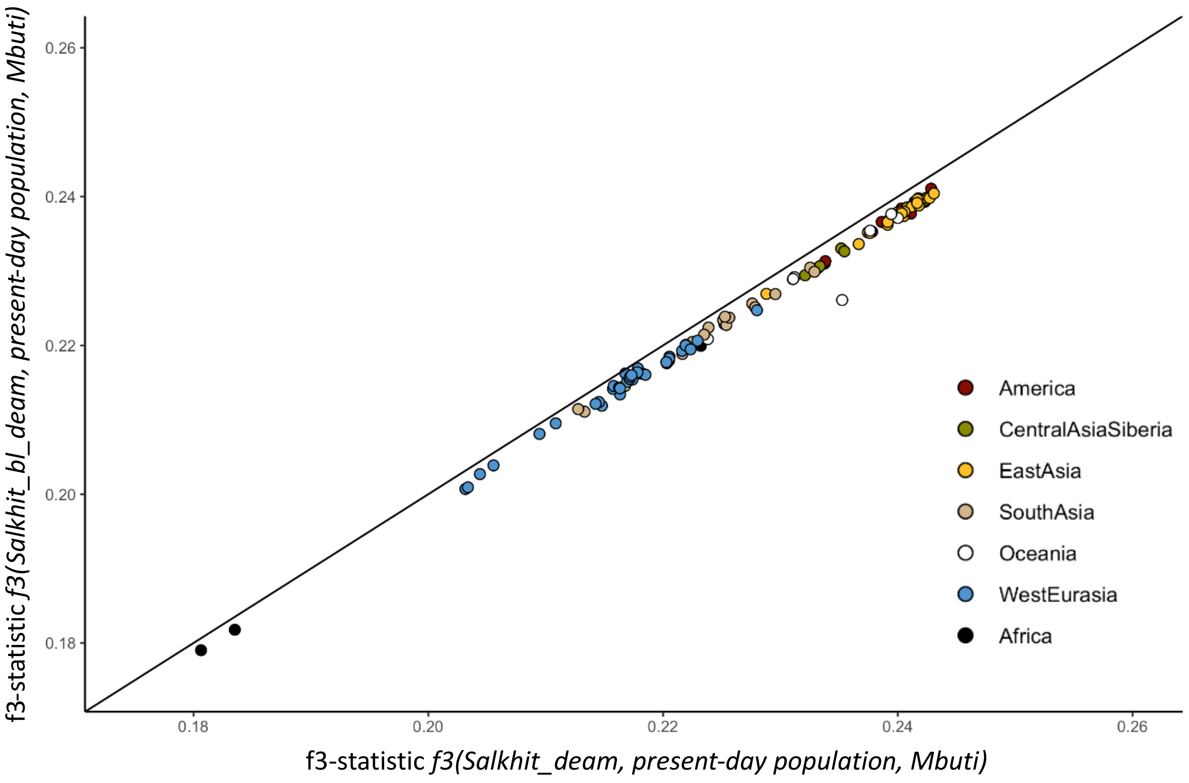


**Fig. S5. Difference of genetic affinity of present-day population to different Salkhit genetic datasets.** Correlation plot of the *f3 statistics* of present-day population to the Salkhit_all_deam dataset composed of deaminated fragments from all libraries and to the Salkhit_bl_all_deam composed of deaminated fragments from bleached libraries of the Salkhit.

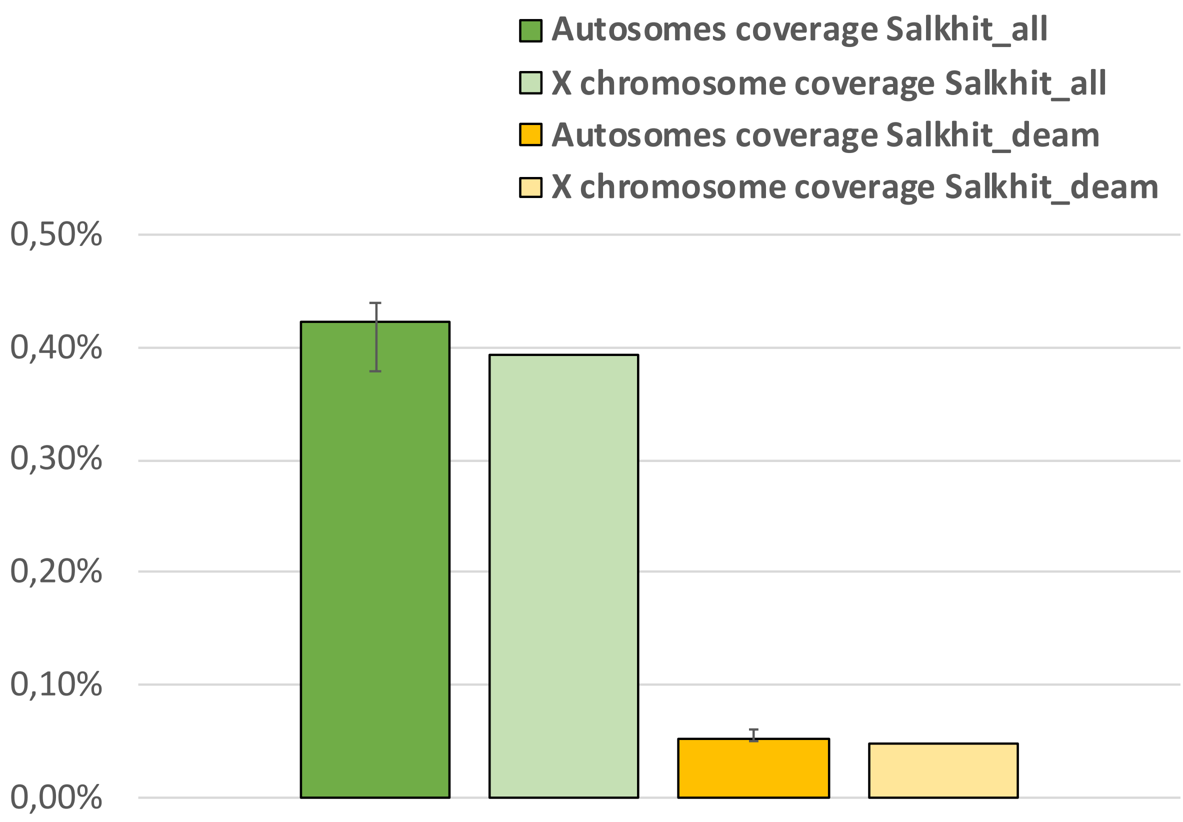


**Fig. S6. Sex determination of the Salkhit individual.** Coverage of the X chromosome versus average coverage of the autosomes in Salkhit shotgun datasets for all fragments (green) and for deaminated fragments (yellow).

A)

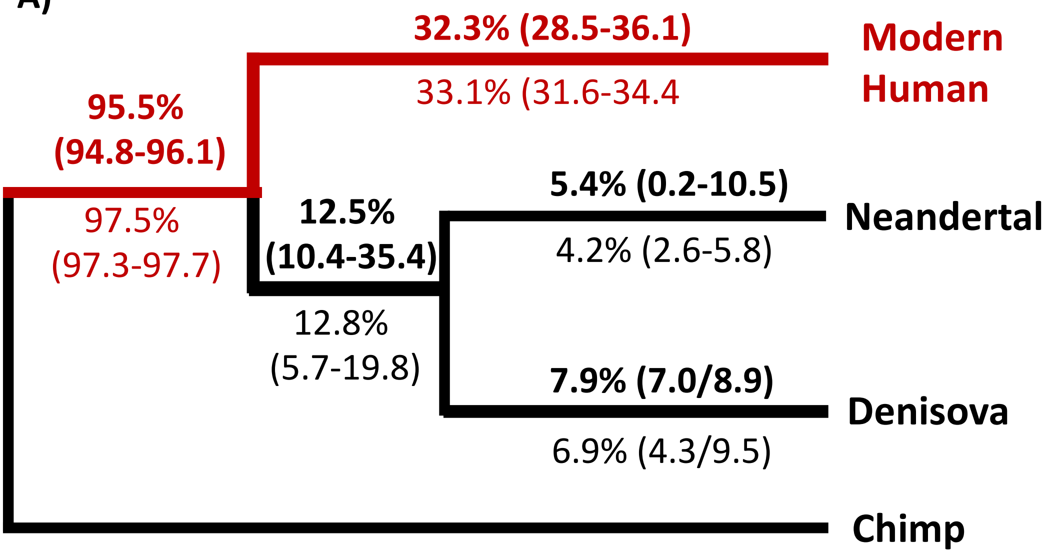


B.

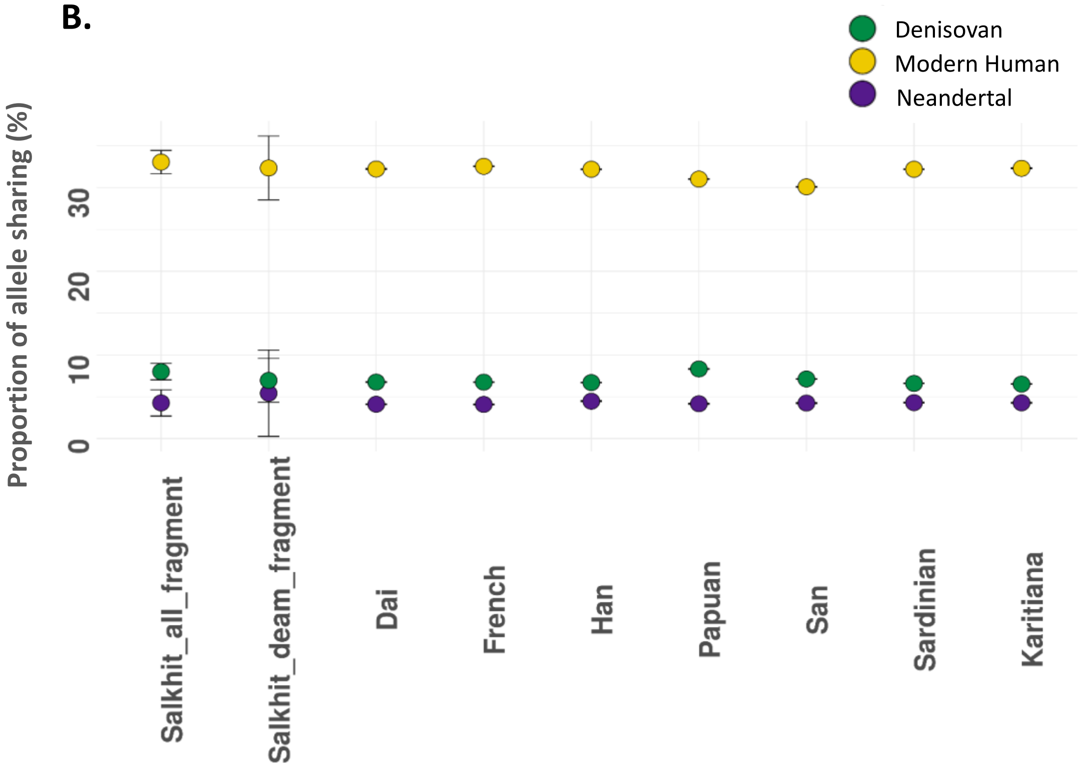


**Fig. S7. Hominin group assignment of the Salkhit individual.** A) Percentage of derived alleles shared between the Salkhit individual and human, Neanderthal and Denisovan genomes. In bold the estimates based on the deaminated fragment, while the other are based on all fragment. 95% binomial confidence interval are in parenthesis. B) Derived allele sharing with modern human, Neanderthal, and Denisovan branches of Salkhit all fragments and deaminated fragments and 7 present-day human individuals.

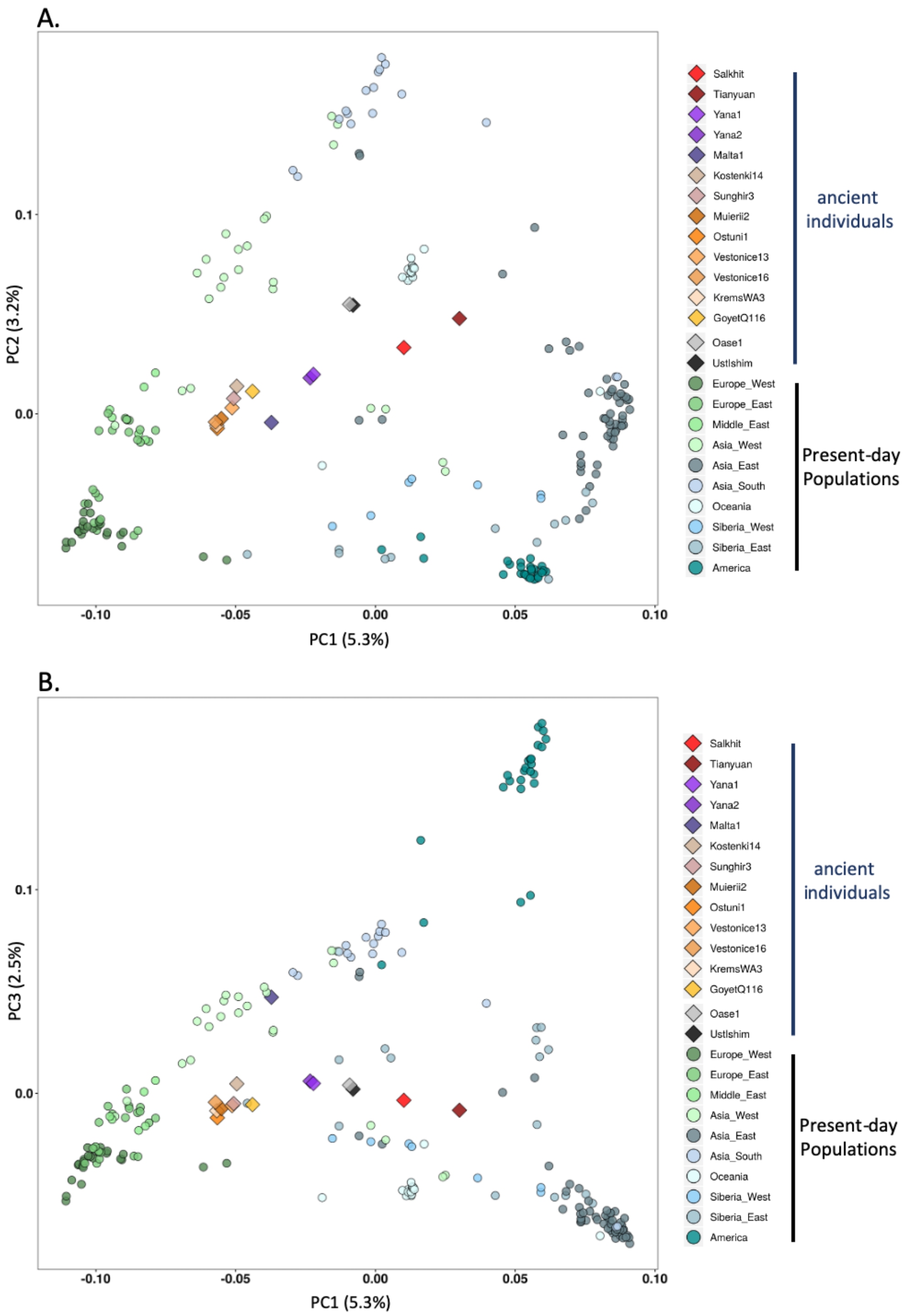


**Fig. S8. Projection of the Salkhit individual and 14 others ancient individuals on a principal component analyses of present-day non-African genetic variation.** A) PC1 and PC2. B) PC1 and PC3.

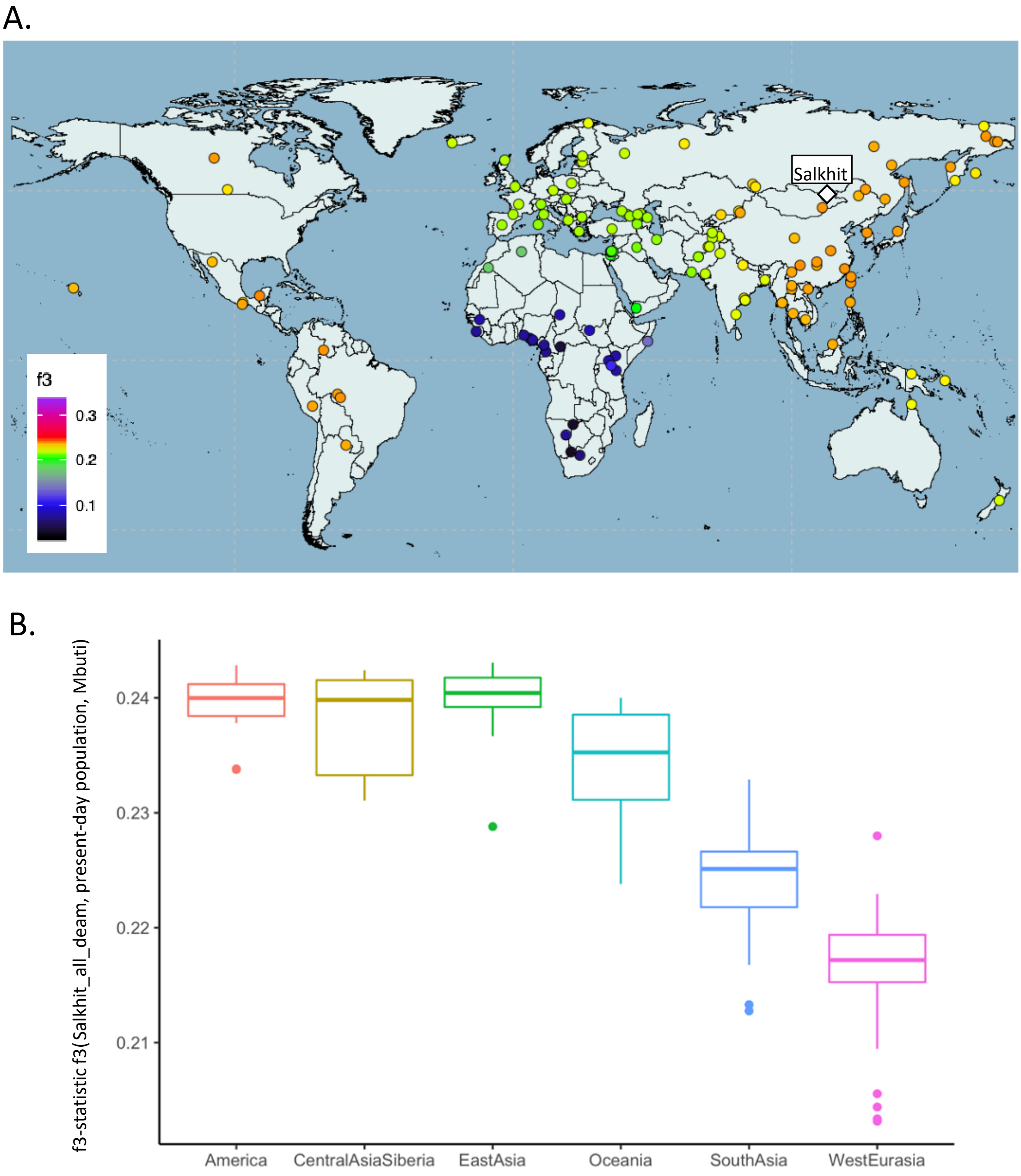


**Fig. S9. Genetic similarity computed by f3-statistic between the Salkhit individual and present-day populations.** A) Heat map for *f3*(Salkhit, X, Mbuti) where x is a present-day human population. B) Boxplot showing the f3-statistic between the Salkhit individual and present-day Native Americans, Central Asians and Siberians, East Asians, Oceanians, South Asians and West Eurasians.

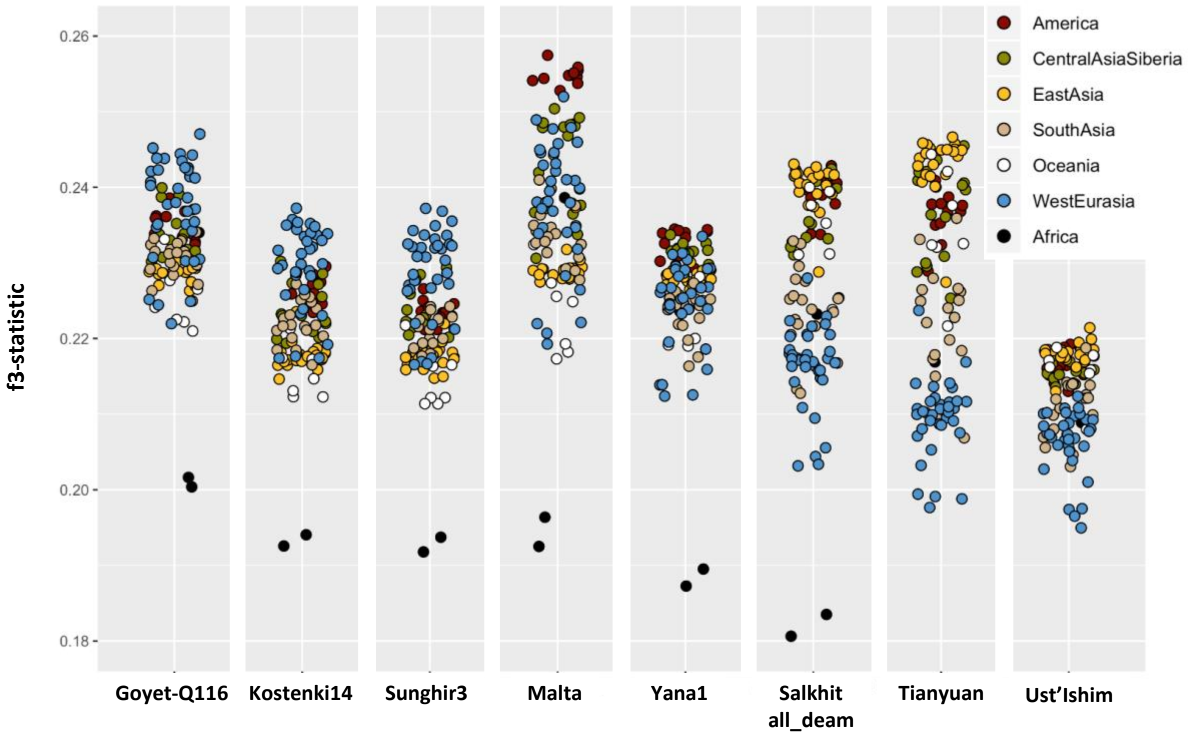


**Fig. S10 Present-day population genetic similarity to 8 ancient modern humans.** Genetic similarity between present-day populations and 8 modern human older than 20,000 years computed by f3-statistic of the form *f3(ancient modern human, present-day population, Mbuti)*.

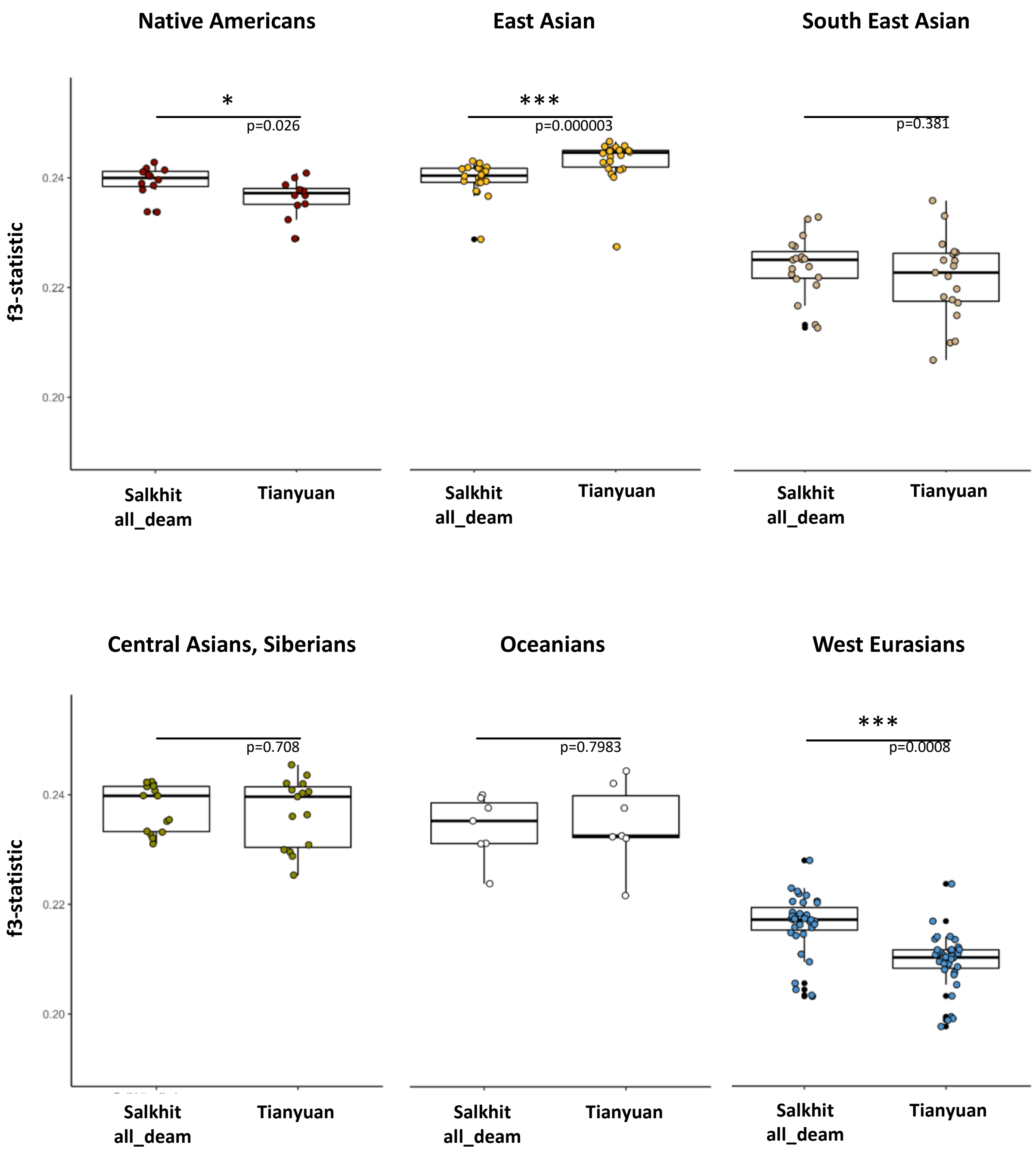


**Fig. S11. Comparison of genetic similarity between the Salkhit and the Tianyuan individuals and present-day non-African populations.** Genetic similarity computed by f3-statistic of the form f3(Salkhit/Tianyuan, present-day population, Mbuti). Significance of the difference between the f3-statistic of Salkhit and Tianyuan to a given population was assessed by an unpaired two-samples Wilcoxon test, the difference is considered significant when the p-value is higher than the significance level alpha = 0.05.

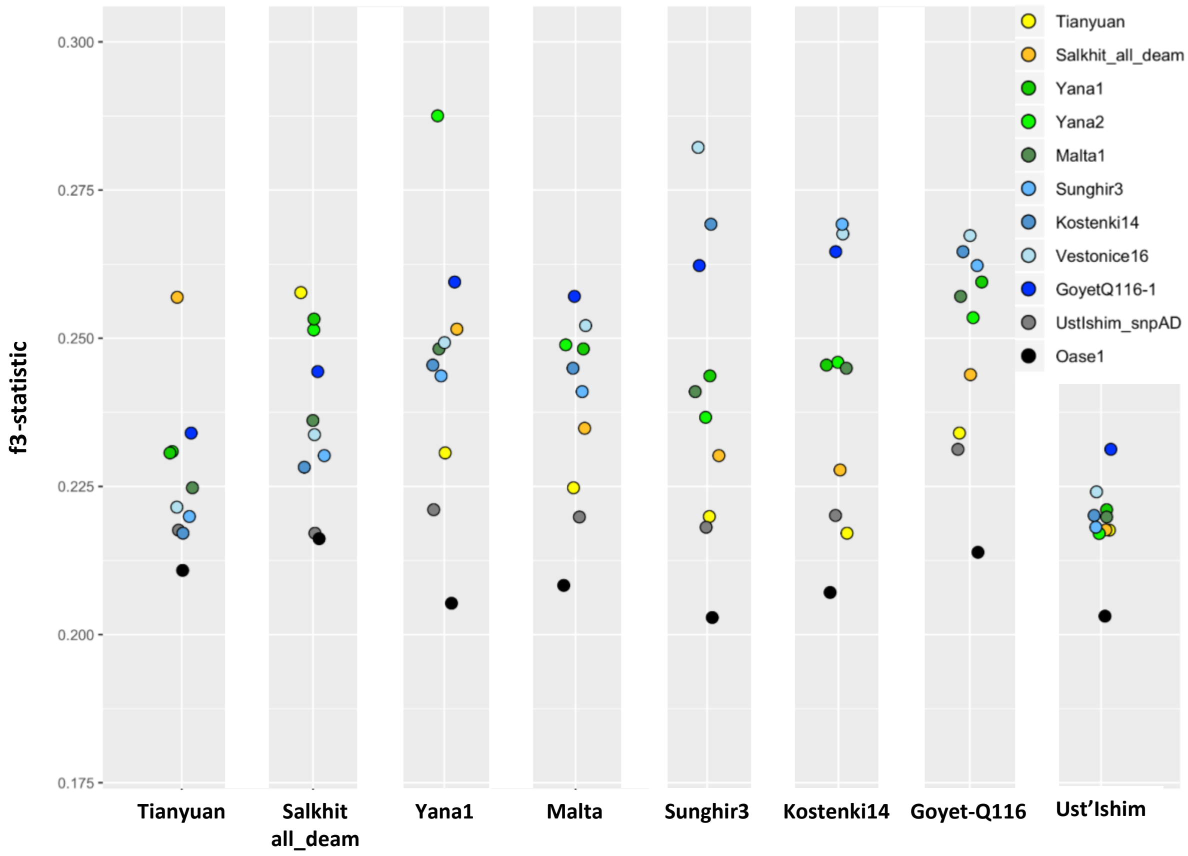


**Fig. S12. Genetic similarity between ancient modern human.** Plot of the genetic similarity computed by f3-statistic of the forme f3(ancient individual1, ancient individual2, Mbuti).

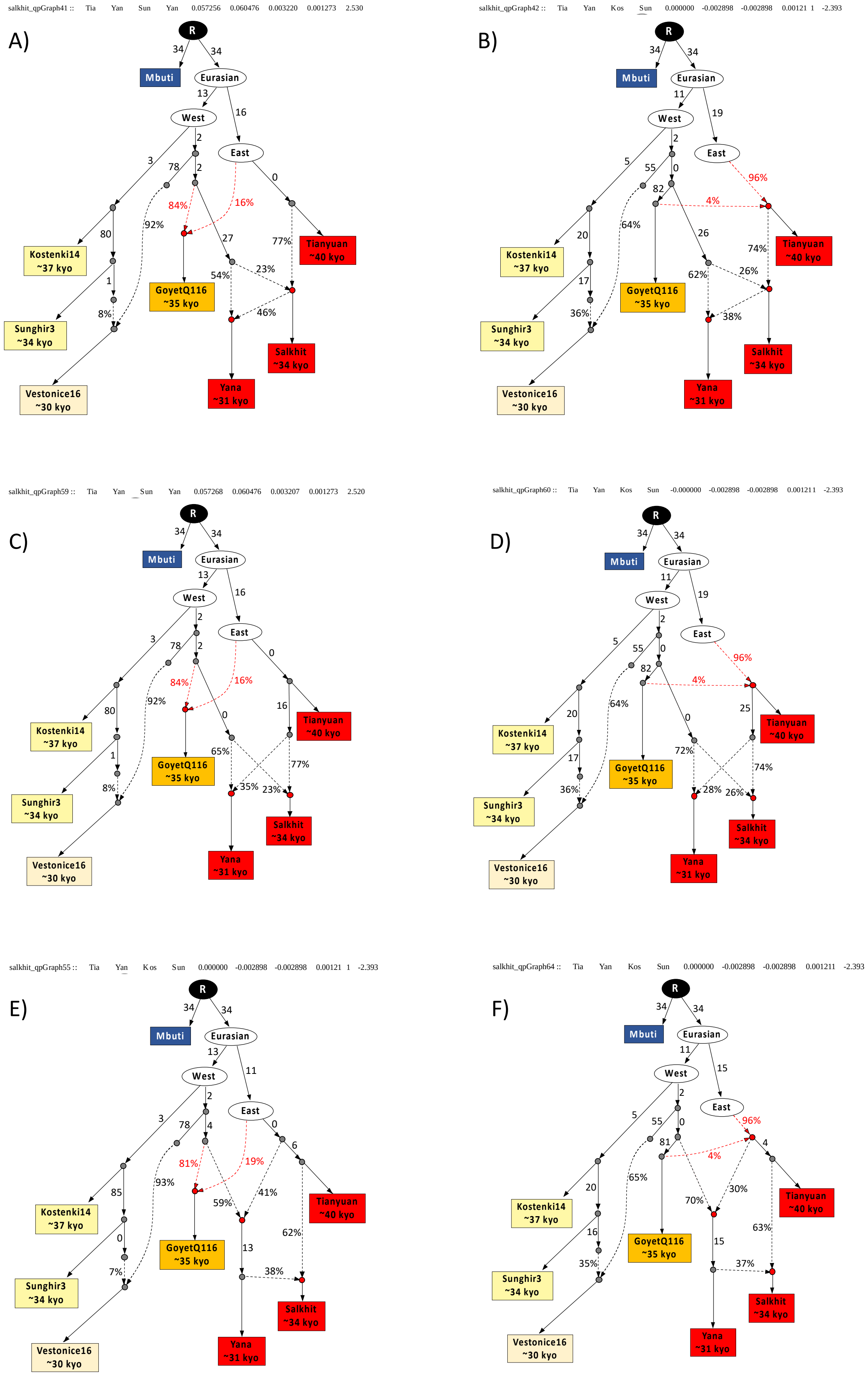

salkhit_qpGraph41 :: Tia Yan Sun Yan 0.057256 0.060476 0.003220 0.001273 2.530
A)
salkhit_qpGraph42 :: Tia Yan Kos Sun 0.000000 -0.002898 -0.002898 0.00121 1 -2.393
B)
salkhit_qpGraph59 :: Tia Yan Sun Yan 0.057268 0.060476 0.003207 0.001273 2.520
C)
salkhit_qpGraph60 :: Tia Yan Kos Sun -0.000000 -0.002898 -0.002898 0.001211 -2.393
D)
salkhit_qpGraph55 :: Tia Yan Kos Sun 0.000000 -0.002898 -0.002898 0.00121 1 -2.393
E)
salkhit_qpGraph64 :: Tia Yan Kos Sun 0.000000 -0.002898 -0.002898 0.001211 -2.393
F)
R
Mbuti
Eurasian
West
East
Kostenki14 ~37 kyo
Sunghir3 ~34 kyo
Vestonice16 ~30 kyo
GoyetQ116 ~35 kyo
Tianyuan ~40 kyo
Salkhit ~34 kyo
Yana ~31 kyo

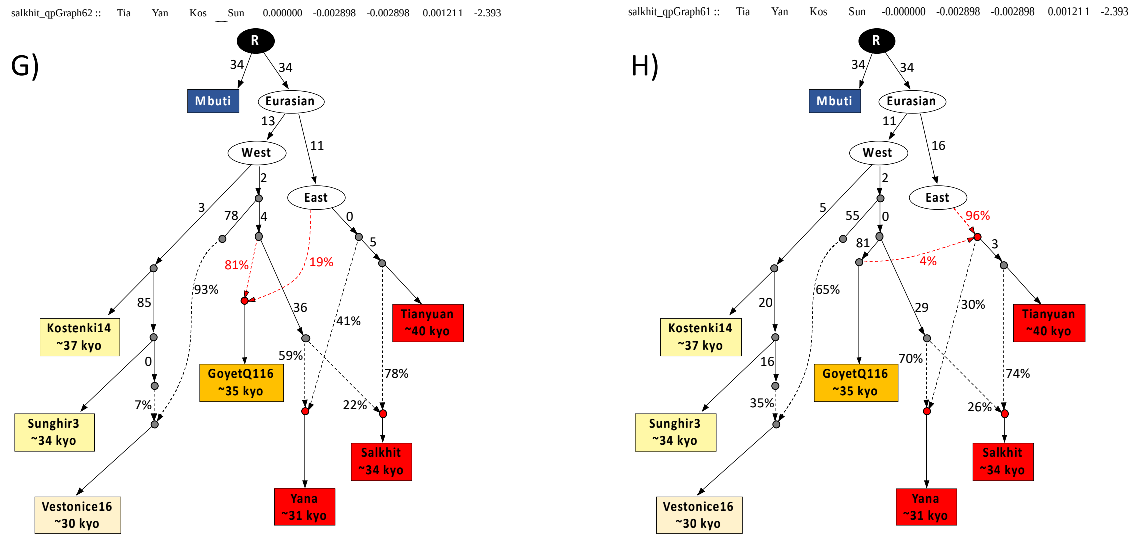


**Figure S13 Admixture graph models of the relationships of Salkhit and other Upper Pleistocene modern humans.** Solid lines represent descent by drift with branch length in unit of genetic drift and dashed lines represent admixture with admixture proportion indicated in percent. The West Eurasian ancestry in the Salkhit individual and the East Asian ancestry in the Yana individuals is explained by gene flow in both directions between populations ancestral to the Yana and Salkhit individuals after the latter diverged from the Tianyuan individual (models A, B, C, D) or between populations ancestral to the Yana individuals and Early East Asians before the divergence of the Salkhit and Tianyuan individuals (models E, F, G, H). The affinity between the 35,000-year-old Goyet-Q116 individuals and Early East Asians can be explained either by gene flow from Early East Asians into Goyet-Q116 ancestors (models A, C, E, G) or from Goyet-Q116 ancestors into Early East Eurasians (models B, D, F, H).
All models fit the data with Z scores between -3 and 3 (indicated above each graph).

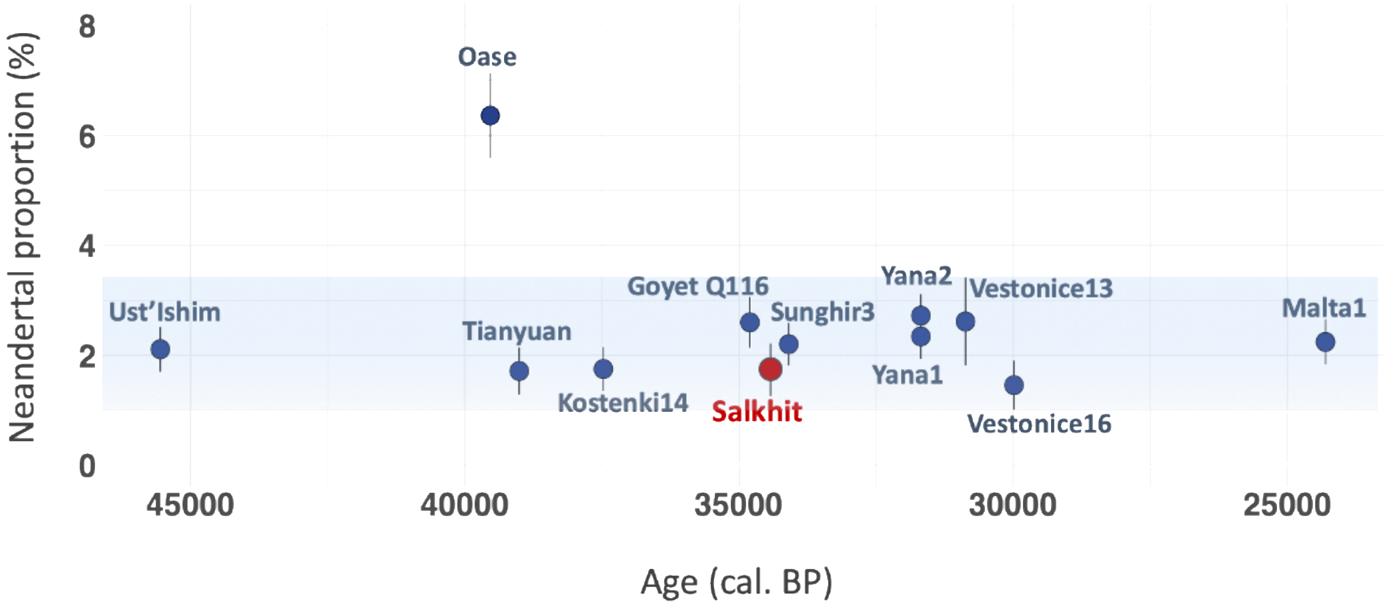


**Fig. S14 Neandertal ancestry in the Salkhit individual and other early modern humans.** The proportion of Neandertal ancestry was estimated by *f4*-ratio statistic using two high coverage Neandertal genomes.

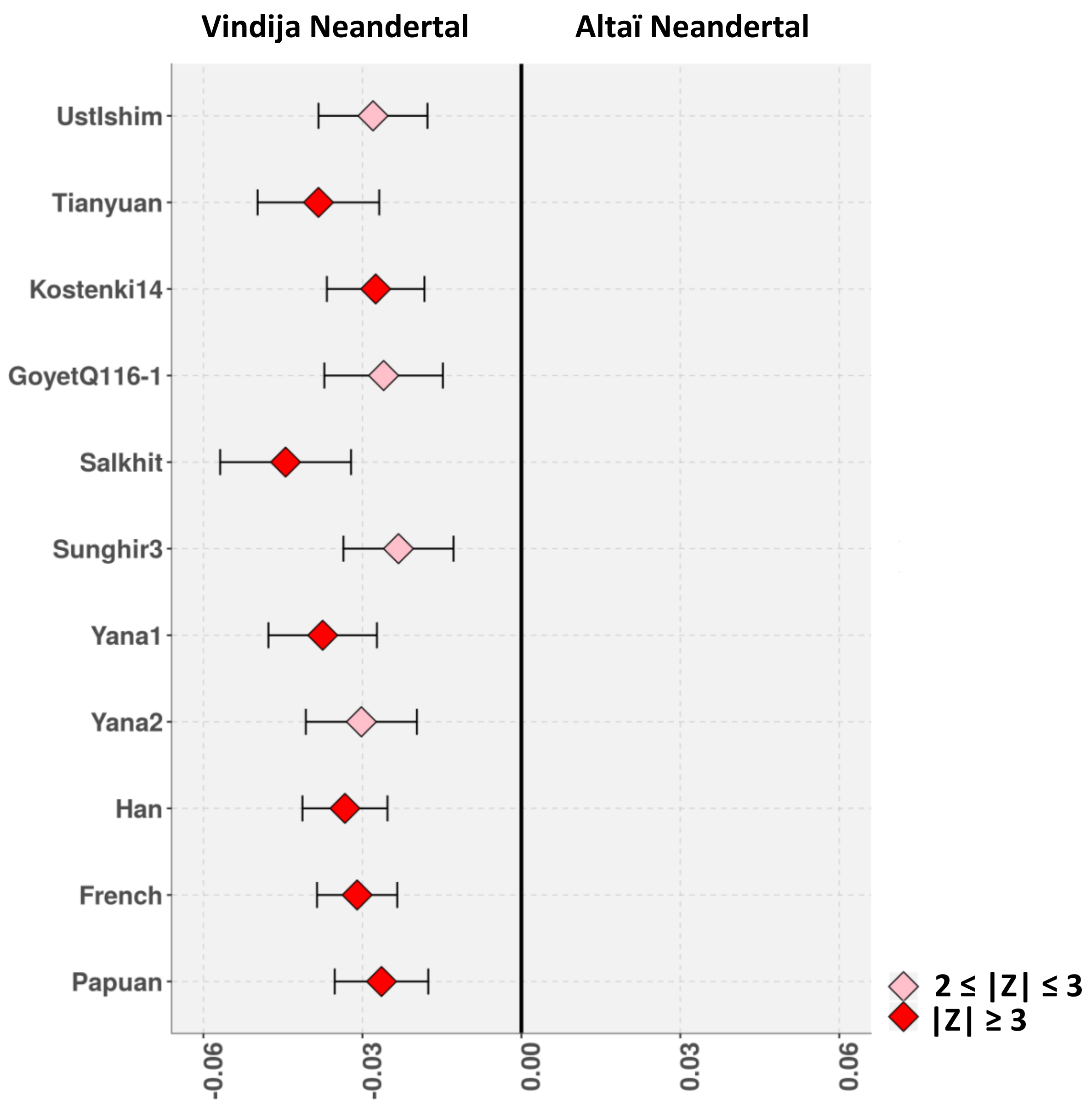


**Fig. S15 Neandertal ancestry in Salkhit and other modern humans.** D-statistic of the form *D(Altaï Neandertal, Vindija Neandertal; X, Mbuti)* when X is an early modern human individual or a present-day population to assess to which of the two high coverage neandertal the neandertal ancestry of those individuals is closer to.

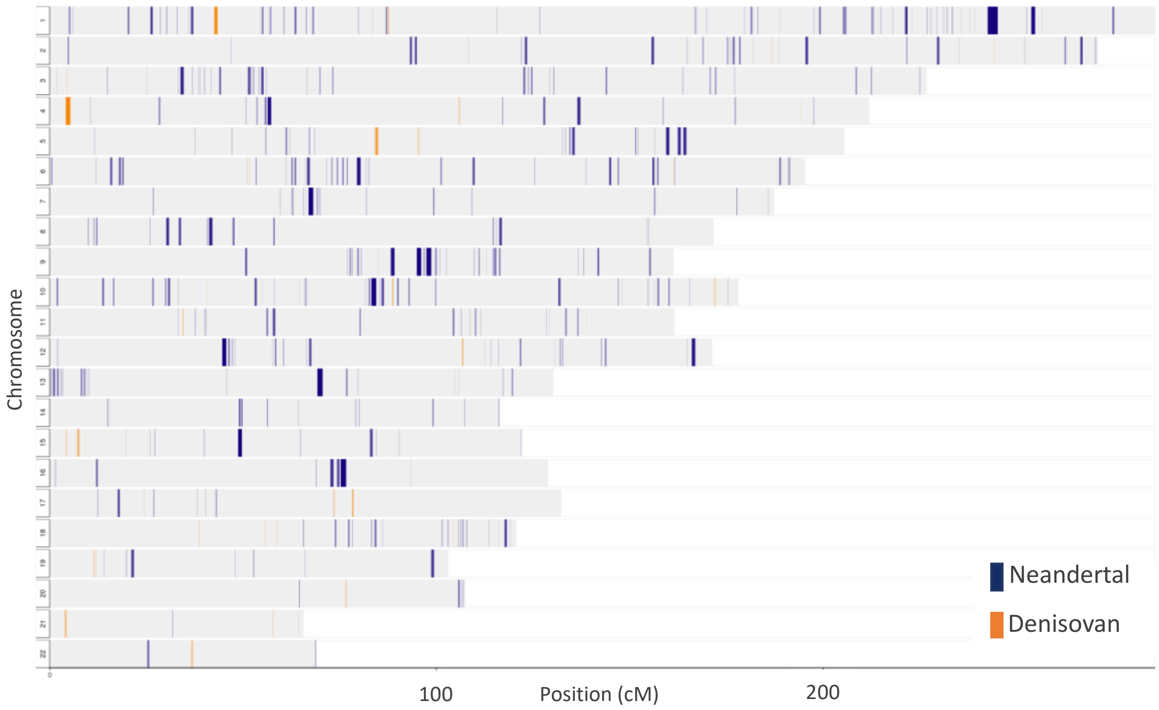


**Fig. S16 Neandertal (blue) and Denisovan (orange) ancestry tracks in the Salkhit genome inferred using the *admixfrog* program.**

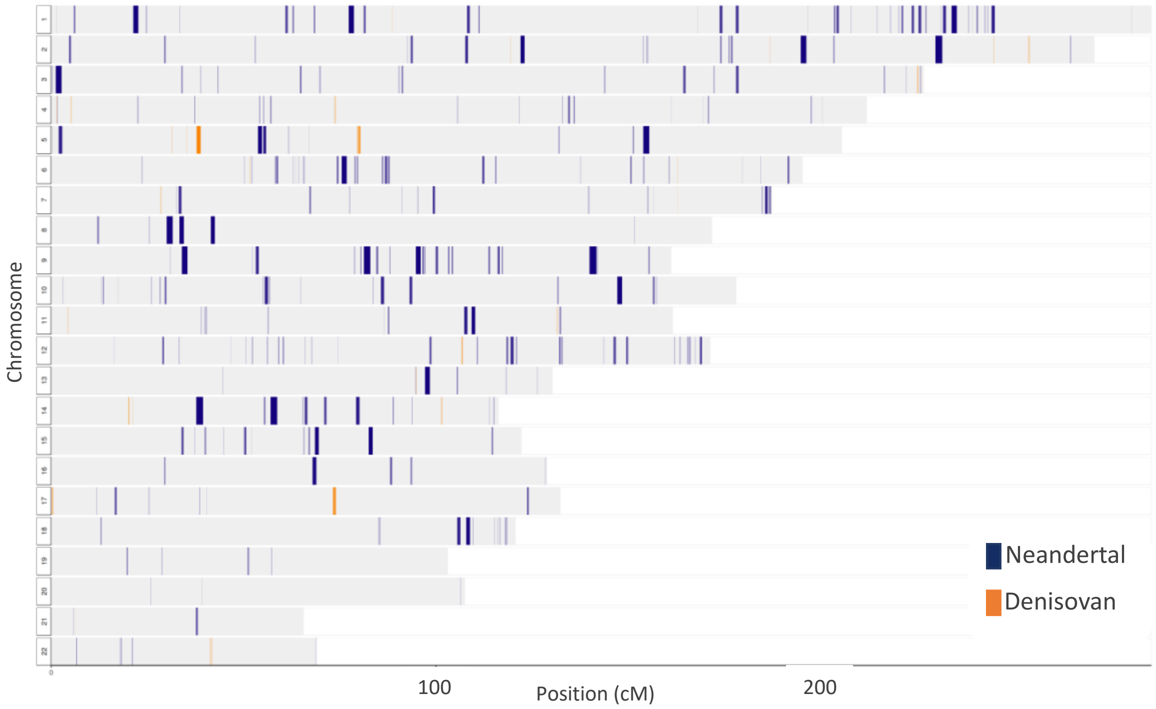


**Fig. S17 Neandertal (blue) and Denisovan (orange) ancestry tracks in the Tianyuan genome inferred using the *admixfrog* program.**

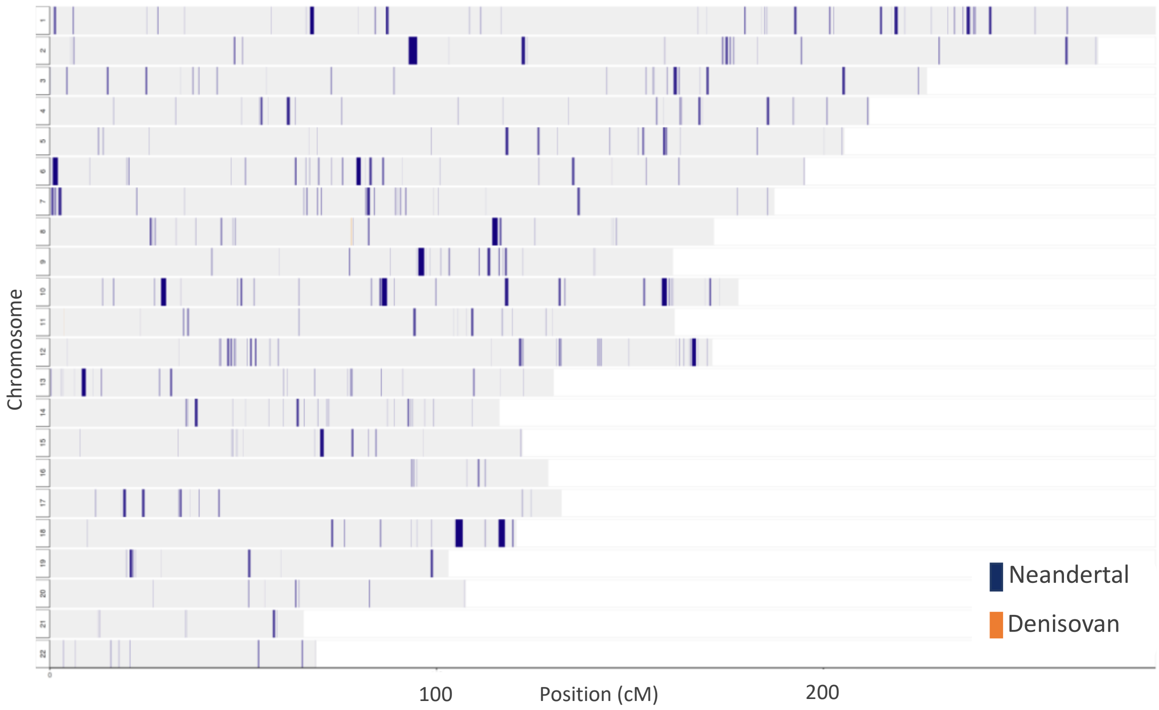


**Fig. S18. Neandertal (blue) and Denisovan (orange) ancestry tracks in the *Yana1* genome inferred using the *admixfrog* program.**

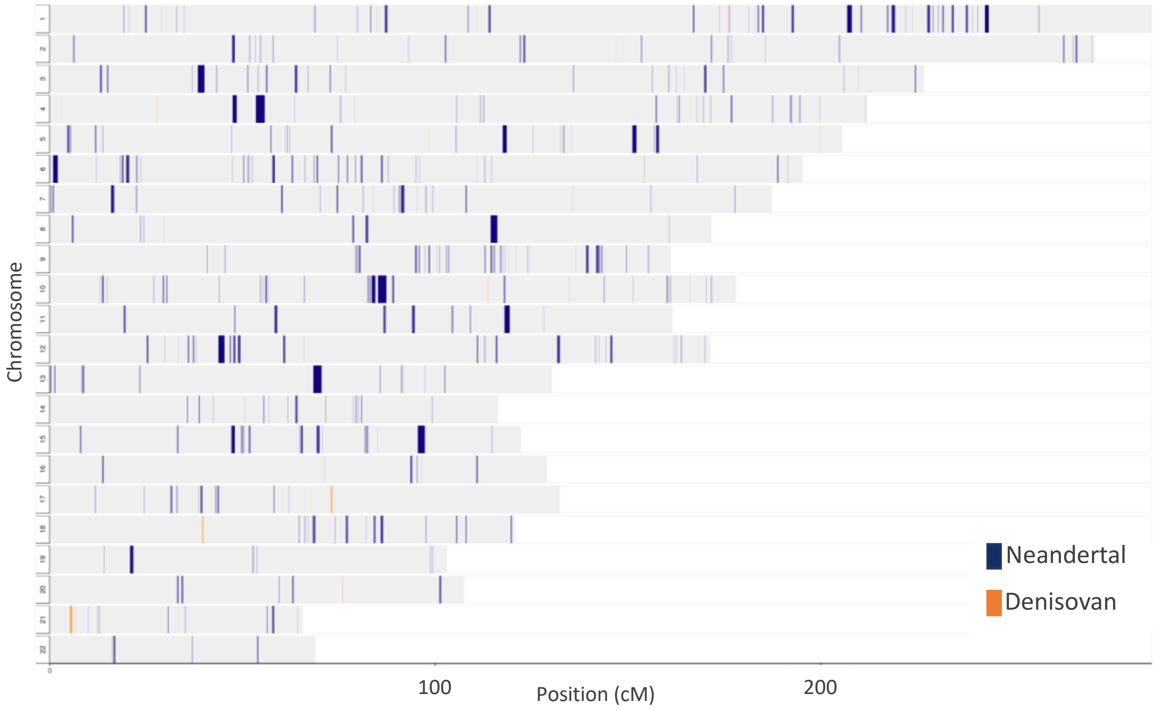


**Fig. S19 Neandertal (blue) and Denisovan (orange) ancestry tracks in the *Yana2* genome inferred using the *admixfrog* program.**

.

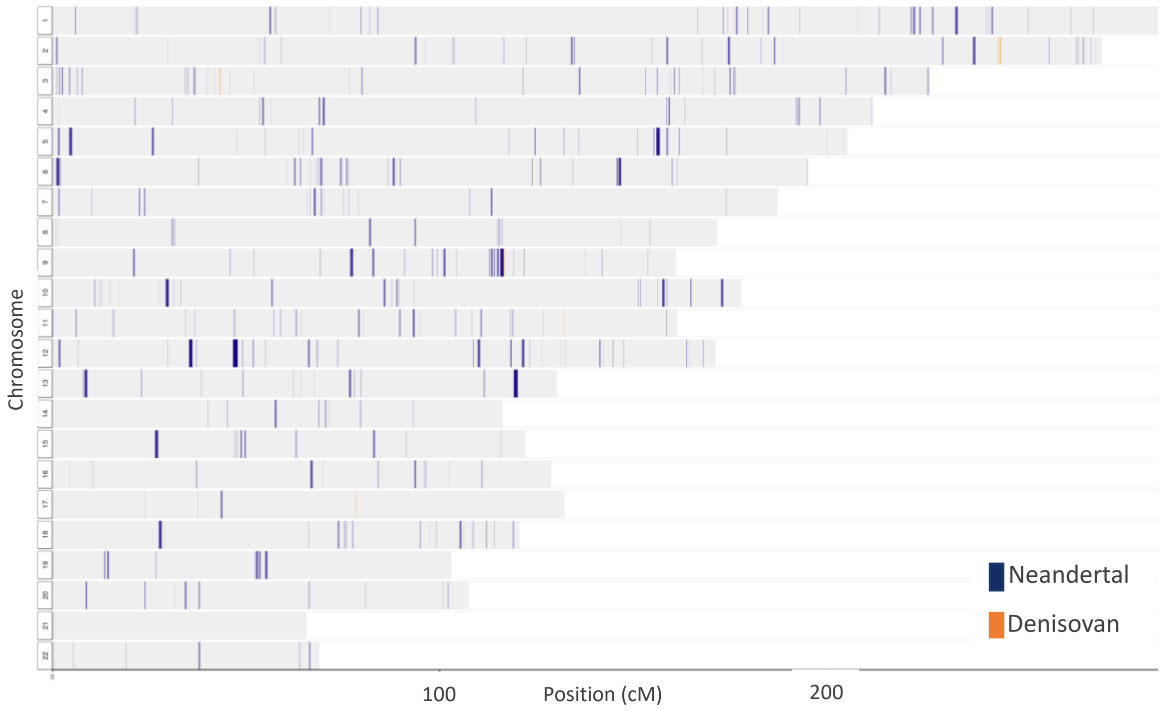


**Fig. S20 Neandertal (blue) and Denisovan (orange) ancestry tracks in the *Mal'ta 1* genome inferred using the *admixfrog* program.**

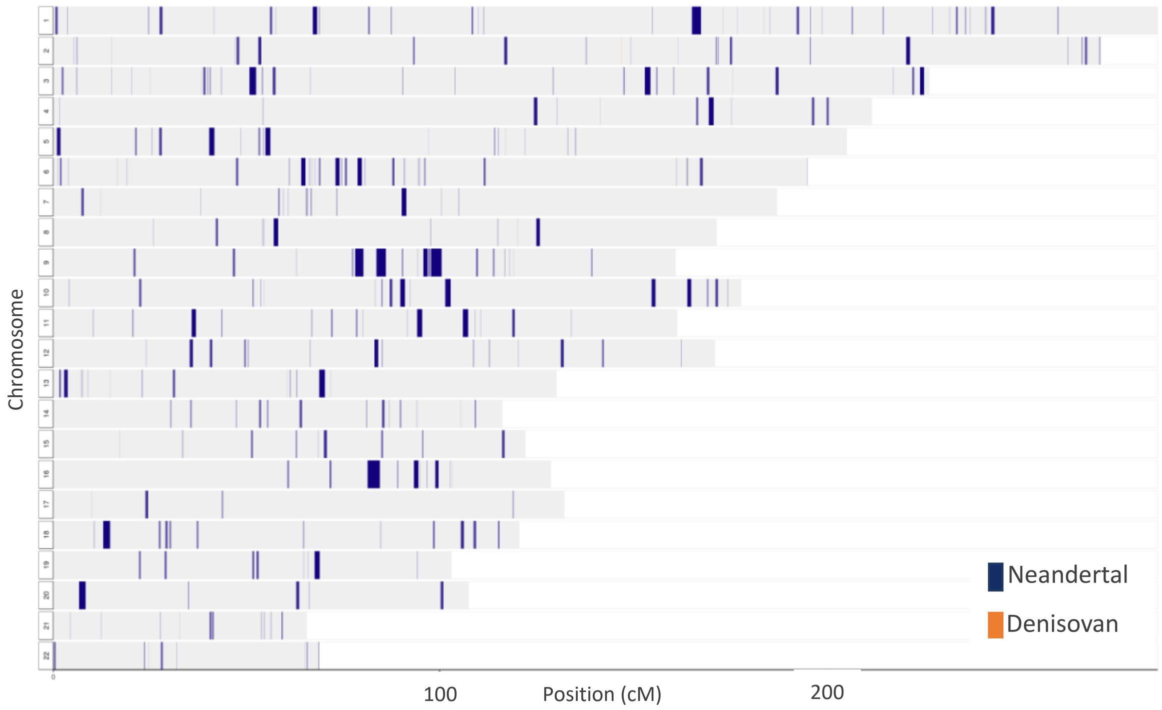


**Fig. S21. Neandertal (blue) and Denisovan (orange) ancestry tracks in the *Kostenki14* genome inferred using the *admixfrog* program.**

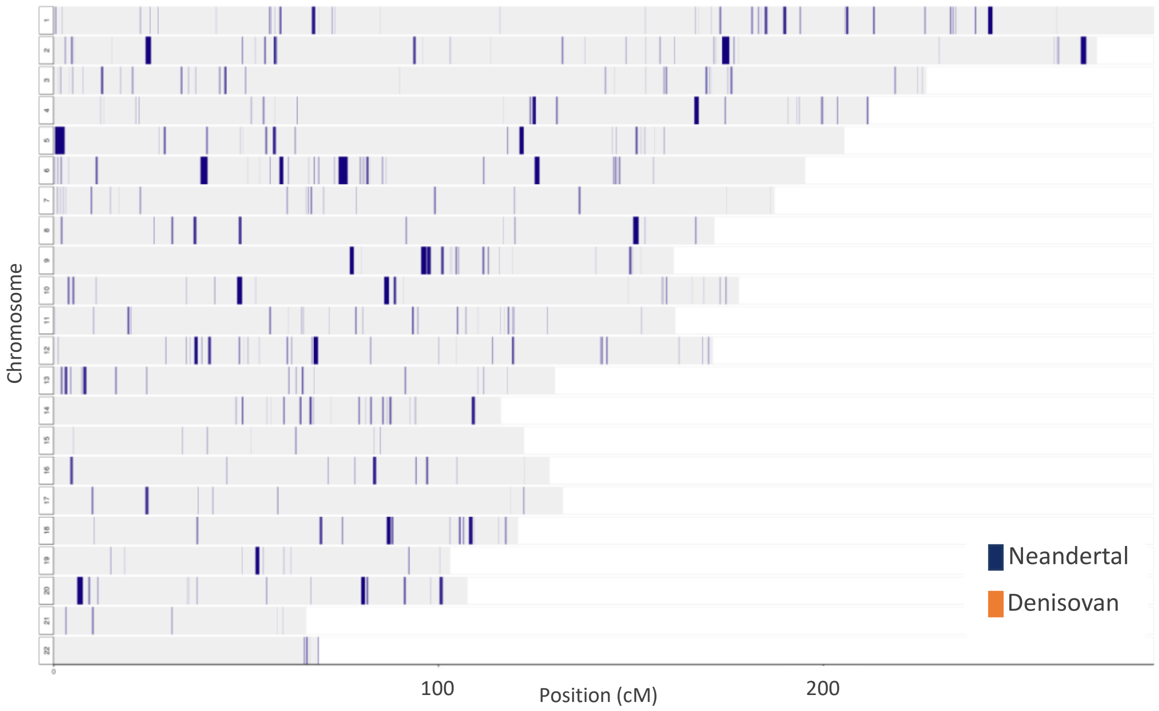


**Fig. S22 Neandertal (blue) and Denisovan (orange) ancestry tracks in the *Sunghir3* genome inferred using the *admixfrog* program.**

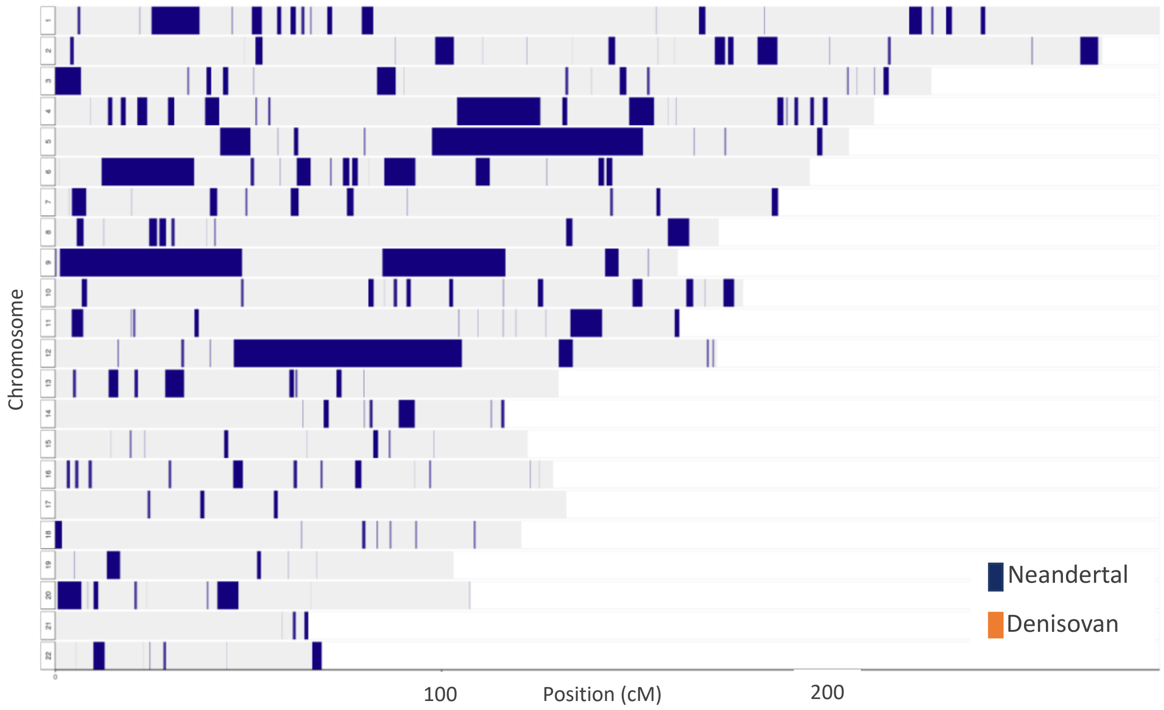


**Fig. S23 Neandertal (blue) and Denisovan (orange) ancestry tracks in the *Oase1* genome inferred using the *admixfrog* program.**

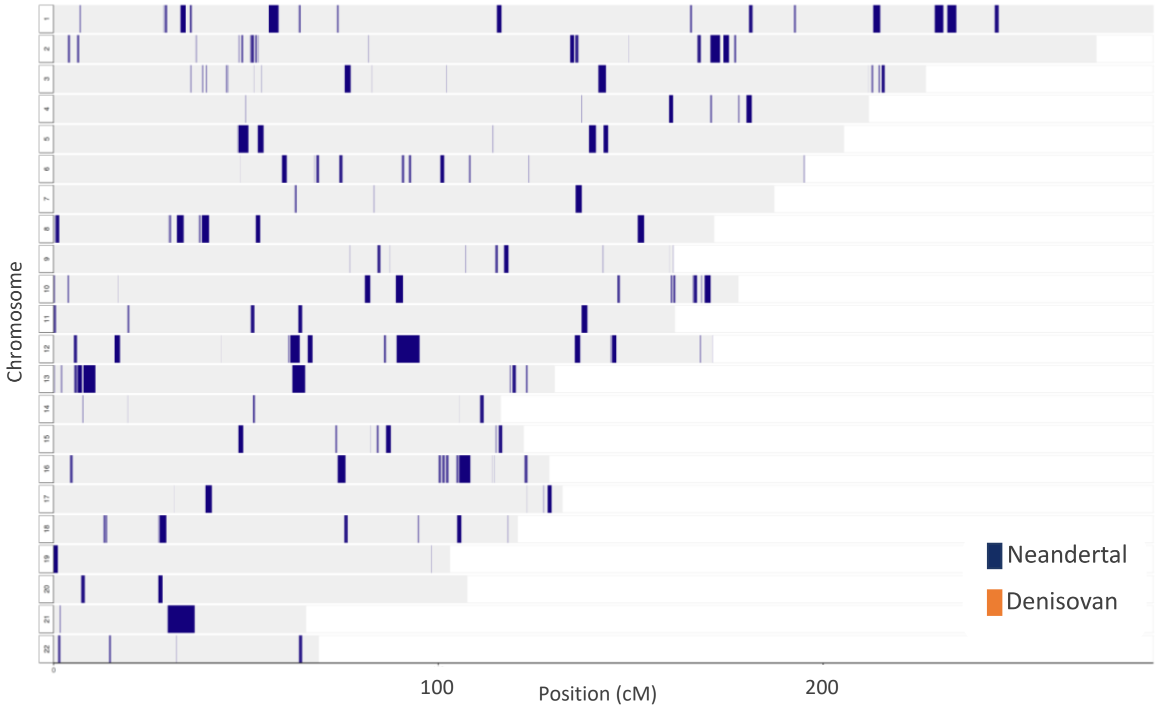


**Fig. S24 Neandertal (blue) and Denisovan (orange) ancestry tracks in the Ust'Ishim genome inferred using the *admixfrog* program.**

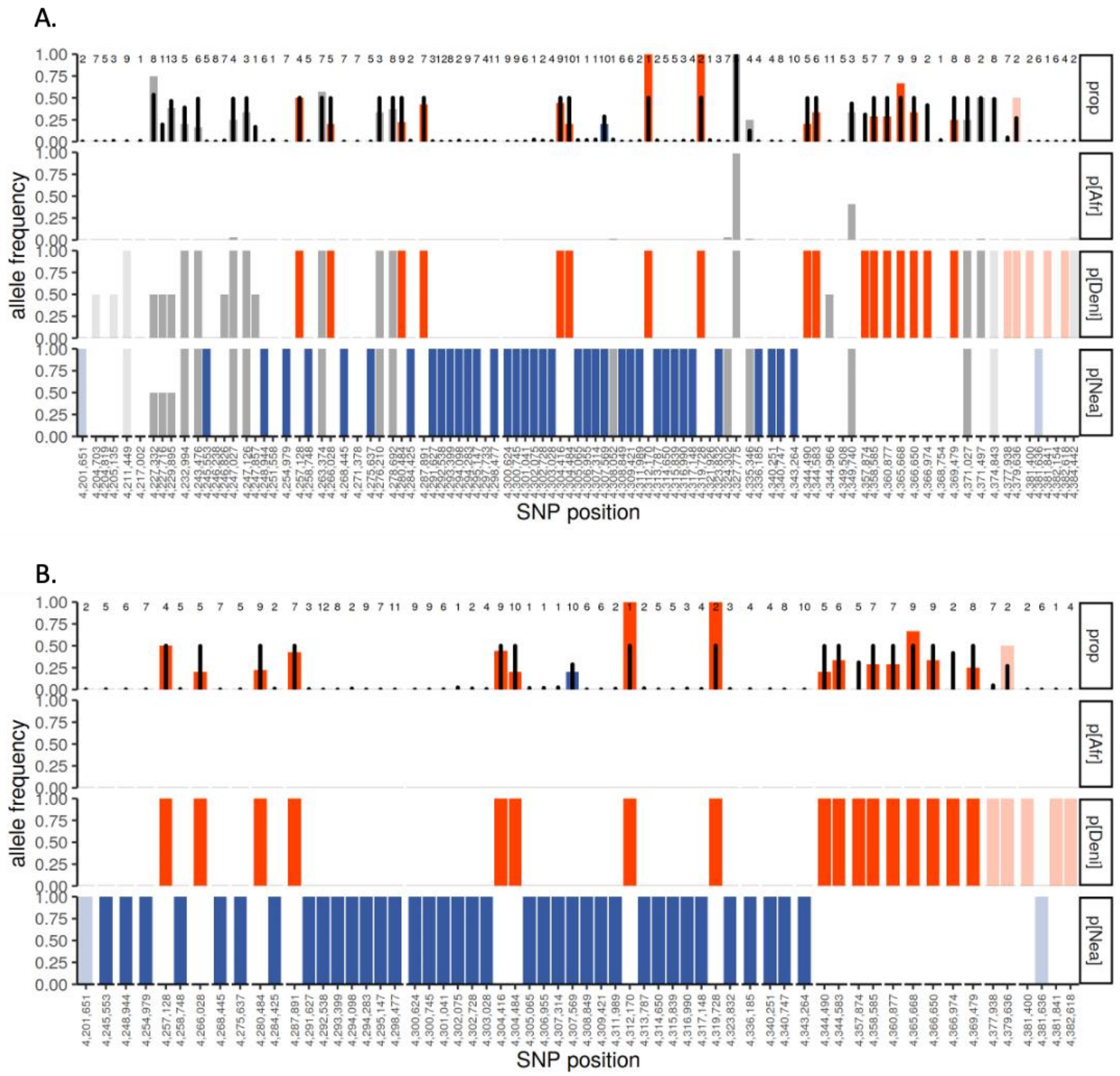


**Fig. S25 Allele frequencies for SNPs in the longest Denisovan introgressed track of the Salkhit genome**. The three lower panels present the frequencies for Neandertal-like alleles (blue bars), Denisovan-like alleles (orange bars), or both archaic-like alleles (grey bars) African-like alleles (no bar) in three reference genomes: a present-day African individual (panel p[Afr]), a Denisovan individual (panel p[Deni]) and a Neandertal individual (panel p[Nea]). The upper panel [prop] presents the number of DNA fragment covering each SNPs of the targeted track and carrying an archaic allele, Neandertal in blue highlight, Denisovan in orange highlight or shared in grey highlight. A black bar underlines the presence of an archaic allele. A) Allele frequencies of all SNPs in the targeted track; B) Allele frequency of SNPs carrying either a derived Denisovan or Neandertal allele.

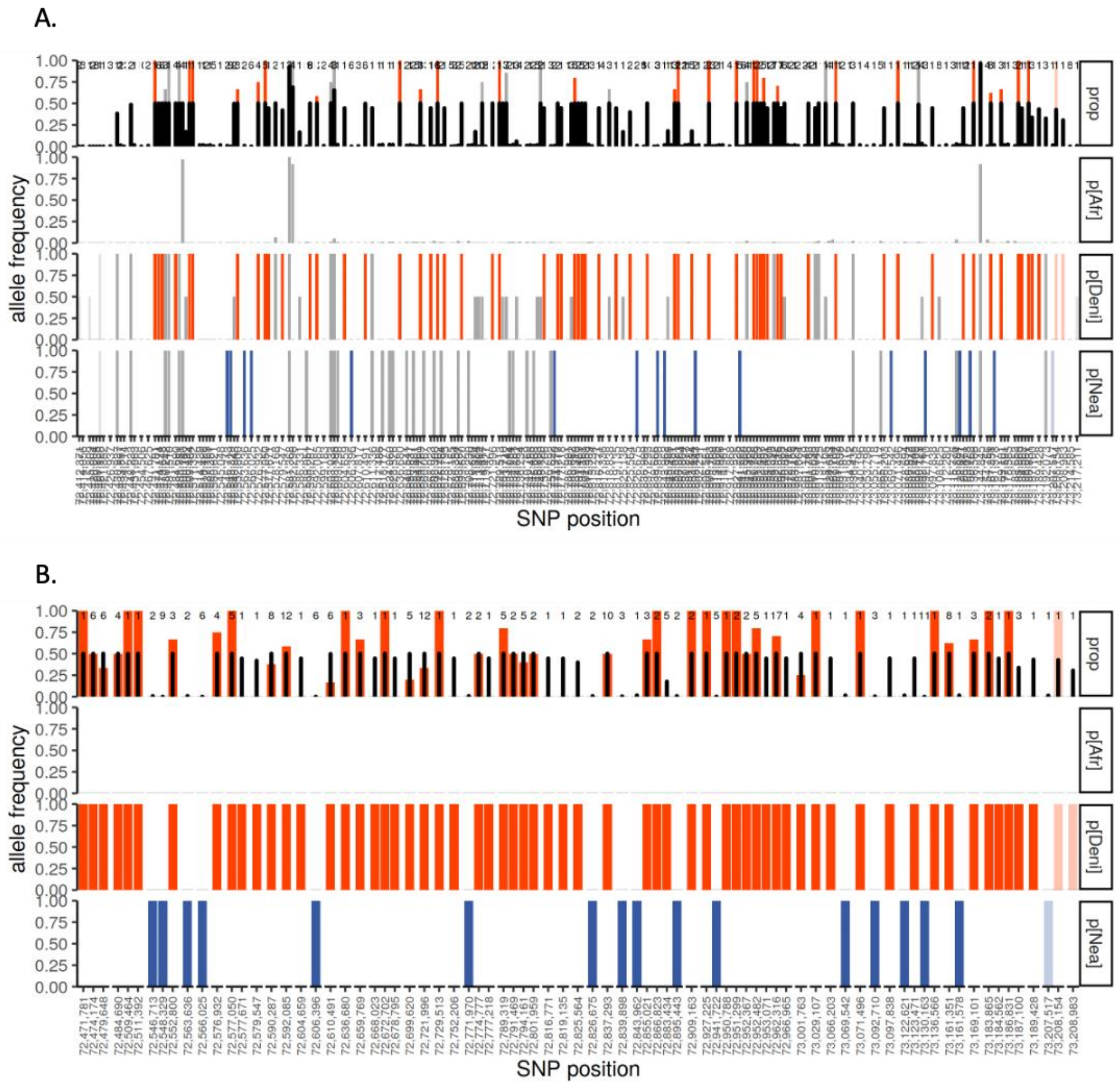


**Fig. S26 Allele frequencies for SNPs in the longest Denisovan introgressed track of the Tianyuan genome**. The three lower panels present the frequencies for Neandertal-like alleles (blue bars), Denisovan-like alleles (orange bars), or both archaic-like alleles (grey bars) African-like alleles (no bar) in three reference genomes: a present-day African individual (panel p[Afr]), a Denisovan individual (panel p[Deni]) and a Neandertal individual (panel p[Nea]). The upper panel [prop] presents the number of DNA fragment covering each SNPs of the targeted track and carrying an archaic allele, Neandertal in blue highlight, Denisovan in orange highlight or shared in grey highlight. A black bar underlines the presence of an archaic allele. A) Allele frequencies of all SNPs in the targeted track; B) Allele frequency of SNPs carrying either a derived Denisovan or Neandertal allele.

.

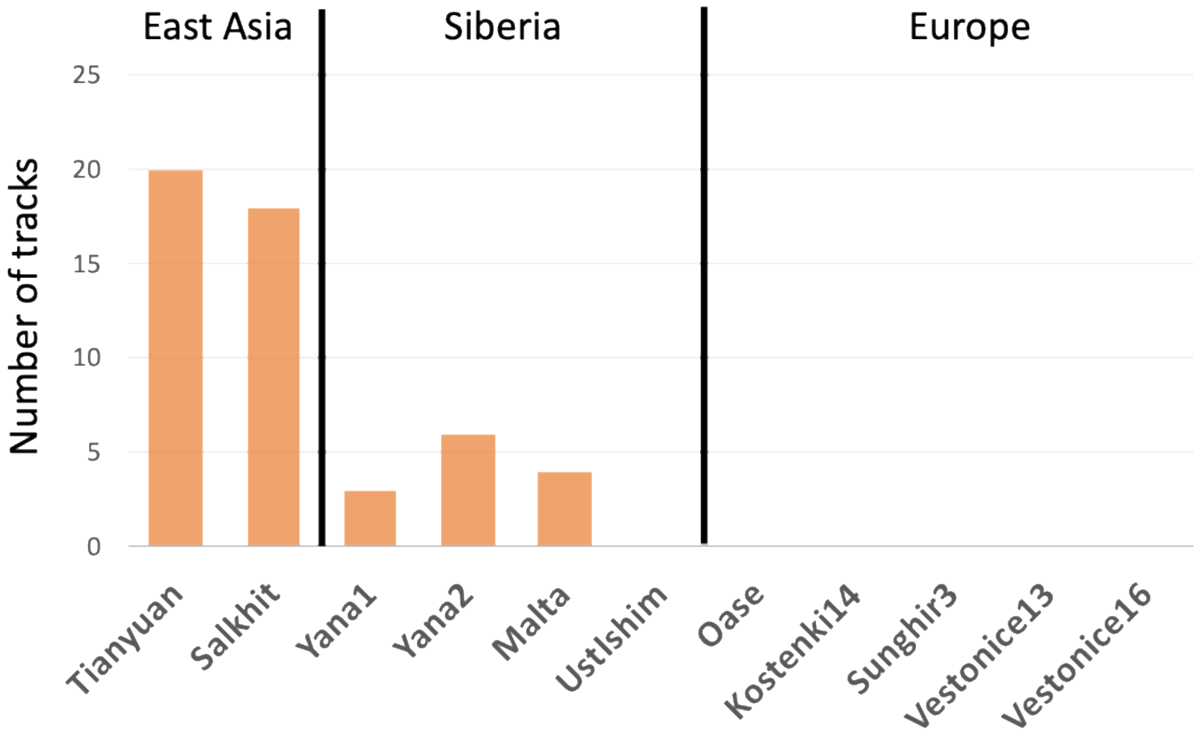


**Fig. S27 Denisovan ancestry in early modern humans in Eurasia.** Number of Denisovan ancestry tracks longer than 0.2cm in 10 early modern humans from Eurasia inferred using the *admixfrog* program.

.

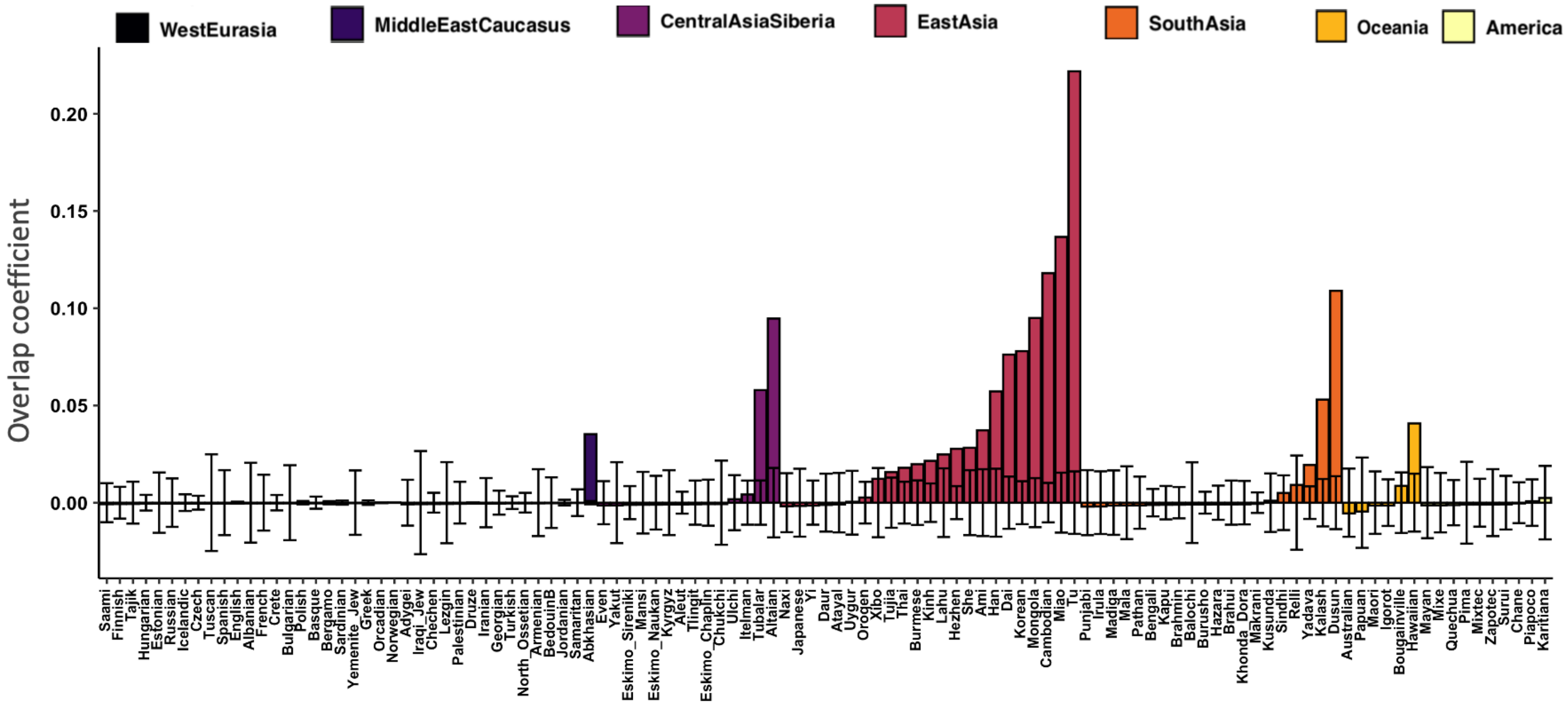


**Fig. S28 Overlap between Denisovan ancestry tracks in the Salkhit genome and in present-day populations.** Each bar represents the correlation coefficient of the overlap between Denisovan ancestry tracks in the Salkhit genome and in each of the present-day population. The confidence interval is generated by 500 bootstrap iteration. A coefficient of correlation higher than the confidence interval indicates that the overlap is higher than what is expected by chance.

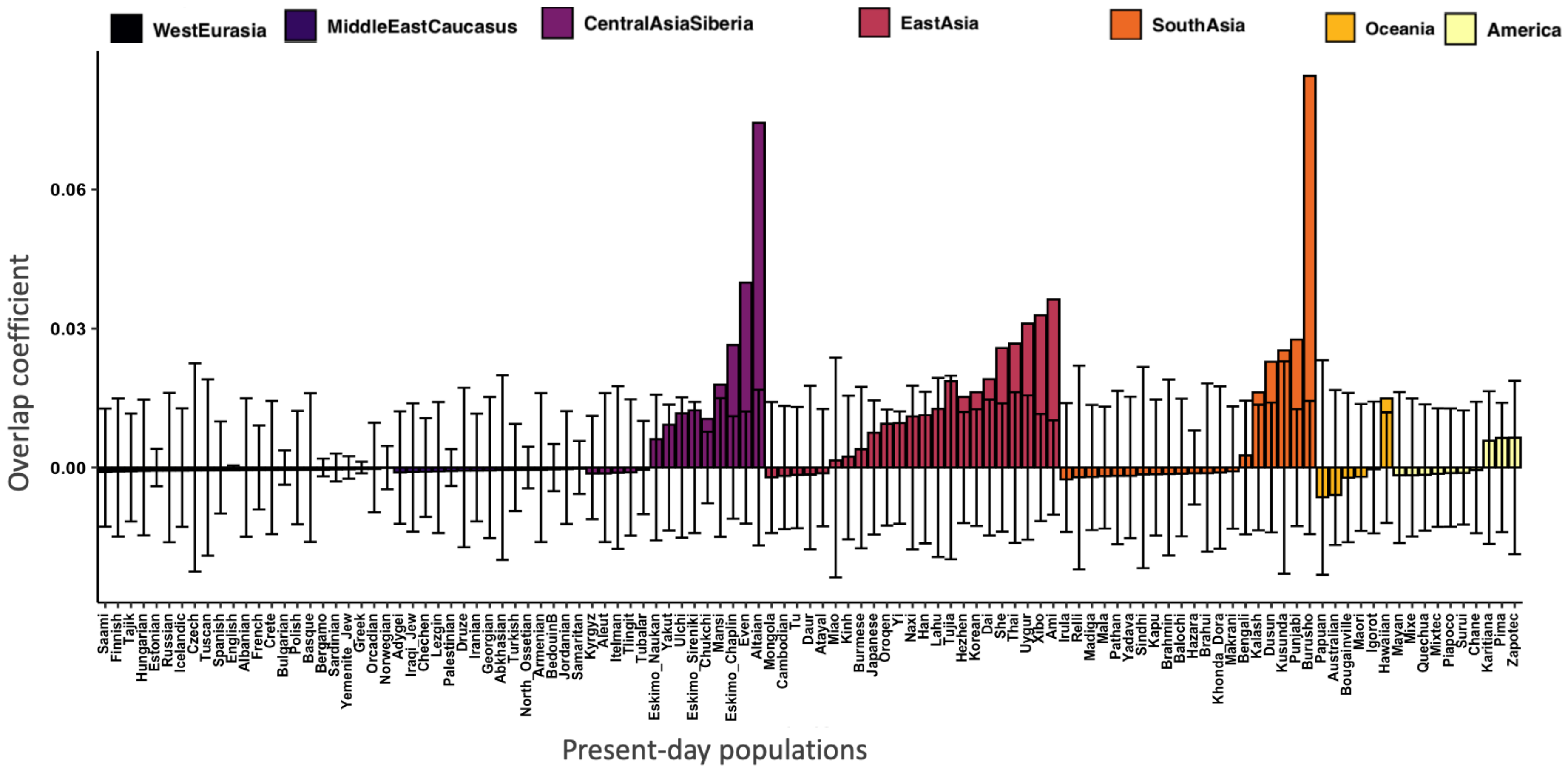


**Fig. S29 Overlap between the Denisovan ancestry tracks in the Tianyuan genome and present-day populations.** Each bar represents the correlation coefficient of the overlap between Denisovan ancestry tracks in the Salkhit genome and in each of the present-day population. The confidence interval is generated by 500 bootstrap iteration. A coefficient of correlation higher than the confidence interval indicates that the overlap is higher than what is expected by chance.

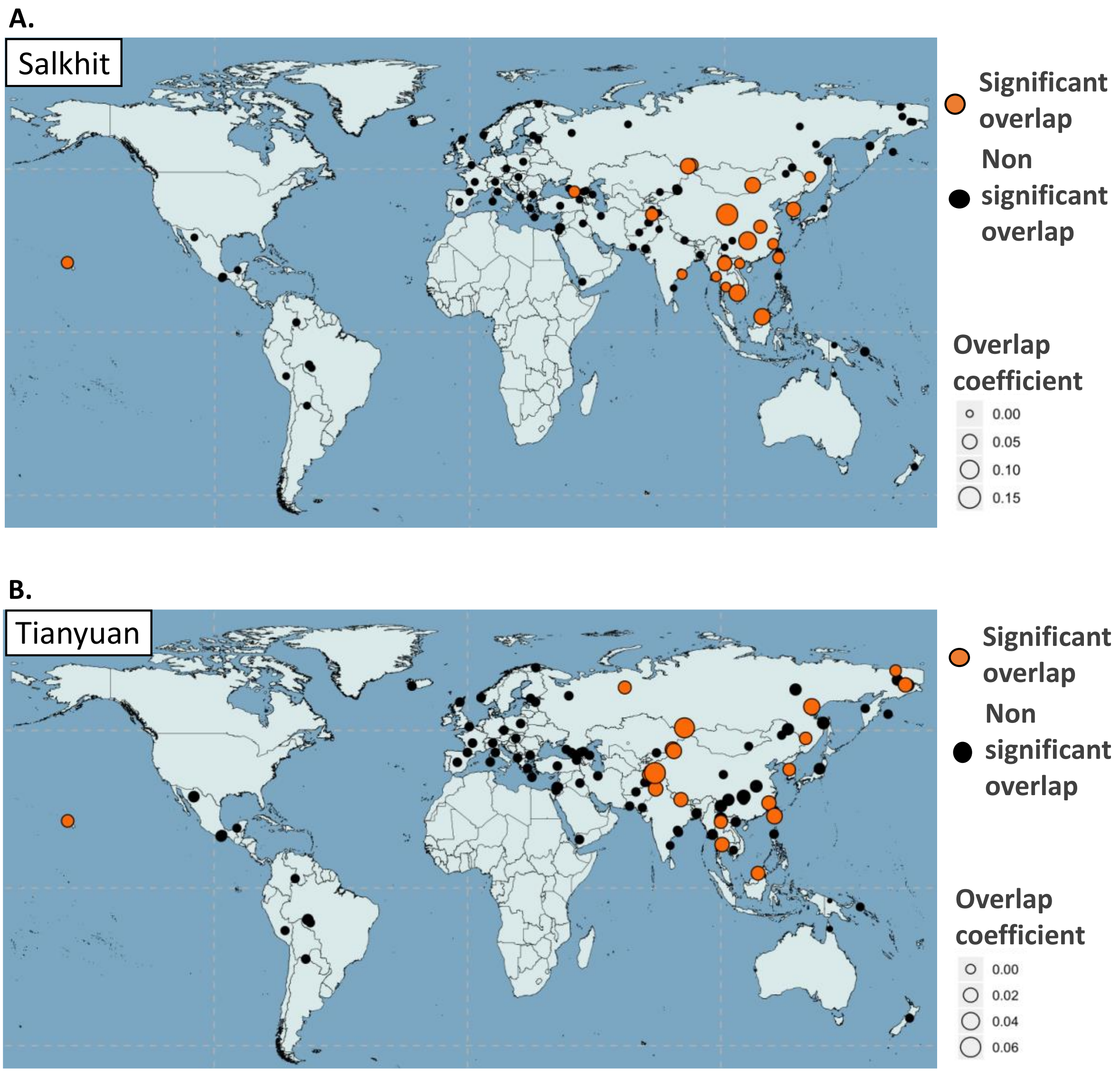


**Fig. S30 Overlap of the Denisovan ancestry tracks in the Salkhit (A) and Tianyuan (B) genomes and present-day populations.** Orange circles indicate the geographic locations of present-day populations from which we see a correlation of Denisovan ancestry with the Salkhit individual (A) or the Tianyuan individual (B).

| Sample | Sex | Archaeological site | Geographic Origin | Country | Latitude | Longitude | Uncalibrated (BP) | | Calibrated (IntCal13) (BP) | | | 2.2 million SNPs coverage | | Source |
|---|---|---|---|---|---|---|---|---|---|---|---|---|---|---|
| | | | | | | | ages | ± | upper | lower | mean | number | % | |
| Salkhit | F | out of context | EastAsia | Mongolia | 48.27 | 112.36 | 30,430 | 300 | 34,950 | 33,900 | 34,425 | 884,795 | 41.25% | This study. |
| Tianyuan | M | Tianyuan Cave | EastAsia | China | 39.65 | 115.87 | 34,430 | 510 | 40,254 | 37,761 | 39,007 | 1,474,871 | 68.77% | Yang & al, 2017 |
| Yana1 | M | Yana RHS | Siberia | Russia | 70.7 | 135.4 | 27,940 | 115 | 32,047 | 31,321 | 31,684 | 1,959,414 | 91.37% | Sikora & al, unpublished |
| Yana2 | M | Yana RHS | Siberia | Russia | 70.7 | 135.4 | 27,940 | 115 | 32,047 | 31,321 | 31,684 | 1,956,460 | 91.23% | Sikora & al, unpublished |
| Kolyma1 | M | Duvanni Yar | Siberia | Russia | 68.6 | 159.1 | 8,770 | 27 | 9,904 | 9,668 | 9,786 | 1,958,983 | 91.35% | Sikora & al, unpublished |
| Goyet-Q116 | M | Troisième Caverne of Goyet | WestEurasia | France | 50.26 | 4.28 | 30,880 | 170 | 35,166 | 34,436 | 34,801 | 846,983 | 39.49% | Fu & al, 2016 |
| Ostuni1 | F | The Grotta di Santa Maria di Agnano | WestEurasia | Italy | 40.73 | 17.57 | 23,446 | 107 | 27,810 | 27,430 | 27,620 | 369,313 | 17.22% | Fu & al, 2016 |
| Malta1 | M | Mal'ta | CentralAsiaSiberia | Russia | 52.9 | 103.5 | 20,240 | 60 | 24,520 | 24,090 | 24,305 | 1,439,501 | 67.12% | Raghavan & al, 2014 |
| Vestonice16* | M | Dolní Vēstonice I | WestEurasia | Italy | 48.53 | 16.39 | 25,740 | 210 | 30,575 | 29,395 | 29,985 | 945,292 | 44.08% | Fu & al, 2016 |
| KremsWA3* | M | Krems-Wachtberg | WestEurasia | Austria | 48.41 | 15.59 | 26,870 | 220 | 31,250 | 30,690 | 30,970 | 236,831 | 11.04% | Fu & al, 2016 |
| ElMiron | F | El Mirón Cave | WestEurasia | Spain | 43.26 | -3.45 | 15,460 | 40 | 18,830 | 18,610 | 18,720 | 797,714 | 37.2% | Fu & al, 2016 |
| Muierii2 | F | Muierilor Cave | WestEurasia | Romania | 45.11 | 23.46 | 29,110 | 190 | 33,760 | 32,840 | 33,300 | 98,618 | 4.6% | Fu & al, 2016 |
| Sunghir3 | M | Sunghir | WestEurasia | Russia | 56.17 | 40.5 | 30,000 | 550 | 35,154 | 33,031 | 34,093 | 1,958,733 | 91.34% | Sikora & al, 2020 |
| Vestonice13* | M | Dolní Vēstonice I | WestEurasia | Italy | 48.53 | 16.39 | 26,640 | 110 | 31,070 | 30,670 | 30,870 | 119,094 | 5.55% | Fu & al, 2016 |
| Kostenki14 | M | Markina Gora, Kostenki-Borshchevo | WestEurasia | Russia | 51.23 | 39.3 | 33,250 | 500 | 38,684 | 36,262 | 37,473 | 1,774,156 | 82.73% | Fu & al, 2016; S. Orlando & al, 2014 |
| Oase1 | M | Peștera cu Oase | WestEurasia | Romania | 45.12 | 21.9 | 34,950 | 990 | 41,761 | 37,311 | 39,536 | 285,076 | 13.29% | Fu & al, 2015 |
| UstIshim | M | out of context | CentralAsiaSiberia | Russia | 57.43 | 71.1 | 41,400 | 1400 | 48,313 | 42,793 | 45,553 | 2,137,615 | 99.68% | Fu & al, 2014 |

**Table S1. Ancient modern human dataset used in this study.**
The ages of samples marked with "*" are based on associated artifacts.

| Sample | Shotgun - Nuclear SNPs Capture | nbr of molecules in the library | Total nbr of seq. generated | Total nbr of seq. merged and filtered* | Total mapped seq. (MAPQ 25) | % mapped seq. | Unique mapped seq. | Duplication rate | average size | Average genome covered / library (bases) | % 5' C to T | % 3' C to T | Total nbr of deaminated seq. | average size of deaminated seq. | Total nbr of targetted SNPs | Nbr of SNPs covered (all seq.) | % of targetted SNPs covered (all seq.) | Nbr of SNPs covered (deam seq. only) | % of targetted SNPs covered (deam seq.only) |
|---|---|---|---|---|---|---|---|---|---|---|---|---|---|---|---|---|---|---|---|
| Salkhit skullcap - A | Shotgun | 7.69E+9 | 1,723,860 | 1,079,127 | 16,272 | 1.50 | 16,225 | 1 | 60.6 | 4.49E+9 | 22.8 | 13.1 | 1,983 | 55.3 | NA | NA | NA | NA | NA |
| Salkhit skullcap - B | Shotgun | 7.35E+9 | 1,753,089 | 1,088,188 | 7,120 | 0.65 | 7,094 | 1 | 56.6 | 1.68E+9 | 32 | 22.3 | 1,442 | 52.9 | NA | NA | NA | NA | NA |
| Salkhit skullcap - C | Shotgun | 5.78E+9 | 1,824,021 | 1,113,871 | 8,841 | 0.79 | 8,820 | 1 | 55 | 1.54E+9 | 38.7 | 23.9 | 1,933 | 54.5 | NA | NA | NA | NA | NA |
| Extraction Negative Control | Shotgun | 9.41E+7 | 253,965 | 3,034 | 64 | 2.10 | 63 | 1.02 | 49.3 | 1.15E+6 | 0 | 0 | NA | NA | NA | NA | NA | NA | NA |
| Bleached - Salkhit skullcap - A | Shotgun | 1.22E+9 | 3,481,774 | 1,569,481 | 88,480 | 5.63 | 87,942 | 1.01 | 55.9 | 1.72E+9 | 39.4 | 25.6 | 20,935 | 54.4 | NA | NA | NA | NA | NA |
| Bleached - Salkhit skullcap - B | Shotgun | 1.10E+9 | 4,759,556 | 2,120,461 | 43,421 | 2.05 | 43,140 | 1.01 | 54.8 | 5.46E+8 | 40 | 25.7 | 10,423 | 53 | NA | NA | NA | NA | NA |
| Bleached - Salkhit skullcap - C | Shotgun | 1.39E+9 | 5,187,312 | 2,419,823 | 55,457 | 2.29 | 55,147 | 1.01 | 56.2 | 8.3E+8 | 36.4 | 24.4 | 12,223 | 53.9 | NA | NA | NA | NA | NA |
| Bleached Extraction Negative Control | Shotgun | 8.46E+7 | 408,438 | 3,282 | 216 | 6.58 | 214 | 1.01 | 60.3 | 2.67E+6 | 0 | 2.5 | 1 | 53 | NA | NA | NA | NA | NA |
| Salkhit skullcap - A | Panel 1 - 390k | NA | 37,041,716 | 28,475,353 | 11,417,875 | 40.1 | 1,788,184 | 6.39 | 69.4 | - | 19.3 | 10.9 | 197,866 | 62.3 | 393,788 | 300,018 | 76.19 | 39,282 | 9.97 |
| Salkhit skullcap - B | Panel 1 - 390k | NA | 38,262,085 | 29,306,367 | 8,280,795 | 28.26 | 741,599 | 11.17 | 65 | - | 27.6 | 16.7 | 130,103 | 60.7 | 393,788 | 162,285 | 41.21 | 22,553 | 5.73 |
| Salkhit skullcap - C | Panel 1 - 390k | NA | 32,365,945 | 24,629,981 | 7,303,418 | 29.65 | 687,176 | 10.63 | 63.1 | - | 32.7 | 20.1 | 143,623 | 61.1 | 393,788 | 149,110 | 37.86 | 23,922 | 6.07 |
| Extraction Negative Control | Panel 1 - 390k | NA | 1,381,368 | 621,748 | 430,832 | 69.29 | 6,571 | 65.57 | 67.1 | - | 5.5 | 3.9 | 189 | 62.2 | 393,788 | 2,181 | 0.55 | 114 | 0.03 |
| Bleached - Salkhit skullcap - A | Panel 1 - 390k | NA | 39,250,483 | 28,849,269 | 15,246,146 | 52.85 | 2,420,793 | 6.3 | 61.7 | - | 36.1 | 23.8 | 583,264 | 60.7 | 393,788 | 280,121 | 71.13 | 71,127 | 18.06 |
| Bleached - Salkhit skullcap - B | Panel 1 - 390k | NA | 41,188,066 | 29,092,265 | 11,733,498 | 40.33 | 1,026,394 | 11.43 | 60.3 | - | 37 | 23.9 | 246,475 | 58.4 | 393,788 | 154,514 | 39.24 | 28,141 | 7.15 |
| Bleached Extraction Negative Control | Panel 1 - 390k | NA | 40,812,151 | 18,269,941 | 13,060,607 | 71.49 | 57,527 | 227.03 | 64.4 | - | 6.8 | 4.7 | 1916 | 61.6 | 393,788 | 12,952 | 3.3 | 1,082 | 0.27 |
| Salkhit skullcap - A | Panel 2 - 840k | NA | 43,518,390 | 33,832,643 | 11,735,703 | 34.69 | 3,171,259 | 3.7 | 70.5 | - | 18.7 | 10.6 | 347,093 | 63.9 | 842,630 | 526,596 | 62.49 | 59,231 | 7.03 |
| Salkhit skullcap - B | Panel 2 - 840k | NA | 41,631,541 | 32,163,728 | 7,251,994 | 22.55 | 1,253,164 | 5.79 | 66.5 | - | 26.9 | 16.1 | 215,878 | 62.3 | 842,630 | 259,710 | 30.82 | 33,515 | 3.98 |
| Salkhit skullcap - C | Panel 2 - 840k | NA | 35,115,205 | 26,748,990 | 6,375,569 | 23.83 | 1,322,570 | 4.82 | 64.6 | - | 32 | 19.5 | 274,530 | 62.6 | 842,630 | 261,749 | 31.06 | 40,391 | 4.8 |
| Extraction Negative Control | Panel 2 - 840k | NA | 3,865,759 | 1,646,987 | 894,744 | 54.33 | 14,124 | 63.35 | 65.8 | - | 14 | 7.4 | 1238 | 63.6 | 842,630 | 6,081 | 0.72 | 605 | 0.07 |
| Bleached - Salkhit skullcap - A | Panel 2 - 840k | NA | 40,865,609 | 29,921,771 | 13,488,103 | 45.08 | 3,596,302 | 3.75 | 63.4 | - | 35.7 | 23.3 | 865,430 | 62.4 | 842,630 | 505,368 | 59.97 | 117,688 | 13.97 |
| Bleached - Salkhit skullcap - B | Panel 2 - 840k | NA | 47,381,399 | 33,344,588 | 10,861,631 | 32.57 | 1,590,315 | 6.83 | 62 | - | 36.2 | 23.4 | 379,459 | 60.1 | 842,630 | 276,248 | 32.78 | 48,829 | 5.79 |
| Bleached Extraction Negative Control | Panel 2 - 840k | NA | 48,353,676 | 21,378,677 | 13,042,556 | 61.01 | 72,378 | 180.2 | 64.9 | - | 6.8 | 4.4 | 2347 | 63.1 | 842,630 | 13,397 | 1.58 | 1,003 | 0.12 |
| Salkhit skullcap - A | Panel 3 - Big yoruba/Altai | NA | 40,141,919 | 18,073,023 | 7,935,790 | 43.91 | 1,809,720 | 4.39 | 78.4 | - | 16.7 | 9.5 | 175,646 | 72 | 997,780 | 438,435 | 43.94 | 32,472 | 3.25 |
| Salkhit skullcap - B | Panel 3 - Big yoruba/Altai | NA | 40,794,331 | 18,057,777 | 4,490,358 | 24.87 | 842,718 | 5.33 | 73.5 | - | 25.3 | 15 | 135,477 | 69.6 | 997,780 | 213,665 | 21.41 | 22,758 | 2.28 |
| Salkhit skullcap - C | Panel 3 - Big yoruba/Altai | NA | 35,049,085 | 14,473,877 | 4,084,159 | 28.22 | 761,055 | 5.37 | 72.5 | - | 30.1 | 18.8 | 150,948 | 70.3 | 997,780 | 191,291 | 19.17 | 24,132 | 2.42 |
| Extraction Negative Control | Panel 3 - Big yoruba/Altai | NA | 2,649,767 | 721,805 | 475,497 | 65.88 | 8,805 | 54 | 73.9 | - | 6.7 | 3.9 | 285 | 70.6 | 997,780 | 2,473 | 0.25 | 116 | 0.01 |
| Bleached - Salkhit skullcap - A | Panel 3 - Big yoruba/Altai | NA | 44,646,122 | 18,407,167 | 10,929,442 | 59.38 | 2,449,998 | 4.46 | 73.1 | - | 34.1 | 23.1 | 570,874 | 71.9 | 997,780 | 455,510 | 45.65 | 82,683 | 8.29 |
| Bleached - Salkhit skullcap - B | Panel 3 - Big yoruba/Altai | NA | 46,000,138 | 17,617,053 | 7,324,079 | 41.57 | 950,859 | 7.7 | 71.8 | - | 33.5 | 22.4 | 212,920 | 69.8 | 997,780 | 213,969 | 21.44 | 30,395 | 3.05 |
| Bleached - Salkhit skullcap - C | Panel 3 - Big yoruba/Altai | NA | 41,761,119 | 16,876,757 | 7,950,837 | 47.11 | 1,292,989 | 6.15 | 73.7 | - | 31 | 20.1 | 257,508 | 71.3 | 997,780 | 293,164 | 29.38 | 39,544 | 3.96 |
| Bleached Extraction Negative Control | Panel 3 - Big yoruba/Altai | NA | 3,081,961 | 1,014,409 | 659,429 | 65.01 | 18,307 | 36.02 | 74 | - | 3 | 1.4 | 209 | 72.4 | 997,780 | 3,769 | 0.38 | 65 | 0.01 |
| Salkhit skullcap - A | Panel 4 - Archaic Admixture | NA | 56,865,111 | 43,931,204 | 10,977,528 | 24.99 | 4,473,960 | 2.45 | 68.4 | - | 18.6 | 11 | 501,166 | 61.3 | 1,749,385 | 1,050,397 | 60.04 | 13,5542 | 7.75 |
| Salkhit skullcap - B | Panel 4 - Archaic Admixture | NA | 60,658,434 | 45,965,132 | 5,751,294 | 12.51 | 1,887,854 | 3.05 | 63.9 | - | 25.6 | 16.4 | 324,303 | 59.5 | 1,749,385 | 594,643 | 33.99 | 83,554 | 4.78 |
| Salkhit skullcap - C | Panel 4 - Archaic Admixture | NA | 56,051,712 | 41,664,007 | 5,886,274 | 14.13 | 1,883,455 | 3.13 | 61.9 | - | 31.8 | 19.6 | 387,440 | 59.5 | 1,749,385 | 585,170 | 33.45 | 98,434 | 5.63 |
| Bleached - Salkhit skullcap - A | Panel 4 - Archaic Admixture | NA | 65,298,294 | 46,016,340 | 18,631,792 | 40.49 | 4,386,526 | 4.25 | 61.5 | - | 35.9 | 23.8 | 1,056,376 | 60 | 1,749,385 | 912,645 | 52.17 | 200,323 | 11.45 |
| Bleached - Salkhit skullcap - B | Panel 4 - Archaic Admixture | NA | 68,815,447 | 45,527,528 | 10,283,638 | 22.59 | 1,871,786 | 5.49 | 59.5 | - | 36.3 | 23.5 | 442,868 | 57.1 | 1,749,385 | 471,416 | 26.95 | 80,948 | 4.63 |
| Bleached - Salkhit skullcap - C | Panel 4 - Archaic Admixture | NA | 59,509,257 | 41,040,207 | 11,026,555 | 26.87 | 2,279,450 | 4.84 | 61.9 | - | 33.5 | 21.4 | 485,948 | 58.9 | 1,749,385 | 596,647 | 34.11 | 98,271 | 5.68 |
| Bleached Extraction Negative Control | Panel 4 - Archaic Admixture | NA | 4,281,363 | 1,785,886 | 930,953 | 52.13 | 29,067 | 32.03 | 66.2 | - | 2.5 | 2 | 314 | 64.4 | 1,749,385 | 6,709 | 0.38 | 89 | 0.01 |

**Table S2. DNA libraries prepared from the Salkhit specimen.**

The total number of DNA molecules incorporated in the library is estimated by qPCR and is used to estimate the proportion of genome covered per library content. The %5' and %3' C to T represent the frequency of cytosine (C) to thymine (T) mismatches to the reference at the first and last base of each sequence.

*Sequences merged and filtered for mapping quality 25 and minimal size of 35 bases. Seq=Sequences, nbr = number, deam = sequences showing evidence for deamination (C to T mismatch) at the first or last base, NA = Not Applicable.

| Library ID | Sample | Shotgun - Nuclear SNPs Capture | % 5' C to T | % 5' C to T 95CI | cond. % 5' C to T | cond. % 5' C to T 95CI | % 3' C to T | % 3' C to T 95CI | cond. % 3' C to T | cond. % 3' C to T 95CI | diff. % 5' C to T and cond. | diff. % 3' C to T and cond. | Contamination estimate cond. 5' | Contamination estimate cond. 3' | av. Contamination estimate |
|---|---|---|---|---|---|---|---|---|---|---|---|---|---|---|---|
| F8405 | Salkhit skullcap - A | Shotgun | 22.8 | 21.4-24.1 | 38.3 | 32.0-45.1 | 13.1 | 11.9-14.4 | 22.8 | 18.0-28.5 | 15.5 | 9.7 | 0.40 | 0.43 | 0.42 |
| F8407 | Salkhit skullcap - B | Shotgun | 32 | 29.8-34.2 | 35.7 | 28.0-42.9 | 22.3 | 20.1-24.6 | 30.4 | 23.9-37.9 | 3.7 | 8.1 | 0.10 | 0.27 | 0.19 |
| F8406 | Salkhit skullcap - C | Shotgun | 38.7 | 36.6-40.8 | 48.5 | 41.2-54.9 | 23.9 | 21.9-25.9 | 27.6 | 22.6-33.3 | 9.8 | 3.7 | 0.20 | 0.13 | 0.17 |
| F8409 | Extraction Negative Control | Shotgun | 0 | 0.0-27.8 | NA | NA | 0 | 0.0-21.5 | NA | NA | NA | NA | NA | NA | NA |
| G2740 | Bleached - Salkhit skullcap - A | Shotgun | 39.4 | 38.7-40.0 | 42.5 | 40.5-44.6 | 25.6 | 24.9-26.3 | 26.2 | 24.5-27.9 | 3.1 | 0.6 | 0.07 | 0.02 | 0.05 |
| G2742 | Bleached - Salkhit skullcap - B | Shotgun | 40 | 39.1-41.0 | 46.5 | 43.6-49.2 | 25.7 | 24.7-26.6 | 29.5 | 27.1-32.0 | 6.5 | 3.8 | 0.14 | 0.13 | 0.13 |
| G2741 | Bleached - Salkhit skullcap - C | Shotgun | 36.4 | 35.5-37.2 | 41.2 | 38.7-43.7 | 24.4 | 23.6-25.2 | 28.4 | 26.2-30.8 | 4.8 | 4 | 0.12 | 0.14 | 0.13 |
| G2743 | Bleached Extraction Negative Control | Shotgun | 0 | 0.0-8.8 | NA | NA | 2.5 | 0.4-12.9 | 0 | 0 | NA | NA | NA | NA | NA |
| G5754 | Salkhit skullcap - A | Panel 1 - 390k | 19.3 | 19.2-19.4 | 35.8 | 35.2-36.4 | 10.9 | 10.8-11.0 | 22.1 | 21.6-22.6 | 16.5 | 11.2 | 0.46 | 0.51 | 0.48 |
| G5756 | Salkhit skullcap - B | Panel 1 - 390k | 27.6 | 27.4-27.8 | 40.2 | 39.4-41.0 | 16.7 | 16.5-16.9 | 25 | 24.3-25.6 | 12.6 | 8.3 | 0.31 | 0.33 | 0.32 |
| G5755 | Salkhit skullcap - C | Panel 1 - 390k | 32.7 | 32.5-32.9 | 39.7 | 39.0-40.5 | 20.1 | 19.9-20.3 | 24.5 | 23.9-25.1 | 7 | 4.4 | 0.18 | 0.18 | 0.18 |
| G5757 | Extraction Negative Control | Panel 1 - 390k | 5.5 | 4.6-6.5 | 18.8 | 4.0-45.6 | 3.9 | 3.0-4.8 | 15 | 3.2-37.9 | 13.3 | 11.1 | 0.71 | 0.74 | 0.72 |
| G5750 | Bleached - Salkhit skullcap - A | Panel 1 - 390k | 36.1 | 36.0-36.3 | 38.3 | 37.9-38.6 | 23.8 | 23.7-24.0 | 25.1 | 24.8-25.4 | 2.2 | 1.3 | 0.06 | 0.05 | 0.05 |
| G5752 | Bleached - Salkhit skullcap - B | Panel 1 - 390k | 37 | 36.9-37.2 | 42.5 | 42.0-43.1 | 23.9 | 23.8-24.1 | 27.6 | 27.1-28.1 | 5.5 | 3.7 | 0.13 | 0.13 | 0.13 |
| G5753 | Bleached Extraction Negative Control | Panel 1 - 390k | 6.8 | 6.5-7.2 | 20.9 | 15.2-27.6 | 4.7 | 4.3-5.0 | 14.7 | 10.6-19.7 | 14.1 | 10 | 0.67 | 0.68 | 0.68 |
| G5762 | Salkhit skullcap - A | Panel 2 - 840k | 18.7 | 18.7-18.8 | 35.3 | 34.8-35.7 | 10.6 | 10.6-10.7 | 21.9 | 21.5-22.3 | 16.6 | 11.3 | 0.47 | 0.52 | 0.49 |
| G5764 | Salkhit skullcap - B | Panel 2 - 840k | 26.9 | 26.7-27.0 | 39.3 | 38.7-39.9 | 16.1 | 16.0-16.2 | 23.8 | 23.3-24.3 | 12.4 | 7.7 | 0.32 | 0.32 | 0.32 |
| G5763 | Salkhit skullcap - C | Panel 2 - 840k | 32 | 31.8-32.1 | 38.7 | 38.2-39.3 | 19.5 | 19.4-19.7 | 24.2 | 23.8-24.6 | 6.7 | 4.7 | 0.17 | 0.19 | 0.18 |
| G5765 | Extraction Negative Control | Panel 2 - 840k | 14 | 13.2-14.9 | 37.4 | 28.5-46.9 | 7.4 | 6.7-8.1 | 92.4 | 90.4-93.9 | 23.4 | 85 | 0.63 | 0.92 | 0.77 |
| G5758 | Bleached - Salkhit skullcap - A | Panel 2 - 840k | 35.7 | 35.6-35.8 | 37.8 | 37.6-38.1 | 23.3 | 23.2-23.4 | 24.6 | 24.4-24.9 | 2.1 | 1.3 | 0.06 | 0.05 | 0.05 |
| G5760 | Bleached - Salkhit skullcap - B | Panel 2 - 840k | 36.2 | 36.0-36.3 | 42.3 | 41.8-42.7 | 23.4 | 23.3-23.6 | 27.4 | 27.0-27.8 | 6.1 | 4 | 0.14 | 0.15 | 0.15 |
| G5761 | Bleached Extraction Negative Control | Panel 2 - 840k | 6.8 | 6.4-7.1 | 26 | 20.4-32.3 | 4.4 | 4.1-4.7 | 17.2 | 13.1-21.4 | 19.2 | 12.8 | 0.74 | 0.74 | 0.74 |
| G5746 | Salkhit skullcap - A | Panel 3 - Big yoruba/Altai | 16.7 | 16.6-16.8 | 34.5 | 33.8-35.1 | 9.5 | 9.4-9.6 | 21.2 | 20.7-21.7 | 17.8 | 11.7 | 0.52 | 0.55 | 0.53 |
| G5748 | Salkhit skullcap - B | Panel 3 - Big yoruba/Altai | 25.3 | 25.1-25.5 | 38.8 | 38.0-39.5 | 15 | 14.8-15.1 | 23.6 | 23.0-24.2 | 13.5 | 8.6 | 0.35 | 0.36 | 0.36 |
| G5747 | Salkhit skullcap - C | Panel 3 - Big yoruba/Altai | 30.1 | 29.9-30.3 | 38.2 | 37.4-38.9 | 18.8 | 18.6-19.0 | 24.3 | 23.7-24.9 | 8.1 | 5.5 | 0.21 | 0.23 | 0.22 |
| G5749 | Extraction Negative Control | Panel 3 - Big yoruba/Altai | 6.7 | 5.9-7.7 | 30.4 | 13.2-52.9 | 3.9 | 3.2-4.7 | 20 | 8.4-36.9 | 23.7 | 16.1 | 0.78 | 0.81 | 0.79 |
| G5742 | Bleached - Salkhit skullcap - A | Panel 3 - Big yoruba/Altai | 34.1 | 34.0-34.2 | 36.3 | 36.0-36.7 | 23.1 | 23.0-23.2 | 24.4 | 24.1-24.7 | 2.2 | 1.3 | 0.06 | 0.05 | 0.06 |
| G5744 | Bleached - Salkhit skullcap - B | Panel 3 - Big yoruba/Altai | 33.5 | 33.3-33.7 | 41 | 40.4-41.6 | 22.4 | 22.3-22.6 | 28.1 | 27.6-28.7 | 7.5 | 5.7 | 0.18 | 0.20 | 0.19 |
| G5743 | Bleached - Salkhit skullcap - C | Panel 3 - Big yoruba/Altai | 31 | 30.8-31.1 | 38.8 | 38.3-39.4 | 20.1 | 20.0-20.3 | 25.9 | 25.4-26.3 | 7.8 | 5.8 | 0.20 | 0.22 | 0.21 |
| G5745 | Bleached Extraction Negative Control | Panel 3 - Big yoruba/Altai | 3 | 2.6-3.5 | 13.3 | 0.2-31.9 | 1.4 | 1.0-1.8 | 7.1 | 0.1-18.3 | 10.3 | 5.7 | 0.77 | 0.80 | 0.79 |
| G3549 | Salkhit skullcap - A | Panel 4 - Archaic Admixture | 18.6 | 18.6-18.7 | 34.6 | 34.2-35.0 | 11 | 10.9-11.0 | 21.8 | 21.5-22.1 | 16 | 10.8 | 0.46 | 0.50 | 0.48 |
| G3551 | Salkhit skullcap - B | Panel 4 - Archaic Admixture | 25.6 | 25.5-25.7 | 38.4 | 37.9-38.8 | 16.4 | 16.3-16.6 | 24.8 | 24.4-25.2 | 12.8 | 8.4 | 0.33 | 0.34 | 0.34 |
| G3550 | Salkhit skullcap - C | Panel 4 - Archaic Admixture | 31.8 | 31.6-31.9 | 38.7 | 38.3-39.2 | 19.6 | 19.5-19.7 | 24.5 | 24.1-24.8 | 6.9 | 4.9 | 0.18 | 0.20 | 0.19 |
| G5736 | Bleached - Salkhit skullcap - A | Panel 4 - Archaic Admixture | 35.9 | 35.8-36.0 | 38 | 37.7-38.2 | 23.8 | 23.7-23.9 | 25 | 24.8-25.2 | 2.1 | 1.2 | 0.06 | 0.05 | 0.05 |
| G5738 | Bleached - Salkhit skullcap - B | Panel 4 - Archaic Admixture | 36.3 | 36.2-36.4 | 42.4 | 42.0-42.8 | 23.5 | 23.4-23.6 | 27.8 | 27.4-28.1 | 6.1 | 4.3 | 0.14 | 0.15 | 0.15 |
| G5737 | Bleached - Salkhit skullcap - C | Panel 4 - Archaic Admixture | 33.5 | 33.4-33.7 | 40.7 | 40.3-41.1 | 21.4 | 21.3-21.6 | 26.2 | 25.9-26.6 | 7.2 | 4.8 | 0.18 | 0.18 | 0.18 |
| G5739 | Bleached Extraction Negative Control | Panel 4 - Archaic Admixture | 2.5 | 2.1-2.8 | 5.6 | 0.7-18.7 | 2 | 1.6-2.4 | 4.3 | 0.1-11.5 | 3.1 | 2.3 | 0.55 | 0.53 | 0.54 |

**Table S3. Frequencies of terminal C to T mismatches (5’ and 3’) to the human genome and contamination estimates.**
*Sequences merged and filtered for mapping quality 25 and minimal size of 35 bases.
Seq=Sequences, deam = sequences showing evidence for deamination at the first or last base, NA = Not Applicable, Cond. = Conditional C to T mismatches frequency, Av = Average.

| | mtDNA | | Cond. C to T mismatch | | | AdmixFrog | |
|---|---|---|---|---|---|---|---|
| | All | deam. only | 5' end | 3' end | Average | All | deam. only |
| **Salkhit skullcap - A** | 0.42 | 0.03 | 0.48 | 0.52 | 0.50 | 0.47 | 0.03 |
| **Salkhit skullcap - B** | 0.35 | 0.01 | 0.34 | 0.34 | 0.34 | 0.25 | 0.03 |
| **Salkhit skullcap - C** | 0.19 | 0.04 | 0.18 | 0.2 | 0.19 | 0.15 | 0.02 |
| **Bleached - Salkhit skullcap - A** | NA | NA | 0.06 | 0.05 | 0.05 | 0.02 | 0.01 |
| **Bleached - Salkhit skullcap - B** | NA | NA | 0.14 | 0.15 | 0.15 | 0.20 | 0.01 |
| **Bleached - Salkhit skullcap - C** | NA | NA | 0.19 | 0.21 | 0.20 | 0.23 | 0.02 |

**Table S4. Contamination estimates for the Salkhit DNA libraries.**
All= All mapped sequences, deam = sequences showing evidence for deamination at the first or last base, NA = Not Applicable, Cond. = Conditional C to T mismatches frequency.

| Sample | Nuclear SNPs Capture | Total nbr of seq. generated | Total nbr of seq. merged and filtered* | Total mapped seq. (MAPQ 25) | % mapped seq. | Unique mapped seq. | Duplication rate | % 5' C to T | % 3' C to T | Total nbr of deaminated seq. | average size of deam. reads | Contamination estimate cond. 5' | Contamination estimate cond. 3' | Total nbr of targetted SNPs | Nbr of SNPs covered (all reads) | % of targetted SNPs covered (all reads) | Nbr of SNPs covered (deam reads only) | % of targetted SNPs covered (deam reads only) |
|---|---|---|---|---|---|---|---|---|---|---|---|---|---|---|---|---|---|---|
| Salkhit_all | Merged - Panel 1, 2, 3 - 2.2 million SNPs | 6.77E+8 | 4.17E+8 | 1.51E+8 | 36.43 | 22,369,743 | 6.79 | 29.7% | 18.4% | 2,478,541 | 63.4 | 21.84% | 25.51% | 2,144,502 | 1,877,094 | 87.53% | 603,300 | 28.13% |
| Salkhit_Bl_all | Merged - Panel 1, 2, 3 - 2.2 million SNPs | 3.19E+8 | 1.83E+8 | 8.08E+7 | 44.17 | 11,288,144 | 7.16 | 35.3% | 23.2% | 1,538,622 | 63.1 | 8.55% | 9.38% | 2,144,502 | 1,543,474 | 71.97% | 395,035 | 18.42% |
| Salkhit_Bl_A** | Merged - Panel 1, 2, 3 - 2.2 million SNPs | 1.32E+8 | 8.12E+7 | 4.14E+7 | 51.00 | 7,053,801 | 5.87 | 35.7% | 23.6% | 988,449 | 63 | 4.03% | 4.84% | 2,144,502 | 1,219,892 | 56.88% | 271,137 | 12.64% |
| Salkhit_all | Panel 4 - Archaic Admixture | 3.95E+8 | 2.83E+8 | 6.62E+7 | 23.38 | 17,385,083 | 3.81 | 29.5% | 18.6% | 1,951,273 | 59.5 | 22.6% | 25.3% | 1,749,385 | 1,498,291 | 85.65% | 543,742 | 31.08% |
| Salkhit_Bl_all | Panel 4 - Archaic Admixture | 2,03E+8 | 1.38E+8 | 4.12E+7 | 29.93 | 8,702,369 | 4.74 | 35.3% | 23.1% | 1,189,841 | 59 | 9.9% | 10.8% | 1,749,385 | 1,253,626 | 71.66% | 343,537 | 19.64% |
| Salkhit_Bl_A** | Panel 4 - Archaic Admixture | 6.92E+7 | 4.81E+7 | 1.93E+7 | 40.20 | 4,490,908 | 4.31 | 35.8% | 23.8% | 637,429 | 60 | 4.3% | 4.8% | 1,749,385 | 912,645 | 52.17% | 200,323 | 11.45% |

**Table S5. Salkhit data sets.**
*Sequences merged and filtered for mapping quality 25 and minimal size of 35 bases.
**Library made from bleached extract A.
Bl=Bleached, deam = sequences showing evidence for deamination at the first or last base, NA = Not Applicable, Cond. = Conditional C to T mismatches frequency.

| Pop1 | Pop2 | Pop3 | Outgroup | Dscore | Zscore | std.err | nbr of BABA sites | nbr of ABBA sites | nbr of SNPs present in the all dataset | Pop3 geographic origin |
|---|---|---|---|---|---|---|---|---|---|---|
| Salkhit_all | Salkhit_all_deam | Ami | Mbuti | 0.0008 | 0.18 | 0.0042 | 12,617 | 12,597 | 553,564 | East Eurasia |
| Salkhit_all | Salkhit_all_deam | Han | Mbuti | 0.001 | 0.24 | 0.0040 | 12,608 | 12,583 | 553,606 | East Eurasia |
| Salkhit_all | Salkhit_all_deam | Dai | Mbuti | 0.001 | 0.27 | 0.0036 | 12,561 | 12,536 | 553,608 | East Eurasia |
| Salkhit_all | Salkhit_all_deam | Korean | Mbuti | 0.0053 | 1.25 | 0.0042 | 12,664 | 12,530 | 553,571 | East Eurasia |
| Salkhit_all | Salkhit_all_deam | Japanese | Mbuti | 0.0034 | 0.83 | 0.0040 | 12,623 | 12,536 | 553,600 | East Eurasia |
| Salkhit_all | Salkhit_all_deam | Atayal | Mbuti | -0.0007 | -0.15 | 0.0045 | 12,561 | 12,580 | 553,364 | Southeast Asia |
| Salkhit_all | Salkhit_all_deam | Thai | Mbuti | 0.001 | 0.23 | 0.0042 | 12,543 | 12,519 | 553,579 | Southeast Asia |
| Salkhit_all | Salkhit_all_deam | Cambodian | Mbuti | -0.0006 | -0.13 | 0.0044 | 12,504 | 12,519 | 553,584 | Southeast Asia |
| Salkhit_all | Salkhit_all_deam | Kinh | Mbuti | 0.004 | 0.97 | 0.0041 | 12,621 | 12,521 | 553,564 | Southeast Asia |
| Salkhit_all | Salkhit_all_deam | Igorot | Mbuti | 0.0018 | 0.40 | 0.0044 | 12,583 | 12,538 | 553,554 | Southeast Asia |
| Salkhit_all | Salkhit_all_deam | Hawaiian | Mbuti | -0.0026 | -0.57 | 0.0045 | 12,532 | 12,598 | 553,339 | Oceania |
| Salkhit_all | Salkhit_all_deam | Australian | Mbuti | -0.0083 | -1.96 | 0.0042 | 12,266 | 12,471 | 553,594 | Oceania |
| **Salkhit_all** | **Salkhit_all_deam** | **Papuan** | **Mbuti** | **-0.0141** | **-3.45** | **0.0040** | **12,198** | **12,546** | **553,609** | **Oceania** |
| Salkhit_all | Salkhit_all_deam | Bougainville | Mbuti | -0.0079 | -1.76 | 0.0044 | 12,287 | 12,482 | 553,556 | Oceania |
| **Salkhit_all** | **Salkhit_all_deam** | **Maori** | **Mbuti** | **0.0189** | **4.08** | **0.0046** | **12,627** | **12,159** | **553,338** | **Oceania** |
| Salkhit_all | Salkhit_all_deam | Ulchi | Mbuti | 0.0075 | 1.84 | 0.0040 | 12,706 | 12,516 | 553,571 | Siberia |
| Salkhit_all | Salkhit_all_deam | Yakut | Mbuti | 0.006 | 1.47 | 0.0040 | 12,675 | 12,523 | 553,573 | Siberia |
| Salkhit_all | Salkhit_all_deam | Itelman | Mbuti | 0.0069 | 1.42 | 0.0048 | 12,692 | 12,518 | 553,423 | Siberia |
| **Salkhit_all** | **Salkhit_all_deam** | **Aleut** | **Mbuti** | **0.0135** | **3.07** | **0.0043** | **12,702** | **12,364** | **553,578** | **Siberia** |
| **Salkhit_all** | **Salkhit_all_deam** | **Chukchi** | **Mbuti** | **0.0164** | **3.43** | **0.0047** | **12,756** | **12,344** | **553,404** | **Siberia** |
| **Salkhit_all** | **Salkhit_all_deam** | **Altaian** | **Mbuti** | **0.0141** | **3.04** | **0.0046** | **12,707** | **12,354** | **553,367** | **Central Eurasia** |
| **Salkhit_all** | **Salkhit_all_deam** | **Kyrgyz** | **Mbuti** | **0.0123** | **3.04** | **0.0040** | **12,729** | **12,419** | **553,590** | **Central Eurasia** |
| **Salkhit_all** | **Salkhit_all_deam** | **Iranian** | **Mbuti** | **0.0234** | **5.86** | **0.0039** | **12,633** | **12,056** | **553,579** | **Central Eurasia** |
| Salkhit_all | Salkhit_all_deam | Mala | Mbuti | 0.0053 | 1.37 | 0.0038 | 12,449 | 12,317 | 553,605 | Central Eurasia |
| **Salkhit_all** | **Salkhit_all_deam** | **Russian** | **Mbuti** | **0.0283** | **6.71** | **0.0042** | **12,839** | **12,132** | **553,574** | **West Eurasia** |
| **Salkhit_all** | **Salkhit_all_deam** | **Czech** | **Mbuti** | **0.0302** | **6.39** | **0.0047** | **12,875** | **12,119** | **553,403** | **West Eurasia** |
| **Salkhit_all** | **Salkhit_all_deam** | **Saami** | **Mbuti** | **0.0262** | **5.98** | **0.0043** | **12,900** | **12,240** | **553,594** | **West Eurasia** |
| **Salkhit_all** | **Salkhit_all_deam** | **French** | **Mbuti** | **0.0333** | **8.03** | **0.0041** | **12,881** | **12,052** | **553,604** | **West Eurasia** |
| **Salkhit_all** | **Salkhit_all_deam** | **Spanish** | **Mbuti** | **0.0338** | **7.96** | **0.0042** | **12,838** | **11,998** | **553,582** | **West Eurasia** |
| **Salkhit_all** | **Salkhit_all_deam** | **Sardinian** | **Mbuti** | **0.0366** | **9.09** | **0.0040** | **12,853** | **11,945** | **553,599** | **West Eurasia** |
| Salkhit_all | Salkhit_all_deam | Mayan | Mbuti | 0.0021 | 0.5 | 0.0042 | 12,685 | 12,631 | 553,559 | America |
| Salkhit_all | Salkhit_all_deam | Quechua | Mbuti | 0.0027 | 0.64 | 0.0042 | 12,639 | 12,570 | 553,596 | America |
| **Salkhit_all** | **Salkhit_all_deam** | **Cree** | **Mbuti** | **0.0144** | **3.45** | **0.0041** | **12,732** | **12,371** | **553,576** | **America** |
| Salkhit_all | Salkhit_all_deam | Karitiana | Mbuti | 0.0061 | 1.37 | 0.0044 | 12,746 | 12,590 | 553,588 | America |
| Salkhit_all | Salkhit_all_deam | Mixe | Mbuti | 0.0048 | 1.1 | 0.0043 | 12,660 | 12,540 | 553,584 | America |

**Table S6A.** *D-statistic* comparing the genetic similarity of present-day human populations to the set of all Salkhit sequences "Salkhit_all" and the set of deaminated sequences only "Salkhit_all_deam". Significant D-statistics (|Zscore| > 2) are underline in red.

| Pop1 | Pop2 | Pop3 | Outgroup | Dscore | Zscore | std.err | nbr of BABA sites | nbr of ABBA sites | nbr of SNPs present in the all dataset | Pop3 geographic origin |
|---|---|---|---|---|---|---|---|---|---|---|
| Salkhit_bl_all | Salkhit_bl_all_deam | Ami | Mbuti | -0.0006 | -0.1 | 0.0060 | 4,601 | 4,607 | 360,359 | East Eurasia |
| Salkhit_bl_all | Salkhit_bl_all_deam | Han | Mbuti | 0.0107 | 1.8 | 0.0059 | 4,623 | 4,524 | 360,389 | East Eurasia |
| Salkhit_bl_all | Salkhit_bl_all_deam | Dai | Mbuti | 0.0109 | 1.85 | 0.0059 | 4,607 | 4,507 | 360,392 | East Eurasia |
| Salkhit_bl_all | Salkhit_bl_all_deam | Korean | Mbuti | 0.01 | 1.60 | 0.0062 | 4,603 | 4,512 | 360,364 | East Eurasia |
| Salkhit_bl_all | Salkhit_bl_all_deam | Japanese | Mbuti | 0.0066 | 1.06 | 0.0061 | 4,598 | 4,537 | 360,387 | East Eurasia |
| Salkhit_bl_all | Salkhit_bl_all_deam | Atayal | Mbuti | -0.0041 | -0.54 | 0.0076 | 4,551 | 4,588 | 360,225 | Southeast Asia |
| Salkhit_bl_all | Salkhit_bl_all_deam | Thai | Mbuti | 0.0091 | 1.40 | 0.0064 | 4,600 | 4,517 | 360,369 | Southeast Asia |
| Salkhit_bl_all | Salkhit_bl_all_deam | Cambodian | Mbuti | 0.0094 | 1.47 | 0.0063 | 4,604 | 4,518 | 360,374 | Southeast Asia |
| Salkhit_bl_all | Salkhit_bl_all_deam | Kinh | Mbuti | 0.0095 | 1.49 | 0.0063 | 4,602 | 4,515 | 360,362 | Southeast Asia |
| Salkhit_bl_all | Salkhit_bl_all_deam | Igorot | Mbuti | 0.0088 | 1.35 | 0.0065 | 4,618 | 4,538 | 360,358 | Southeast Asia |
| Salkhit_bl_all | Salkhit_bl_all_deam | Hawaiian | Mbuti | 0.0078 | 1.1 | 0.0070 | 4,596 | 4,525 | 360,223 | Oceania |
| Salkhit_bl_all | Salkhit_bl_all_deam | Australian | Mbuti | -0.004 | -0.60 | 0.0066 | 4,460 | 4,496 | 360,383 | Oceania |
| Salkhit_bl_all | Salkhit_bl_all_deam | Papuan | Mbuti | -0.0051 | -0.79 | 0.0064 | 4,494 | 4,539 | 360,394 | Oceania |
| Salkhit_bl_all | Salkhit_bl_all_deam | Bougainville | Mbuti | -0.0066 | -0.98 | 0.0067 | 4,462 | 4,522 | 360,355 | Oceania |
| Salkhit_bl_all | Salkhit_bl_all_deam | Maori | Mbuti | 0.0027 | 0.39 | 0.0069 | 4,511 | 4,487 | 360,203 | Oceania |
| **Salkhit_bl_all** | **Salkhit_bl_all_deam** | **Ulchi** | **Mbuti** | **0.0133** | **2.08** | **0.0064** | **4,640** | **4,518** | **360,366** | **Siberia** |
| **Salkhit_bl_all** | **Salkhit_bl_all_deam** | **Yakut** | **Mbuti** | **0.0133** | **2.07** | **0.0064** | **4,635** | **4,514** | **360,367** | **Siberia** |
| **Salkhit_bl_all** | **Salkhit_bl_all_deam** | **Itelman** | **Mbuti** | **0.0154** | **2.17** | **0.0071** | **4,650** | **4,508** | **360,258** | **Siberia** |
| Salkhit_bl_all | Salkhit_bl_all_deam | Aleut | Mbuti | 0.0073 | 1.17 | 0.0062 | 4,579 | 4,512 | 360,373 | Siberia |
| Salkhit_bl_all | Salkhit_bl_all_deam | Chukchi | Mbuti | 0.0081 | 1.18 | 0.0068 | 4,562 | 4,489 | 360,249 | Siberia |
| Salkhit_bl_all | Salkhit_bl_all_deam | Altaian | Mbuti | 0.0041 | 0.56 | 0.0072 | 4,559 | 4,522 | 360,223 | Central Eurasia |
| Salkhit_bl_all | Salkhit_bl_all_deam | Kyrgyz | Mbuti | 0.0095 | 1.55 | 0.0061 | 4,603 | 4,516 | 360,378 | Central Eurasia |
| Salkhit_bl_all | Salkhit_bl_all_deam | Iranian | Mbuti | 0.0073 | 1.24 | 0.0058 | 4,508 | 4,443 | 360,378 | Central Eurasia |
| Salkhit_bl_all | Salkhit_bl_all_deam | Mala | Mbuti | 0.0051 | 0.85 | 0.0059 | 4,517 | 4,471 | 360,390 | Central Eurasia |
| **Salkhit_bl_all** | **Salkhit_bl_all_deam** | **Russian** | **Mbuti** | **0.0137** | **2.26** | **0.0060** | **4,576** | **4,452** | **360,367** | **West Eurasia** |
| Salkhit_bl_all | Salkhit_bl_all_deam | Czech | Mbuti | 0.0046 | 0.62 | 0.0073 | 4,528 | 4,487 | 360,245 | West Eurasia |
| Salkhit_bl_all | Salkhit_bl_all_deam | Saami | Mbuti | 0.0123 | 1.94 | 0.0063 | 4,589 | 4,477 | 360,386 | West Eurasia |
| Salkhit_bl_all | Salkhit_bl_all_deam | French | Mbuti | 0.0084 | 1.41 | 0.0059 | 4,537 | 4,461 | 360,390 | West Eurasia |
| Salkhit_bl_all | Salkhit_bl_all_deam | Spanish | Mbuti | 0.0081 | 1.35 | 0.0060 | 4,529 | 4,456 | 360,379 | West Eurasia |
| Salkhit_bl_all | Salkhit_bl_all_deam | Sardinian | Mbuti | 0.0101 | 1.77 | 0.0057 | 4,538 | 4,447 | 360,386 | West Eurasia |
| Salkhit_bl_all | Salkhit_bl_all_deam | Mayan | Mbuti | 0.0024 | 0.36 | 0.0067 | 4,600 | 4,578 | 360,358 | America |
| Salkhit_bl_all | Salkhit_bl_all_deam | Quechua | Mbuti | 0.0042 | 0.64 | 0.0065 | 4,605 | 4,567 | 360,385 | America |
| Salkhit_bl_all | Salkhit_bl_all_deam | Cree | Mbuti | 0.0098 | 1.53 | 0.0064 | 4,579 | 4,490 | 360,369 | America |
| Salkhit_bl_all | Salkhit_bl_all_deam | Karitiana | Mbuti | 0.0087 | 1.30 | 0.0066 | 4,627 | 4,548 | 360,376 | America |
| Salkhit_bl_all | Salkhit_bl_all_deam | Mixe | Mbuti | 0.0014 | 0.20 | 0.0068 | 4,604 | 4,592 | 360,380 | America |

**Table S6B.** *D-statistic* comparing the genetic similarity of present-day human populations to the set of all Salkhit sequences from bleached extracts only "Salkhit_bl_all" and the according set of deaminated sequences "Salkhit_bl_all_deam". Significant D-statistics (|Zscore| > 2) are underline in red.

| Pop1 | Pop2 | Pop3 | Outgroup | Dscore | Zscore | std.err | nbr of BABA sites | nbr of ABBA sites | nbr of SNPs present in the all dataset | Pop3 geographic origin |
|---|---|---|---|---|---|---|---|---|---|---|
| Salkhit_bl_A | salkhit_bl_A_deam | Ami | Mbuti | 0.0052 | 0.59 | 0.0088 | 2,341 | 2,317 | 246,845 | East Eurasia |
| Salkhit_bl_A | salkhit_bl_A_deam | Han | Mbuti | 0.0062 | 0.74 | 0.0084 | 2,346 | 2,318 | 246,866 | East Eurasia |
| Salkhit_bl_A | salkhit_bl_A_deam | Dai | Mbuti | 0.0082 | 0.97 | 0.0084 | 2,344 | 2,305 | 246,870 | East Eurasia |
| Salkhit_bl_A | salkhit_bl_A_deam | Korean | Mbuti | -0.006 | -0.66 | 0.0090 | 2,317 | 2,345 | 246,852 | East Eurasia |
| Salkhit_bl_A | salkhit_bl_A_deam | Japanese | Mbuti | 0.0103 | 1.2 | 0.0086 | 2,365 | 2,317 | 246,864 | East Eurasia |
| Salkhit_bl_A | salkhit_bl_A_deam | Atayal | Mbuti | 0.0018 | 0.16 | 0.0111 | 2,344 | 2,335 | 246,755 | Southeast Asia |
| Salkhit_bl_A | salkhit_bl_A_deam | Thai | Mbuti | 0.0096 | 1.06 | 0.0090 | 2,354 | 2,310 | 246,855 | Southeast Asia |
| Salkhit_bl_A | salkhit_bl_A_deam | Cambodian | Mbuti | 0.0156 | 1.71 | 0.0091 | 2,359 | 2,286 | 246,855 | Southeast Asia |
| Salkhit_bl_A | salkhit_bl_A_deam | Kinh | Mbuti | 0.0113 | 1.25 | 0.0090 | 2,349 | 2,296 | 246,849 | Southeast Asia |
| Salkhit_bl_A | salkhit_bl_A_deam | Igorot | Mbuti | 0.0083 | 0.86 | 0.0096 | 2,348 | 2,310 | 246,844 | Southeast Asia |
| Salkhit_bl_A | salkhit_bl_A_deam | Hawaiian | Mbuti | -0.0039 | -0.39 | 0.0100 | 2,313 | 2,331 | 246,769 | Oceania |
| Salkhit_bl_A | salkhit_bl_A_deam | Australian | Mbuti | -0.0008 | -0.09 | 0.0093 | 2,290 | 2,294 | 246,866 | Oceania |
| Salkhit_bl_A | salkhit_bl_A_deam | Papuan | Mbuti | 0.0024 | 0.27 | 0.0088 | 2,297 | 2,286 | 246,870 | Oceania |
| Salkhit_bl_A | salkhit_bl_A_deam | Bougainville | Mbuti | 0.0027 | 0.29 | 0.0093 | 2,300 | 2,287 | 246,841 | Oceania |
| Salkhit_bl_A | salkhit_bl_A_deam | Maori | Mbuti | 0.0111 | 1.09 | 0.01020 | 2,314 | 2,263 | 246,756 | Oceania |
| Salkhit_bl_A | salkhit_bl_A_deam | Ulchi | Mbuti | 0.0108 | 1.18 | 0.0091 | 2,356 | 2,305 | 246,847 | Siberia |
| Salkhit_bl_A | salkhit_bl_A_deam | Yakut | Mbuti | 0.0036 | 0.39 | 0.0093 | 2,344 | 2,327 | 246,851 | Siberia |
| Salkhit_bl_A | salkhit_bl_A_deam | Itelman | Mbuti | 0.0096 | 0.98 | 0.0098 | 2,354 | 2,309 | 246,792 | Siberia |
| Salkhit_bl_A | salkhit_bl_A_deam | Aleut | Mbuti | 0.0103 | 1.12 | 0.0092 | 2,346 | 2,298 | 246,856 | Siberia |
| **Salkhit_bl_A** | **salkhit_bl_A_deam** | **Chukchi** | **Mbuti** | **0.0235** | **2.39** | **0.0098** | **2,373** | **2,264** | **246,776** | **Siberia** |
| Salkhit_bl_A | salkhit_bl_A_deam | Altaian | Mbuti | 0.0099 | 0.96 | 0.0103 | 2,363 | 2,316 | 246,744 | Central Eurasia |
| Salkhit_bl_A | salkhit_bl_A_deam | Kyrgyz | Mbuti | 0.0103 | 1.2 | 0.0086 | 2,349 | 2,301 | 246,861 | Central Eurasia |
| Salkhit_bl_A | salkhit_bl_A_deam | Iranian | Mbuti | 0.0122 | 1.33 | 0.0091 | 2,302 | 2,246 | 246,859 | Central Eurasia |
| Salkhit_bl_A | salkhit_bl_A_deam | Mala | Mbuti | 0.0116 | 1.35 | 0.0086 | 2,316 | 2,263 | 246,864 | Central Eurasia |
| Salkhit_bl_A | salkhit_bl_A_deam | Russian | Mbuti | 0.0098 | 1.08 | 0.0090 | 2,311 | 2,266 | 246,851 | West Eurasia |
| Salkhit_bl_A | salkhit_bl_A_deam | Czech | Mbuti | 0.0139 | 1.41 | 0.0098 | 2,298 | 2,235 | 246,771 | West Eurasia |
| Salkhit_bl_A | salkhit_bl_A_deam | Saami | Mbuti | 0.0076 | 0.85 | 0.0089 | 2,320 | 2,285 | 246,865 | West Eurasia |
| Salkhit_bl_A | salkhit_bl_A_deam | French | Mbuti | 0.014 | 1.64 | 0.0085 | 2,310 | 2,246 | 246,867 | West Eurasia |
| Salkhit_bl_A | salkhit_bl_A_deam | Spanish | Mbuti | 0.0119 | 1.29 | 0.0092 | 2,307 | 2,253 | 246,859 | West Eurasia |
| Salkhit_bl_A | salkhit_bl_A_deam | Sardinian | Mbuti | 0.0153 | 1.74 | 0.0088 | 2,322 | 2,252 | 246,866 | West Eurasia |
| Salkhit_bl_A | salkhit_bl_A_deam | Mayan | Mbuti | 0.0045 | 0.46 | 0.0096 | 2,352 | 2,331 | 246,845 | America |
| Salkhit_bl_A | salkhit_bl_A_deam | Quechua | Mbuti | 0.0005 | 0.05 | 0.0106 | 2,341 | 2,338 | 246,865 | America |
| Salkhit_bl_A | salkhit_bl_A_deam | Cree | Mbuti | 0.0054 | 0.57 | 0.0094 | 2,319 | 2,294 | 246,857 | America |
| Salkhit_bl_A | salkhit_bl_A_deam | Karitiana | Mbuti | 0.0131 | 1.33 | 0.0098 | 2,378 | 2,317 | 246,857 | America |
| Salkhit_bl_A | salkhit_bl_A_deam | Mixe | Mbuti | 0.004 | 0.41 | 0.0096 | 2,359 | 2,340 | 246,862 | America |

**Table S6C.** *D-statistic* comparing the genetic similarity of present-day human populations to the set of Salkhit sequences from bleached library A "Salkhit_bl_A" and the according set of deaminated sequences "Salkhit_bl_A_deam". Significant D-statistics (|Zscore| > 2) are underline in red.

| Pop1 | Pop2 | Pop3 | Outgroup | Dscore | Zscore | std.err | nbr of BABA sites | nbr of ABBA sites | nbr of SNPs present in the all dataset | Pop3 geographic origin |
|---|---|---|---|---|---|---|---|---|---|---|
| Salkhit_all_deam | Salkhit_bl_all_deam | Ami | Mbuti | 0.0041 | 0.34 | 0.0122 | 1,238 | 1,228 | 361,003 | East Eurasia |
| Salkhit_all_deam | Salkhit_bl_all_deam | Han | Mbuti | 0.0085 | 0.76 | 0.0112 | 1,229 | 1,208 | 361,034 | East Eurasia |
| Salkhit_all_deam | Salkhit_bl_all_deam | Dai | Mbuti | 0.0073 | 0.63 | 0.0116 | 1,227 | 1,209 | 361,037 | East Eurasia |
| Salkhit_all_deam | Salkhit_bl_all_deam | Korean | Mbuti | 0.0023 | 0.19 | 0.0120 | 1,232 | 1,226 | 361,008 | East Eurasia |
| Salkhit_all_deam | Salkhit_bl_all_deam | Japanese | Mbuti | 0.0053 | 0.45 | 0.0117 | 1,227 | 1,214 | 361,031 | East Eurasia |
| Salkhit_all_deam | Salkhit_bl_all_deam | Atayal | Mbuti | 0.0048 | 0.33 | 0.0145 | 1,218 | 1,206 | 360,869 | Southeast Asia |
| Salkhit_all_deam | Salkhit_bl_all_deam | Thai | Mbuti | 0.0009 | 0.07 | 0.0123 | 1,225 | 1,223 | 361,014 | Southeast Asia |
| Salkhit_all_deam | Salkhit_bl_all_deam | Cambodian | Mbuti | 0.0162 | 1.26 | 0.0128 | 1,248 | 1,208 | 361,019 | Southeast Asia |
| Salkhit_all_deam | Salkhit_bl_all_deam | Kinh | Mbuti | 0.0064 | 0.50 | 0.0127 | 1,229 | 1,213 | 361,006 | Southeast Asia |
| Salkhit_all_deam | Salkhit_bl_all_deam | Igorot | Mbuti | 0.002 | 0.16 | 0.0128 | 1,224 | 1,220 | 361,002 | Southeast Asia |
| Salkhit_all_deam | Salkhit_bl_all_deam | Hawaiian | Mbuti | 0.0217 | 1.58 | 0.0136 | 1,257 | 1,204 | 360,867 | Oceania |
| Salkhit_all_deam | Salkhit_bl_all_deam | Australian | Mbuti | 0.0117 | 0.9 | 0.0130 | 1,222 | 1,194 | 361,028 | Oceania |
| Salkhit_all_deam | Salkhit_bl_all_deam | Papuan | Mbuti | 0.0161 | 1.33 | 0.0121 | 1,216 | 1,178 | 361,039 | Oceania |
| Salkhit_all_deam | Salkhit_bl_all_deam | Bougainville | Mbuti | 0.0161 | 1.16 | 0.0138 | 1,236 | 1,197 | 361,000 | Oceania |
| Salkhit_all_deam | Salkhit_bl_all_deam | Maori | Mbuti | 0.003 | 0.22 | 0.0134 | 1,221 | 1,214 | 360,847 | Oceania |
| Salkhit_all_deam | Salkhit_bl_all_deam | Ulchi | Mbuti | -0.0018 | -0.14 | 0.0126 | 1,222 | 1,226 | 361,010 | Siberia |
| Salkhit_all_deam | Salkhit_bl_all_deam | Yakut | Mbuti | 0.016 | 1.31 | 0.0122 | 1,241 | 1,202 | 361,011 | Siberia |
| Salkhit_all_deam | Salkhit_bl_all_deam | Itelman | Mbuti | 0.019 | 1.3 | 0.0146 | 1,254 | 1,208 | 360,903 | Siberia |
| Salkhit_all_deam | Salkhit_bl_all_deam | Aleut | Mbuti | 0.0134 | 1.09 | 0.0122 | 1,234 | 1,201 | 361,017 | Siberia |
| Salkhit_all_deam | Salkhit_bl_all_deam | Chukchi | Mbuti | 0.0117 | 0.83 | 0.0141 | 1,231 | 1,203 | 360,894 | Siberia |
| Salkhit_all_deam | Salkhit_bl_all_deam | Altaian | Mbuti | -0.0008 | -0.05 | 0.0145 | 1,203 | 1,205 | 360,866 | Central Eurasia |
| Salkhit_all_deam | Salkhit_bl_all_deam | Kyrgyz | Mbuti | 0.0068 | 0.56 | 0.0120 | 1,221 | 1,205 | 361,023 | Central Eurasia |
| Salkhit_all_deam | Salkhit_bl_all_deam | Iranian | Mbuti | 0.019 | 1.55 | 0.0122 | 1,218 | 1,173 | 361,023 | Central Eurasia |
| Salkhit_all_deam | Salkhit_bl_all_deam | Mala | Mbuti | 0.0188 | 1.62 | 0.0115 | 1,225 | 1,179 | 361,034 | Central Eurasia |
| Salkhit_all_deam | Salkhit_bl_all_deam | Russian | Mbuti | 0.0249 | 1.87 | 0.0133 | 1,225 | 1,166 | 361,012 | West Eurasia |
| Salkhit_all_deam | Salkhit_bl_all_deam | Czech | Mbuti | -0.0006 | -0.04 | 0.0139 | 1,199 | 1,201 | 360,889 | West Eurasia |
| Salkhit_all_deam | Salkhit_bl_all_deam | Saami | Mbuti | 0.025 | 1.82 | 0.0137 | 1,240 | 1,180 | 361,031 | West Eurasia |
| Salkhit_all_deam | Salkhit_bl_all_deam | French | Mbuti | 0.0152 | 1.18 | 0.0129 | 1,220 | 1,184 | 361,035 | West Eurasia |
| Salkhit_all_deam | Salkhit_bl_all_deam | Spanish | Mbuti | 0.0176 | 1.32 | 0.0133 | 1,215 | 1,173 | 361,023 | West Eurasia |
| Salkhit_all_deam | Salkhit_bl_all_deam | Sardinian | Mbuti | 0.005 | 0.40 | 0.0123 | 1,210 | 1,198 | 361,031 | West Eurasia |
| Salkhit_all_deam | Salkhit_bl_all_deam | Mayan | Mbuti | 0.0081 | 0.61 | 0.0133 | 1,243 | 1,223 | 361,003 | America |
| Salkhit_all_deam | Salkhit_bl_all_deam | Quechua | Mbuti | 0.0095 | 0.72 | 0.0131 | 1,238 | 1,215 | 361,030 | America |
| Salkhit_all_deam | Salkhit_bl_all_deam | Cree | Mbuti | 0.0117 | 0.88 | 0.0132 | 1,221 | 1,193 | 361,014 | America |
| Salkhit_all_deam | Salkhit_bl_all_deam | Karitiana | Mbuti | 0.0045 | 0.32 | 0.0141 | 1,242 | 1,231 | 361,021 | America |
| Salkhit_all_deam | Salkhit_bl_all_deam | Mixe | Mbuti | 0.0121 | 0.93 | 0.0129 | 1,244 | 1,214 | 361,024 | America |

**Table S6D.** *D-statistic* comparing the genetic similarity of present-day human populations to the set of Salkhit all deaminated sequences "Salkhit_all_deam" and the set of deaminated sequences from bleached library only "Salkhit_bl_all_deam".

| Population1 | Population2 | Population test | Population3 | Outgroup | *f4-ratio* | std.err | Zscore |
|---|---|---|---|---|---|---|---|
| French | Spanish | Salkhit_all | Tianyuan_MH | Mbuti | 0.32 | 0.03 | 10.5 |
| French | Spanish | Salkhit_all_deam | Tianyuan_MH | Mbuti | 0.17 | 0.04 | 4.1 |
| French | Spanish | Salkhit_bl_all | Tianyuan_MH | Mbuti | 0.19 | 0.04 | 5.4 |
| French | Spanish | Salkhit_bl_all_deam | Tianyuan_MH | Mbuti | 0.16 | 0.04 | 3.6 |
| French | Spanish | Salkhit_bl_A | Tianyuan_MH | Mbuti | 0.19 | 0.04 | 5.0 |
| French | Spanish | Salkhit_bl_A_deam | Tianyuan_MH | Mbuti | 0.12 | 0.045 | 2.6 |

**Table S7 Proportion of present-day European Alleles in the Salkhit genome estimated by *f4-ratio* statistics.**
Bl=Bleached, deam = sequences showing evidence for deamination at the first or last base.

| Chrom. | Chrom. size (base) | # base covered/chrom. in salkhitx6_all | % chrom. covered in salkhitx6_all | # base covered/chrom. in salkhitx6_deam | % chrom. covered in salkhitx6_deam |
|---|---|---|---|---|---|
| 1 | 225,280,621 | 932,729 | 0.00414 | 116,858 | 0.00052 |
| 2 | 238,204,518 | 1,021,922 | 0.00429 | 126,952 | 0.00053 |
| 3 | 194,797,135 | 852,112 | 0.00437 | 99,959 | 0.00051 |
| 4 | 187,661,676 | 808,593 | 0.00431 | 95,549 | 0.00051 |
| 5 | 177,695,260 | 763,216 | 0.00429 | 94,436 | 0.00053 |
| 6 | 167,395,066 | 717,377 | 0.00428 | 84,423 | 0.00050 |
| 7 | 155,353,663 | 656,718 | 0.00423 | 83,628 | 0.00054 |
| 8 | 142,888,922 | 622,711 | 0.00436 | 78,066 | 0.00055 |
| 9 | 120,143,431 | 471,706 | 0.00393 | 59,926 | 0.0005 |
| 10 | 131,314,738 | 553,393 | 0.00421 | 68,916 | 0.00052 |
| 11 | 131,129,516 | 556,312 | 0.00424 | 72,062 | 0.00055 |
| 12 | 130,481,393 | 554,154 | 0.00425 | 72,117 | 0.00055 |
| 13 | 95,589,878 | 413,259 | 0.00432 | 47,741 | 0.0005 |
| 14 | 88,289,540 | 379,877 | 0.00430 | 47,379 | 0.00054 |
| 15 | 81,694,766 | 339,286 | 0.00415 | 41,885 | 0.00051 |
| 16 | 78,884,753 | 312,584 | 0.00396 | 42,500 | 0.00054 |
| 17 | 77,795,210 | 312,452 | 0.00402 | 42,024 | 0.00054 |
| 18 | 74,657,229 | 325,671 | 0.00436 | 39,328 | 0.00053 |
| 19 | 55,808,983 | 216,003 | 0.00387 | 28,568 | 0.00051 |
| 20 | 59,505,520 | 261,103 | 0.00439 | 36,636 | 0.00061 |
| 21 | 35,106,642 | 142,745 | 0.00407 | 18,490 | 0.00053 |
| 22 | 34,894,545 | 132,316 | 0.00379 | 18,559 | 0.00053 |
| X | 151,100,560 | 593,777 | 0.00393 | 74,075 | 0.00049 |
| Y | 22,984,529 | 3,104 | 0.00013 | 127 | 5.52E-06 |
| auto | 2,684,573,005 | 11,346,239 | 0.00423 | 1,416,002 | 0.00052 |

**Table S8 Sex determination of the Salkhit individual.**
Chrom=Chromosome; Salkhitx6_all: All sequences from shotgun of the 6 Salkhit libraries; Salkhitx6_deam: Deaminated sequences from shotgun of the 6 Salkhit libraries

| | % All | % Modern Human | % Neandertal | % Denisova |
|---|---|---|---|---|
| **Salkhit all_fragment** | 97.56 (97.39-97.72) | 33.06 (31.66-34.45) | 4.25 (2.68-5.82) | 7.99 (7.01-8.97) |
| **Salkhit deam_fragment** | 95.5 (94.89-96.11) | 32.35 (28.54-36.17) | 5.41 (0.25-10.56) | 6.94 (4.32-9.57) |
| **Dai** | 98.66 (98.66-98.67) | 32.23 (32.18-32.28) | 4.09 (4.04-4.15) | 6.74 (6.7-6.77) |
| **French** | 98.71 (98.7-98.71) | 32.55 (32.51-32.6) | 4.07 (4.02-4.12) | 6.73 (6.7-6.76) |
| **Han** | 98.67 (98.66-98.67) | 32.2 (32.14-32.25) | 4.45 (4.4-4.51) | 6.68 (6.64-6.71) |
| **Papuan** | 98.73 (98.73-98.74) | 31.02 (30.98-31.07) | 4.15 (4.1-4.2) | 8.31 (8.28-8.35) |
| **Sardinian** | 98.67 (98.66-98.67) | 32.2 (32.15-32.25) | 4.28 (4.22-4.33) | 6.58 (6.55-6.61) |
| **Karitiana** | 98.74 (98.73-98.74) | 32.31 (32.26-32.36) | 4.26 (4.2-4.32) | 6.5 (6.47-6.54) |

**Table S9 Hominin group assignment of the Salkhit individual.**
Proportion of shared derived alleles and 95% binomial confidence interval in brackets.

| Source 2 | Outgroup | Country | nbr_ind. | ancient | Salkhit_all_deam_f3 | std.err | Z | #SNPs |
|---|---|---|---|---|---|---|---|---|
| **Yi** | Mbuti | China | 2 | modern | **0.2431** | 0.0033 | 72.819 | 390,617 |
| **Mayan** | Mbuti | Mexico | 2 | modern | **0.2428** | 0.0033 | 73.677 | 386,998 |
| **She** | Mbuti | China | 2 | modern | **0.2427** | 0.00323 | 75.098 | 389,761 |
| **Eskimo_Naukan** | Mbuti | Russia | 2 | modern | **0.2424** | 0.00333 | 72.717 | 387,885 |
| **Eskimo_Chaplin** | Mbuti | Russia | 1 | modern | **0.2423** | 0.00351 | 69.003 | 375,558 |
| **Mongola** | Mbuti | China | 1 | modern | **0.2423** | 0.00326 | 74.304 | 391,285 |
| **Han** | Mbuti | China | 4 | modern | **0.2419** | 0.00316 | 76.445 | 407,549 |
| **Atayal** | Mbuti | Taiwan | 1 | modern | **0.2419** | 0.00350 | 69.021 | 375,063 |
| **Ami** | Mbuti | Taiwan | 2 | modern | **0.2418** | 0.00329 | 73.426 | 388,349 |
| **Oroqen** | Mbuti | China | 2 | modern | **0.2418** | 0.00333 | 72.556 | 389,896 |
| **Piapoco** | Mbuti | Colombia | 2 | modern | **0.2417** | 0.00329 | 73.531 | 385,562 |
| **Korean** | Mbuti | Korea | 2 | modern | **0.2417** | 0.00324 | 74.466 | 390,592 |
| **Tujia** | Mbuti | China | 2 | modern | **0.2416** | 0.00344 | 70.309 | 390,286 |
| **Ulchi** | Mbuti | Russia | 2 | modern | **0.2416** | 0.00332 | 72.665 | 389,474 |
| **Eskimo_Sireniki** | Mbuti | Russia | 2 | modern | **0.2415** | 0.00331 | 72.911 | 389,109 |
| **Chipewyan** | Mbuti | Canada | 2 | modern | **0.2414** | 0.00341 | 70.883 | 386,188 |
| **Japanese** | Mbuti | Japan | 3 | modern | **0.2411** | 0.00322 | 74.781 | 398,088 |
| **Zapotec** | Mbuti | Mexico | 2 | modern | **0.2411** | 0.00328 | 73.509 | 387,668 |
| **Itelman** | Mbuti | Russia | 1 | modern | **0.2408** | 0.00362 | 66.527 | 375,702 |
| **Naxi** | Mbuti | China | 3 | modern | **0.2406** | 0.00319 | 75.389 | 399,025 |
| **Quechua** | Mbuti | Peru | 3 | modern | **0.2405** | 0.00337 | 71.463 | 393,655 |
| **Hezhen** | Mbuti | China | 2 | modern | **0.2405** | 0.00328 | 73.35 | 390,101 |
| **Xibo** | Mbuti | China | 2 | modern | **0.2403** | 0.00336 | 71.455 | 391,014 |
| **Karitiana** | Mbuti | Brazil | 4 | modern | **0.2403** | 0.0034 | 70.767 | 391,672 |
| **Kinh** | Mbuti | Vietnam | 2 | modern | **0.2403** | 0.00327 | 73.415 | 390,441 |
| **Dusun** | Mbuti | Brunei | 2 | modern | **0.24** | 0.00326 | 73.702 | 388,543 |
| **Dai** | Mbuti | China | 5 | modern | **0.24** | 0.00319 | 75.307 | 408,846 |
| **Yakut** | Mbuti | Russia | 2 | modern | **0.2398** | 0.00332 | 72.232 | 390,872 |
| **Even** | Mbuti | Russia | 3 | modern | **0.2398** | 0.00327 | 73.32 | 399,035 |
| **Chane** | Mbuti | Argentina | 1 | modern | **0.2396** | 0.00338 | 70.932 | 375,395 |
| **Igorot** | Mbuti | Philippines | 2 | modern | **0.2394** | 0.00337 | 71.034 | 387,560 |
| **Lahu** | Mbuti | China | 2 | modern | **0.2394** | 0.00327 | 73.129 | 389,431 |
| **Miao** | Mbuti | China | 2 | modern | **0.2393** | 0.00326 | 73.432 | 389,923 |
| **Thai** | Mbuti | Thailand | 2 | modern | **0.2391** | 0.00332 | 72.1 | 391,341 |
| **Daur** | Mbuti | China | 1 | modern | **0.2391** | 0.00342 | 69.996 | 376,617 |
| **Mixe** | Mbuti | Mexico | 3 | modern | **0.239** | 0.00323 | 73.99 | 392,008 |
| **Nahua** | Mbuti | Mexico | 1 | modern | **0.2386** | 0.00341 | 69.897 | 375,760 |
| **Pima** | Mbuti | Mexico | 2 | modern | **0.2378** | 0.00328 | 72.595 | 385,559 |
| **Hawaiian** | Mbuti | USA | 1 | modern | **0.2376** | 0.00344 | 69.128 | 376,295 |
| **Tu** | Mbuti | China | 2 | modern | **0.2376** | 0.00321 | 73.976 | 391,532 |
| **Cambodian** | Mbuti | Cambodia | 2 | modern | **0.2374** | 0.00329 | 72.179 | 391,322 |
| **Burmese** | Mbuti | Myanmar | 2 | modern | **0.2367** | 0.00321 | 73.758 | 392,036 |
| **Kyrgyz** | Mbuti | Kyrgyzstan | 2 | modern | **0.2354** | 0.00321 | 73.434 | 394,036 |
| **Australian** | Mbuti | Australia | 4 | modern | **0.2352** | 0.00457 | 51.474 | 99,054 |
| **Altaian** | Mbuti | Russia | 1 | modern | **0.2352** | 0.00344 | 68.379 | 377,321 |
| **Mixtec** | Mbuti | Mexico | 2 | modern | **0.2338** | 0.00325 | 71.91 | 390,493 |
| **Cree** | Mbuti | Canada | 2 | modern | **0.2338** | 0.00331 | 70.625 | 392,576 |
| **Tubalar** | Mbuti | Russia | 2 | modern | **0.2333** | 0.00335 | 69.711 | 393,481 |
| **Aleut** | Mbuti | Russia | 2 | modern | **0.2332** | 0.00329 | 70.943 | 392,879 |
| **Kusunda** | Mbuti | Nepal | 2 | modern | **0.2329** | 0.00323 | 72.093 | 388,608 |
| **Mansi** | Mbuti | Russia | 2 | modern | **0.2328** | 0.00325 | 71.557 | 394,054 |
| **Khonda_Dora** | Mbuti | India | 1 | modern | **0.2325** | 0.00338 | 68.706 | 376,718 |
| **Tlingit** | Mbuti | Russia | 2 | modern | **0.2321** | 0.00324 | 71.535 | 394,394 |
| **Bougainville** | Mbuti | PapuaNewGuinea | 2 | modern | **0.2312** | 0.00322 | 71.813 | 386,611 |
| **Papuan** | Mbuti | PapuaNewGuinea | 16 | modern | **0.2311** | 0.0033 | 70.035 | 410,598 |
| **Chukchi** | Mbuti | Russia | 1 | modern | **0.2310** | 0.00334 | 69.068 | 379,167 |
| **Hazara** | Mbuti | Pakistan | 2 | modern | **0.2296** | 0.00329 | 69.71 | 394,462 |
| **Uygur** | Mbuti | China | 2 | modern | **0.2288** | 0.00320 | 71.443 | 395,371 |
| **Saami** | Mbuti | Finland | 2 | modern | **0.228** | 0.00328 | 69.505 | 393,898 |
| **Mala** | Mbuti | India | 3 | modern | **0.2278** | 0.00311 | 73.34 | 403,460 |
| **Madiga** | Mbuti | India | 2 | modern | **0.2276** | 0.00323 | 70.462 | 393,761 |
| **Relli** | Mbuti | India | 2 | modern | **0.2256** | 0.00311 | 72.542 | 393,720 |
| **Kapu** | Mbuti | India | 2 | modern | **0.2254** | 0.0032 | 70.433 | 394,122 |
| **Yadava** | Mbuti | India | 2 | modern | **0.2253** | 0.00312 | 72.148 | 393,455 |
| **Bengali** | Mbuti | Bangladesh | 2 | modern | **0.2252** | 0.00315 | 71.394 | 394,491 |
| **Irula** | Mbuti | India | 2 | modern | **0.2251** | 0.00309 | 72.831 | 392,642 |
| **Burusho** | Mbuti | Pakistan | 2 | modern | **0.2239** | 0.00314 | 71.33 | 395,044 |
| **Maori** | Mbuti | NewZealand | 1 | modern | **0.2238** | 0.00342 | 65.407 | 379,561 |
| **Punjabi** | Mbuti | Pakistan | 4 | modern | **0.2235** | 0.00301 | 74.138 | 411,248 |
| **Kashmiri_Pandit** | Mbuti | India | 1 | modern | **0.2232** | 0.0034 | 65.693 | 378,172 |
| **Estonian** | Mbuti | Estonia | 2 | modern | **0.2229** | 0.00318 | 70.16 | 395,637 |
| **Brahmin** | Mbuti | India | 2 | modern | **0.2225** | 0.00315 | 70.534 | 395,025 |
| **Russian** | Mbuti | Russia | 2 | modern | **0.2223** | 0.00324 | 68.556 | 395,488 |
| **Pathan** | Mbuti | Pakistan | 2 | modern | **0.2219** | 0.0032 | 69.369 | 395,019 |
| **Icelandic** | Mbuti | Iceland | 2 | modern | **0.2219** | 0.00322 | 68.846 | 395,332 |

| | | | | | | | | |
|---|---|---|---|---|---|---|---|---|
| **Sindhi** | Mbuti | Pakistan | 2 | modern | **0.2216** | 0.00314 | 70.438 | 394,647 |
| **Finnish** | Mbuti | Finland | 3 | modern | **0.2216** | 0.00305 | 72.579 | 405,374 |
| **Czech** | Mbuti | Czechoslovia | 1 | modern | **0.2205** | 0.00338 | 65.186 | 379,164 |
| **Kalash** | Mbuti | Pakistan | 2 | modern | **0.2205** | 0.00314 | 70.125 | 392,573 |
| **Polish** | Mbuti | Poland | 1 | modern | **0.2205** | 0.00337 | 65.335 | 379,095 |
| **French** | Mbuti | France | 3 | modern | **0.2203** | 0.00310 | 70.972 | 405,402 |
| **Tajik** | Mbuti | Tajikistan | 2 | modern | **0.2203** | 0.00322 | 68.367 | 394,738 |
| **English** | Mbuti | England | 2 | modern | **0.2185** | 0.00316 | 69.172 | 395,517 |
| **Basque** | Mbuti | France | 2 | modern | **0.2183** | 0.00319 | 68.426 | 395,045 |
| **Abkhasian** | Mbuti | Abkhazia | 2 | modern | **0.2180** | 0.00320 | 68.112 | 395,075 |
| **Albanian** | Mbuti | Albania | 1 | modern | **0.2178** | 0.00324 | 67.325 | 378,851 |
| **Orcadian** | Mbuti | OrkneyIslands | 2 | modern | **0.2178** | 0.00309 | 70.536 | 395,689 |
| **Bulgarian** | Mbuti | Bulgaria | 2 | modern | **0.2174** | 0.00316 | 68.8 | 395,610 |
| **Adygei** | Mbuti | Russia(Caucasus) | 2 | modern | **0.2173** | 0.00320 | 67.904 | 395,564 |
| **Turkish** | Mbuti | Turkey | 2 | modern | **0.2173** | 0.00317 | 68.473 | 395,579 |
| **North_Ossetian** | Mbuti | Russia | 2 | modern | **0.2172** | 0.00315 | 68.866 | 395,743 |
| **Hungarian** | Mbuti | Hungary | 2 | modern | **0.2171** | 0.00319 | 68.105 | 395,723 |
| **Greek** | Mbuti | Greece | 2 | modern | **0.2169** | 0.00310 | 69.902 | 395,128 |
| **Chechen** | Mbuti | Russia | 1 | modern | **0.2168** | 0.00334 | 64.893 | 378,757 |
| **Brahui** | Mbuti | Pakistan | 2 | modern | **0.2167** | 0.00311 | 69.78 | 394,796 |
| **Bergamo** | Mbuti | Italy(Bergamo) | 1 | modern | **0.2163** | 0.00325 | 66.575 | 379,524 |
| **Tuscan** | Mbuti | Italy | 2 | modern | **0.2163** | 0.00312 | 69.248 | 395,617 |
| **Spanish** | Mbuti | Spain | 2 | modern | **0.2162** | 0.00316 | 68.475 | 395,465 |
| **Lezgin** | Mbuti | Russia | 2 | modern | **0.2158** | 0.00319 | 67.641 | 395,233 |
| **Georgian** | Mbuti | Georgia | 2 | modern | **0.215** | 0.00317 | 67.948 | 395,308 |
| **Armenian** | Mbuti | Armenia | 2 | modern | **0.2148** | 0.00307 | 69.844 | 387,088 |
| **Iranian** | Mbuti | Iran | 2 | modern | **0.2145** | 0.00306 | 70.045 | 395,405 |
| **Sardinian** | Mbuti | Italy | 3 | modern | **0.2142** | 0.00309 | 69.392 | 404,732 |
| **Balochi** | Mbuti | Pakistan | 1 | modern | **0.2133** | 0.00337 | 63.229 | 377,253 |
| **Makrani** | Mbuti | Pakistan | 2 | modern | **0.2128** | 0.00307 | 69.314 | 394,607 |
| **Iraqi_Jew** | Mbuti | Iraq | 2 | modern | **0.2108** | 0.00309 | 68.233 | 394,927 |
| **Druze** | Mbuti | Israel(Carmel) | 2 | modern | **0.2095** | 0.00299 | 69.984 | 395,300 |
| **Yemenite_Jew** | Mbuti | Yemen | 2 | modern | **0.2055** | 0.00311 | 66.141 | 394,699 |
| **Jordanian** | Mbuti | Jordan | 3 | modern | **0.2043** | 0.00299 | 68.425 | 405,791 |
| **Palestinian** | Mbuti | Israel | 3 | modern | **0.2034** | 0.00301 | 67.648 | 406,412 |
| **BedouinB** | Mbuti | Israel(Negev) | 2 | modern | **0.2031** | 0.00301 | 67.299 | 392,903 |
| **Saharawi** | Mbuti | Morocco | 2 | modern | **0.1835** | 0.00293 | 62.566 | 396,550 |
| **Mozabite** | Mbuti | Algeria | 2 | modern | **0.1806** | 0.00295 | 61.298 | 396,737 |
| **Somali** | Mbuti | Kenya | 1 | modern | **0.1391** | 0.00298 | 46.716 | 375,646 |
| **Masai** | Mbuti | Kenya | 2 | modern | **0.1130** | 0.00243 | 46.592 | 394,042 |
| **Dinka** | Mbuti | Sudan | 3 | modern | **0.0881** | 0.00216 | 40.809 | 404,506 |
| **Mandenka** | Mbuti | Senegal | 4 | modern | **0.0839** | 0.00207 | 40.563 | 421,753 |
| **Gambian** | Mbuti | Gambia | 2 | modern | **0.0837** | 0.00224 | 37.372 | 396,339 |
| **Luo** | Mbuti | Kenya | 2 | modern | **0.0834** | 0.00225 | 37.09 | 395,139 |
| **Luhya** | Mbuti | Kenya | 2 | modern | **0.0814** | 0.00219 | 37.094 | 394,633 |
| **BantuKenya** | Mbuti | Kenya | 2 | modern | **0.0806** | 0.00216 | 37.287 | 395,617 |
| **Esan** | Mbuti | Nigeria | 2 | modern | **0.0805** | 0.00220 | 36.559 | 396,885 |
| **Yoruba** | Mbuti | Nigeria | 3 | modern | **0.0787** | 0.00202 | 38.822 | 427,461 |
| **Mende** | Mbuti | SierraLeone | 2 | modern | **0.0785** | 0.00209 | 37.493 | 396,640 |
| **Igbo** | Mbuti | Nigeria | 2 | modern | **0.0779** | 0.0022 | 35.415 | 396,838 |
| **Lemande** | Mbuti | Cameroon | 2 | modern | **0.0771** | 0.00212 | 36.4 | 395,758 |
| **Kongo** | Mbuti | Congo | 1 | modern | **0.077** | 0.00227 | 33.86 | 376,192 |
| **BantuHerero** | Mbuti | BotswanaOrNamibia | 2 | modern | **0.0755** | 0.00209 | 36.14 | 394,706 |
| **BantuTswana** | Mbuti | BotswanaOrNamibia | 2 | modern | **0.0680** | 0.00197 | 34.542 | 395,103 |
| **Biaka** | Mbuti | CentralAfricanRepublic | 2 | modern | **0.0501** | 0.00175 | 28.619 | 388,908 |
| **Ju_hoan_North** | Mbuti | Namibia | 4 | modern | **0.0434** | 0.00168 | 25.808 | 407,071 |

**Table S10** Shared genetic drift between the Salkhit individual and present-day populations estimated by the *f3*-statistics.

| Pop1 | Pop2 | Pop3 | Pop4 | D-Stat | Z | std.err | BABA | ABBA | #SNPs | Pop3 - origin |
|---|---|---|---|---|---|---|---|---|---|---|
| Salkhit_all_deam | Tianyuan | GoyetQ116 | Mbuti | 0.0238 | 2.99 | 0.0079 | 14,790 | 14,103 | 283,946 | ancient |
| Salkhit_all_deam | Tianyuan | Yana1 | Mbuti | 0.0501 | 6.54 | 0.0076 | 22,447 | 20,304 | 454,951 | ancient |
| Salkhit_all_deam | Tianyuan | Yana2 | Mbuti | 0.0461 | 5.88 | 0.0078 | 22,189 | 20,234 | 454,623 | ancient |
| Salkhit_all_deam | Tianyuan | Malta1 | Mbuti | 0.0227 | 3.02 | 0.0075 | 16,614 | 15,876 | 342,502 | ancient |
| Salkhit_all_deam | Tianyuan | Sunghir3 | Mbuti | 0.0246 | 3.20 | 0.0076 | 21,602 | 20,563 | 454,900 | ancient |
| Salkhit_all_deam | Tianyuan | Kolyma1 | Mbuti | 0.0036 | 0.51 | 0.0069 | 21,452 | 21,297 | 454,916 | ancient |
| Salkhit_all_deam | Tianyuan | UstIshim | Mbuti | -0.0052 | -0.71 | 0.0073 | 20,828 | 21,046 | 454,500 | ancient |
| Salkhit_all_deam | Tianyuan | Kostenki14 | Mbuti | 0.0264 | 3.5 | 0.0075 | 21,249 | 20,158 | 443,299 | ancient |
| Salkhit_all_deam | Tianyuan | Vestonice16 | Mbuti | 0.0293 | 3.85 | 0.0076 | 15,994 | 15,084 | 322,349 | ancient |
| Salkhit_all_deam | Tianyuan | ElMiron | Mbuti | 0.0249 | 3.22 | 0.0077 | 14,101 | 13,416 | 285,670 | ancient |
| Salkhit_all_deam | Tianyuan | Saami | Mbuti | 0.0119 | 2.09 | 0.0057 | 22,494 | 21,966 | 473,287 | West Eurasia |
| Salkhit_all_deam | Tianyuan | French | Mbuti | 0.0203 | 3.80 | 0.0053 | 22,460 | 21,568 | 473,297 | West Eurasia |
| Salkhit_all_deam | Tianyuan | Russian | Mbuti | 0.0117 | 2.09 | 0.0055 | 22,331 | 21,814 | 473,271 | West Eurasia |
| Salkhit_all_deam | Tianyuan | Estonian | Mbuti | 0.0212 | 3.84 | 0.0055 | 22,545 | 21,607 | 473,279 | West Eurasia |
| Salkhit_all_deam | Tianyuan | Icelandic | Mbuti | 0.0177 | 3.10 | 0.0057 | 22,471 | 21,687 | 473,276 | West Eurasia |
| Salkhit_all_deam | Tianyuan | Finnish | Mbuti | 0.0198 | 3.69 | 0.0053 | 22,513 | 21,636 | 473,292 | West Eurasia |
| Salkhit_all_deam | Tianyuan | Czech | Mbuti | 0.0233 | 3.66 | 0.0063 | 22,539 | 21,513 | 473,148 | West Eurasia |
| Salkhit_all_deam | Tianyuan | Basque | Mbuti | 0.0204 | 3.66 | 0.0055 | 22,384 | 21,489 | 473,273 | West Eurasia |
| Salkhit_all_deam | Tianyuan | Spanish | Mbuti | 0.0207 | 3.64 | 0.0056 | 22,356 | 21,448 | 473,273 | West Eurasia |
| Salkhit_all_deam | Tianyuan | Sardinian | Mbuti | 0.0146 | 2.69 | 0.0054 | 22,223 | 21,585 | 473,291 | West Eurasia |
| Salkhit_all_deam | Tianyuan | English | Mbuti | 0.0189 | 3.38 | 0.0055 | 22,402 | 21,571 | 473,267 | West Eurasia |
| Salkhit_all_deam | Tianyuan | Anatolia_N | Mbuti | 0.0186 | 3.72 | 0.0050 | 21,337 | 20,558 | 447,451 | Middle east |
| Salkhit_all_deam | Tianyuan | Iraqi_Jew | Mbuti | 0.0158 | 2.91 | 0.0054 | 22,093 | 21,407 | 473,283 | Middle east |
| Salkhit_all_deam | Tianyuan | Palestinian | Mbuti | 0.013 | 2.61 | 0.0049 | 21,886 | 21,324 | 473,286 | Middle east |
| Salkhit_all_deam | Tianyuan | Yi | Mbuti | -0.0107 | -1.85 | 0.0057 | 22,245 | 22,728 | 473,276 | East Eurasia |
| Salkhit_all_deam | Tianyuan | She | Mbuti | -0.0072 | -1.21 | 0.0059 | 22,312 | 22,635 | 473,281 | East Eurasia |
| Salkhit_all_deam | Tianyuan | Mongola | Mbuti | -0.0033 | -0.57 | 0.0057 | 22,406 | 22,554 | 473,279 | East Eurasia |
| Salkhit_all_deam | Tianyuan | Han | Mbuti | -0.0111 | -2.03 | 0.0054 | 22,252 | 22,751 | 473,298 | East Eurasia |
| Salkhit_all_deam | Tianyuan | Cambodian | Mbuti | -0.008 | -1.37 | 0.0058 | 22,199 | 22,555 | 473,280 | East Eurasia |
| Salkhit_all_deam | Tianyuan | Korean | Mbuti | -0.0081 | -1.39 | 0.0058 | 22,285 | 22,648 | 473,275 | East Eurasia |
| Salkhit_all_deam | Tianyuan | Japanese | Mbuti | -0.0112 | -2.01 | 0.0055 | 22,208 | 22,710 | 473,291 | East Eurasia |
| Salkhit_all_deam | Tianyuan | Thai | Mbuti | -0.0076 | -1.36 | 0.0055 | 22,225 | 22,567 | 473,274 | East Eurasia |
| Salkhit_all_deam | Tianyuan | Uygur | Mbuti | 0.0012 | 0.21 | 0.0056 | 22,220 | 22,167 | 473,286 | East Eurasia |
| Salkhit_all_deam | Tianyuan | Tibetan | Mbuti | -0.0092 | -1.62 | 0.0056 | 22,141 | 22,553 | 473,270 | East Eurasia |
| Salkhit_all_deam | Tianyuan | Dusun | Mbuti | -0.0066 | -1.15 | 0.0057 | 22,264 | 22,559 | 473,272 | East Eurasia |
| Salkhit_all_deam | Tianyuan | Igorot | Mbuti | -0.0147 | -2.35 | 0.0062 | 22,041 | 22,698 | 473,251 | East Eurasia |
| Salkhit_all_deam | Tianyuan | Hawaiian | Mbuti | -0.0064 | -0.95 | 0.0067 | 22,275 | 22,560 | 473,091 | Oceania |
| Salkhit_all_deam | Tianyuan | Papuan | Mbuti | -0.0029 | -0.49 | 0.0058 | 22,072 | 22,199 | 473,297 | Oceania |
| Salkhit_all_deam | Tianyuan | Bougainville | Mbuti | -0.0024 | -0.39 | 0.0061 | 22,052 | 22,158 | 473,259 | Oceania |
| Salkhit_all_deam | Tianyuan | Australian | Mbuti | -0.0123 | -2.04 | 0.0060 | 21,851 | 22,397 | 473,287 | Oceania |
| Salkhit_all_deam | Tianyuan | Maori | Mbuti | 0.0059 | 0.97 | 0.0060 | 22,251 | 21,990 | 473,083 | Oceania |
| Salkhit_all_deam | Tianyuan | Mayan | Mbuti | 0.0095 | 1.55 | 0.0061 | 22,689 | 22,264 | 473,266 | Native American |
| Salkhit_all_deam | Tianyuan | Piapoco | Mbuti | 0.0098 | 1.58 | 0.0062 | 22,696 | 22,253 | 473,262 | Native American |
| Salkhit_all_deam | Tianyuan | Chipewyan | Mbuti | -0.0026 | -0.44 | 0.0059 | 22,498 | 22,613 | 473,263 | Native American |
| Salkhit_all_deam | Tianyuan | Surui | Mbuti | -0.0037 | -0.58 | 0.0063 | 22,397 | 22,564 | 473,249 | Native American |
| Salkhit_all_deam | Tianyuan | Zapotec | Mbuti | 0.007 | 1.14 | 0.0061 | 22,588 | 22,274 | 473,271 | Native American |
| Salkhit_all_deam | Tianyuan | Quechua | Mbuti | 0.0062 | 1.05 | 0.0059 | 22,585 | 22,307 | 473,292 | Native American |
| Salkhit_all_deam | Tianyuan | Karitiana | Mbuti | 0.0012 | 0.2 | 0.006 | 22,516 | 22,460 | 473,282 | Native American |
| Salkhit_all_deam | Tianyuan | Chane | Mbuti | -0.0063 | -0.97 | 0.0065 | 22,331 | 22,616 | 473,172 | Native American |
| Salkhit_all_deam | Tianyuan | Nahua | Mbuti | 0.0043 | 0.64 | 0.0067 | 22,488 | 22,297 | 473,103 | Native American |
| Salkhit_all_deam | Tianyuan | Mixe | Mbuti | 0.0078 | 1.27 | 0.0061 | 22,602 | 22,252 | 473,278 | Native American |
| Salkhit_all_deam | Tianyuan | Pima | Mbuti | 0 | -0.00 | 0 | 22,389 | 22,390 | 473,257 | Native American |
| Salkhit_all_deam | Tianyuan | Clovis | Mbuti | -0.0014 | -0.20 | 0.0068 | 22,253 | 22,317 | 472,460 | Native American |
| Salkhit_all_deam | Tianyuan | Cree | Mbuti | 0.0025 | 0.43 | 0.0057 | 22,326 | 22,215 | 473,275 | Native American |
| Salkhit_all_deam | Tianyuan | Mixtec | Mbuti | 0.008 | 1.35 | 0.0059 | 22,444 | 22,089 | 473,277 | Native American |
| Salkhit_all_deam | Tianyuan | Botocudo | Mbuti | -0.0091 | -1.36 | 0.0066 | 21,331 | 21,722 | 455,920 | Native American |

**Table S11** *D-statistics* of the form *D(Salkhit, Tianyuan, X, Mbuti)* where X is an ancient individual or present-day population (Pop 3). Significant *D*-statistics (|Z| > 2) are in red.

| Source 2 | Outgroup | Country | nbr_ind | ancient | Salkhit_all_deam_f3 | std.err | Z | SNPs | Tianyuan_f3 | std.err | Z | SNPs | GoyetQ116_f3 | std.err | Z | SNPs | Yana1_f3 | std.err | Z | SNPs |
|---|---|---|---|---|---|---|---|---|---|---|---|---|---|---|---|---|---|---|---|---|
| Salkhit_all_deam | Mbuti | Mongolia | 1 | ancient | - | - | - | - | 0.2569 | 0.0037 | 69.34 | 453,078 | 0.2438 | 0.0038 | 63.46 | 302,108 | 0.2515 | 0.0038 | 65.41 | 515,611 |
| Salkhit_bl_all_deam | Mbuti | Mongolia | 1 | ancient | 0.6840 | 0.0042 | 163.64 | 223,909 | 0.2551 | 0.0039 | 66.05 | 198,777 | 0.2396 | 0.0041 | 58.03 | 131,653 | 0.251 | 0.0040 | 62.47 | 225,438 |
| Salkhit_all | Mbuti | Mongolia | 1 | ancient | 0.5112 | 0.0038 | 135.16 | 352,655 | 0.2474 | 0.0033 | 74.62 | 880,445 | 0.2416 | 0.0034 | 70.76 | 538,250 | 0.2454 | 0.0034 | 72.27 | 1,106,919 |
| Tianyuan | Mbuti | China | 1 | ancient | 0.2577 | 0.0038 | 68.01 | 309,508 | - | - | - | - | 0.234 | 0.0038 | 62.09 | 494,218 | 0.2306 | 0.0037 | 62.99 | 892,831 |
| Yana1 | Mbuti | Russia | 1 | ancient | 0.2532 | 0.0039 | 64.52 | 347,749 | 0.2306 | 0.0037 | 62.99 | 892,831 | 0.2595 | 0.0039 | 66.69 | 524,527 | - | - | - | - |
| Yana2 | Mbuti | Russia | 1 | ancient | 0.2514 | 0.0038 | 66.17 | 346,701 | 0.2309 | 0.0035 | 65.83 | 889,915 | 0.2535 | 0.0038 | 65.88 | 523,209 | 0.2875 | 0.0041 | 70.71 | 1,257,132 |
| GoyetQ116-1 | Mbuti | France | 1 | ancient | 0.2444 | 0.0039 | 62.06 | 209,974 | 0.234 | 0.0038 | 62.09 | 494,218 | - | - | - | - | 0.2595 | 0.0039 | 66.69 | 524,527 |
| Kolyma1 | Mbuti | Russia | 1 | ancient | 0.2434 | 0.0036 | 67.31 | 348,341 | 0.2407 | 0.0035 | 68.39 | 892,283 | 0.2348 | 0.0035 | 67.23 | 526,890 | 0.2365 | 0.0036 | 65.36 | 1,268,209 |
| Malta1 | Mbuti | Russia | 1 | ancient | 0.2361 | 0.0038 | 61.49 | 261,339 | 0.2248 | 0.0035 | 64.66 | 667,974 | 0.2570 | 0.0039 | 66.17 | 402,418 | 0.2482 | 0.0037 | 66.55 | 869,938 |
| Vestonice13 | Mbuti | Italy | 1 | ancient | 0.2361 | 0.0061 | 38.37 | 37,208 | 0.2197 | 0.0048 | 46.16 | 86,379 | 0.2707 | 0.0051 | 52.25 | 68,976 | 0.2446 | 0.0046 | 53.57 | 86,734 |
| El Miron | Mbuti | Spain | 1 | ancient | 0.2341 | 0.0038 | 61.04 | 200,050 | 0.2230 | 0.0035 | 62.68 | 484,386 | 0.2983 | 0.0040 | 74.25 | 368,747 | 0.2477 | 0.0038 | 64.46 | 493,540 |
| Vestonice16 | Mbuti | Italy | 1 | ancient | 0.2337 | 0.0038 | 61.62 | 228 ,490 | 0.2215 | 0.0034 | 64.92 | 559,371 | 0.2673 | 0.0038 | 69.74 | 421,805 | 0.2493 | 0.0036 | 68.36 | 579,486 |
| Sunghir3 | Mbuti | Russia | 1 | ancient | 0.2302 | 0.0037 | 62.79 | 348,667 | 0.2199 | 0.0035 | 63.05 | 894,365 | 0.2623 | 0.0039 | 67.79 | 524,833 | 0.2436 | 0.0035 | 69.5 | 1,267,331 |
| Kostenki14 | Mbuti | Russia | 1 | ancient | 0.2282 | 0.0038 | 59.14 | 336 ,520 | 0.2171 | 0.0035 | 62.46 | 875,490 | 0.2646 | 0.0039 | 67.16 | 534,553 | 0.2455 | 0.0035 | 70.2 | 1,042,124 |
| UstIshim | Mbuti | Russia | 1 | ancient | 0.2171 | 0.0034 | 63.15 | 359,758 | 0.2176 | 0.0035 | 62.36 | 921,445 | 0.2312 | 0.0037 | 62.68 | 544,563 | 0.2211 | 0.0036 | 61.88 | 1,303,928 |
| Oase1 | Mbuti | Romania | 1 | ancient | 0.2161 | 0.0046 | 47.03 | 69,239 | 0.2108 | 0.0039 | 53.45 | 166,081 | 0.2139 | 0.0042 | 50.34 | 117,684 | 0.2053 | 0.0038 | 54.49 | 171,728 |

| Source 2 | Outgroup | Country | nbr_ind | ancient | Malta_f3 | std.err | Z | SNPs | Kostenki14_f3 | std.err | Z | SNPs | Sunghir3_f3 | std.err | Z | SNPs | UstIshim_f3 | std.err | Z | SNPs |
|---|---|---|---|---|---|---|---|---|---|---|---|---|---|---|---|---|---|---|---|---|
| Salkhit_all_deam | Mbuti | Mongolia | 1 | ancient | 0.2348 | 0.0037 | 62.96 | 385,846 | 0.2278 | 0.0038 | 60.03 | 496,864 | 0.2302 | 0.0037 | 62.8 | 348,667 | 0.2177 | 0.0034 | 62.91 | 533,296 |
| Salkhit_bl_all_deam | Mbuti | Mongolia | 1 | ancient | 0.234 | 0.004 | 58.68 | 168,887 | 0.2262 | 0.0039 | 57.48 | 216,414 | 0.2269 | 0.0037 | 61.09 | 226,058 | 0.2148 | 0.0035 | 60.91 | 233,364 |
| Salkhit_all | Mbuti | Mongolia | 1 | ancient | 0.2357 | 0.0033 | 70.27 | 812,952 | 0.2284 | 0.0033 | 68.23 | 1,019,116 | 0.2312 | 0.0032 | 73.08 | 1,118,728 | 0.2166 | 0.0031 | 69.08 | 1,143,318 |
| Tianyuan | Mbuti | China | 1 | ancient | 0.2248 | 0.0035 | 64.66 | 667,974 | 0.2171 | 0.00348 | 62.46 | 875,490 | 0.22 | 0.0035 | 63.05 | 894,365 | 0.2176 | 0.0035 | 62.36 | 921,445 |
| Yana1 | Mbuti | Russia | 1 | ancient | 0.2482 | 0.0037 | 66.55 | 869,938 | 0.2455 | 0.0035 | 70.2 | 1,042,124 | 0.2436 | 0.0035 | 69.50 | 1,267,331 | 0.2211 | 0.0036 | 61.88 | 1,303,928 |
| Yana2 | Mbuti | Russia | 1 | ancient | 0.2489 | 0.0037 | 67.21 | 866,900 | 0.2459 | 0.0035 | 70.28 | 1,038,817 | 0.2366 | 0.0035 | 68.04 | 1,263,654 | 0.2170 | 0.0035 | 61.97 | 1,300,564 |
| GoyetQ116-1 | Mbuti | France | 1 | ancient | 0.2571 | 0.0039 | 66.17 | 402,418 | 0.2646 | 0.0039 | 67.16 | 534,553 | 0.2623 | 0.0039 | 67.79 | 524,833 | 0.2312 | 0.0037 | 62.68 | 544,563 |
| Kolyma1 | Mbuti | Russia | 1 | ancient | 0.249 | 0.0037 | 67.59 | 870,082 | 0.224 | 0.0034 | 65.67 | 1,044,865 | 0.2224 | 0.0033 | 66.99 | 1,270,026 | 0.22 | 0.0034 | 63.74 | 1,304,421 |
| Malta1 | Mbuti | Russia | 1 | ancient | - | - | - | - | 0.2449 | 0.0037 | 66.89 | 775,359 | 0.241 | 0.0036 | 67.04 | 870,863 | 0.2198 | 0.0035 | 62.34 | 895,611 |
| Vestonice13 | Mbuti | Italy | 1 | ancient | 0.2504 | 0.0048 | 51.69 | 66,880 | 0.2704 | 0.0048 | 56.33 | 85,658 | 0.2806 | 0.0048 | 58.48 | 86,325 | 0.2213 | 0.0041 | 54.17 | 89,927 |
| ElMiron | Mbuti | Spain | 1 | ancient | 0.2485 | 0.0037 | 66.82 | 377,596 | 0.2572 | 0.0036 | 70.36 | 497,369 | 0.2582 | 0.0037 | 69.78 | 493,122 | 0.2215 | 0.0035 | 62.89 | 510,882 |
| Vestonice16 | Mbuti | Italy | 1 | ancient | 0.2521 | 0.0038 | 66.17 | 441,337 | 0.2676 | 0.0037 | 72.35 | 586,521 | 0.2822 | 0.0039 | 72.86 | 578,055 | 0.2241 | 0.0034 | 65.28 | 600,410 |
| Sunghir3 | Mbuti | Russia | 1 | ancient | 0.2409 | 0.0036 | 67.04 | 870,863 | 0.2692 | 0.0038 | 71.49 | 1,039,793 | - | - | - | - | 0.2181 | 0.0034 | 63.43 | 1,304,231 |
| Kostenki14 | Mbuti | Russia | 1 | ancient | 0.2449 | 0.0037 | 66.89 | 775,359 | - | - | - | - | 0.2692 | 0.0038 | 71.49 | 1,039,793 | 0.2201 | 0.0035 | 61.97 | 1,076,371 |
| UstIshim | Mbuti | Russia | 1 | ancient | 0.2198 | 0.0035 | 62.34 | 895,611 | 0.2201 | 0.0035 | 61.97 | 1,076,371 | 0.2181 | 0.0034 | 63.43 | 1,304,231 | - | - | - | - |
| Oase1 | Mbuti | Romania | 1 | ancient | 0.2083 | 0.0039 | 53.72 | 131,286 | 0.2071 | 0.0037 | 55.73 | 168,029 | 0.2029 | 0.0039 | 51.45 | 172,007 | 0.2031 | 0.0039 | 51.96 | 177,532 |

**Table S12** Shared genetic drift between ancient modern humans estimated using the *f3*-statistic.

| Pop1 | Pop2 | Pop3 | Pop4 | D-Stat | Z | std.err | BABA | ABBA | #SNPs | Pop3 - origin |
|---|---|---|---|---|---|---|---|---|---|---|
| Salkhit_all_deam | Yana1 | Tianyuan | Mbuti | 0.0593 | 8.32 | 0.0071 | 22,863 | 20,304 | 454,951 | ancient |
| Salkhit_all_deam | Yana2 | Tianyuan | Mbuti | 0.0619 | 8.63 | 0.0071 | 22,906 | 20,234 | 454,623 | ancient |
| Salkhit_all_deam | Kolyma1 | Tianyuan | Mbuti | 0.0378 | 5.13 | 0.0073 | 22,969 | 21,297 | 454,916 | ancient |
| Salkhit_all_deam | Malta1 | Tianyuan | Mbuti | 0.0718 | 9.36 | 0.0076 | 18,331 | 15,876 | 342,502 | ancient |
| Salkhit_all_deam | Sunghir3 | Tianyuan | Mbuti | 0.0866 | 11.51 | 0.0075 | 24,460 | 20,563 | 454,900 | ancient |
| Salkhit_all_deam | GoyetQ116 | Tianyuan | Mbuti | 0.0703 | 8.81 | 0.0079 | 16,236 | 14,103 | 283,946 | ancient |
| Salkhit_all_deam | UstIshim | Tianyuan | Mbuti | 0.0875 | 11.32 | 0.0077 | 25,081 | 21,046 | 454,500 | ancient |
| Salkhit_all_deam | Kostenki14 | Tianyuan | Mbuti | 0.0941 | 12.37 | 0.0076 | 24,346 | 20,158 | 443,299 | ancient |
| Salkhit_all_deam | Vestonice16 | Tianyuan | Mbuti | 0.0905 | 11.76 | 0.0076 | 18,084 | 15,084 | 322,349 | ancient |
| Salkhit_all_deam | ElMiron | Tianyuan | Mbuti | 0.0842 | 10.46 | 0.0080 | 15,881 | 13,416 | 285,670 | ancient |
| Salkhit_all_deam | Saami | Tianyuan | Mbuti | 0.0786 | 12.41 | 0.0063 | 25,714 | 21,966 | 473,287 | West Eurasia |
| Salkhit_all_deam | French | Tianyuan | Mbuti | 0.1042 | 17.02 | 0.0061 | 26,585 | 21,568 | 473,297 | West Eurasia |
| Salkhit_all_deam | Russian | Tianyuan | Mbuti | 0.0927 | 14.92 | 0.0062 | 26,273 | 21,814 | 473,271 | West Eurasia |
| Salkhit_all_deam | Estonian | Tianyuan | Mbuti | 0.1 | 15.87 | 0.0063 | 26,408 | 21,607 | 473,279 | West Eurasia |
| Salkhit_all_deam | Icelandic | Tianyuan | Mbuti | 0.0985 | 15.69 | 0.0062 | 26,426 | 21,687 | 473,276 | West Eurasia |
| Salkhit_all_deam | Finnish | Tianyuan | Mbuti | 0.1014 | 16.09 | 0.0063 | 26,520 | 21,636 | 473,292 | West Eurasia |
| Salkhit_all_deam | Czech | Tianyuan | Mbuti | 0.1062 | 15.55 | 0.0068 | 26,627 | 21,513 | 473,148 | West Eurasia |
| Salkhit_all_deam | Basque | Tianyuan | Mbuti | 0.109 | 17.49 | 0.0062 | 26,747 | 21,489 | 473,273 | West Eurasia |
| Salkhit_all_deam | Spanish | Tianyuan | Mbuti | 0.1144 | 18.19 | 0.0062 | 26,990 | 21,448 | 473,273 | West Eurasia |
| Salkhit_all_deam | Sardinian | Tianyuan | Mbuti | 0.1121 | 18.1 | 0.0061 | 27,033 | 21,585 | 473,291 | West Eurasia |
| Salkhit_all_deam | English | Tianyuan | Mbuti | 0.1075 | 16.9 | 0.0063 | 26,766 | 21,571 | 473,267 | West Eurasia |
| Salkhit_all_deam | Anatolia_N | Tianyuan | Mbuti | 0.1102 | 18.36 | 0.0060 | 25,650 | 20,558 | 447,451 | Middle east |
| Salkhit_all_deam | Iraqi_Jew | Tianyuan | Mbuti | 0.1201 | 18.73 | 0.0064 | 27,251 | 21,407 | 473,283 | Middle east |
| Salkhit_all_deam | Palestinian | Tianyuan | Mbuti | 0.1335 | 22.50 | 0.0059 | 27,892 | 21,324 | 473,286 | Middle east |
| Salkhit_all_deam | Yi | Tianyuan | Mbuti | 0.0243 | 3.82 | 0.0063 | 23,862 | 22,728 | 473,276 | East Eurasia |
| Salkhit_all_deam | She | Tianyuan | Mbuti | 0.0286 | 4.41 | 0.0064 | 23,966 | 22,635 | 473,281 | East Eurasia |
| Salkhit_all_deam | Mongola | Tianyuan | Mbuti | 0.0323 | 4.80 | 0.0067 | 24,059 | 22,554 | 473,279 | East Eurasia |
| Salkhit_all_deam | Han | Tianyuan | Mbuti | 0.0269 | 4.24 | 0.0063 | 24,009 | 22,751 | 473,298 | East Eurasia |
| Salkhit_all_deam | Cambodian | Tianyuan | Mbuti | 0.0402 | 6.24 | 0.0064 | 24,446 | 22,555 | 473,280 | East Eurasia |
| Salkhit_all_deam | Korean | Tianyuan | Mbuti | 0.0301 | 4.64 | 0.0064 | 24,052 | 22,648 | 473,275 | East Eurasia |
| Salkhit_all_deam | Japanese | Tianyuan | Mbuti | 0.0284 | 4.51 | 0.0062 | 24,038 | 22,710 | 473,291 | East Eurasia |
| Salkhit_all_deam | Thai | Tianyuan | Mbuti | 0.0357 | 5.69 | 0.0062 | 24,240 | 22,567 | 473,274 | East Eurasia |
| Salkhit_all_deam | Uygur | Tianyuan | Mbuti | 0.0682 | 11.17 | 0.0061 | 25,411 | 22,167 | 473,286 | East Eurasia |
| Salkhit_all_deam | Tibetan | Tianyuan | Mbuti | 0.035 | 5.51 | 0.0063 | 24,191 | 22,553 | 473,270 | East Eurasia |
| Salkhit_all_deam | Dusun | Tianyuan | Mbuti | 0.0353 | 5.36 | 0.0065 | 24,211 | 22,559 | 473,272 | East Eurasia |
| Salkhit_all_deam | Igorot | Tianyuan | Mbuti | 0.0293 | 4.28 | 0.0068 | 24,070 | 22,698 | 473,251 | East Eurasia |
| Salkhit_all_deam | Hawaiian | Tianyuan | Mbuti | 0.0423 | 5.85 | 0.0072 | 24,551 | 22,560 | 473,091 | Oceania |
| Salkhit_all_deam | Papuan | Tianyuan | Mbuti | 0.0594 | 8.98 | 0.0066 | 25,004 | 22,199 | 473,297 | Oceania |
| Salkhit_all_deam | Bougainville | Tianyuan | Mbuti | 0.0595 | 8.71 | 0.0068 | 24,964 | 22,158 | 473,259 | Oceania |
| Salkhit_all_deam | Australian | Tianyuan | Mbuti | 0.0568 | 8.42 | 0.0067 | 25,093 | 22,397 | 473,287 | Oceania |
| Salkhit_all_deam | Maori | Tianyuan | Mbuti | 0.0815 | 11.83 | 0.0068 | 25,892 | 21,990 | 473,083 | Oceania |
| Salkhit_all_deam | Mayan | Tianyuan | Mbuti | 0.044 | 6.43 | 0.0068 | 24,311 | 22,264 | 473,266 | Native American |
| Salkhit_all_deam | Piapoco | Tianyuan | Mbuti | 0.0476 | 7.06 | 0.0067 | 24,477 | 22,253 | 473,262 | Native American |
| Salkhit_all_deam | Chipewyan | Tianyuan | Mbuti | 0.0363 | 5.48 | 0.0066 | 24,315 | 22,613 | 473,263 | Native American |
| Salkhit_all_deam | Surui | Tianyuan | Mbuti | 0.0353 | 5.11 | 0.0069 | 24,216 | 22,564 | 473,249 | Native American |
| Salkhit_all_deam | Zapotec | Tianyuan | Mbuti | 0.0463 | 6.94 | 0.0066 | 24,437 | 22,274 | 473,271 | Native American |
| Salkhit_all_deam | Quechua | Tianyuan | Mbuti | 0.0463 | 7.10 | 0.0065 | 24,474 | 22,307 | 473,292 | Native American |
| Salkhit_all_deam | Karitiana | Tianyuan | Mbuti | 0.0421 | 6.17 | 0.0068 | 24,434 | 22,460 | 473,282 | Native American |
| Salkhit_all_deam | Chane | Tianyuan | Mbuti | 0.0362 | 5.01 | 0.0072 | 24,315 | 22,616 | 473,172 | Native American |
| Salkhit_all_deam | Nahua | Tianyuan | Mbuti | 0.0494 | 6.72 | 0.0073 | 24,615 | 22,297 | 473,103 | Native American |
| Salkhit_all_deam | Mixe | Tianyuan | Mbuti | 0.0508 | 7.55 | 0.0067 | 24,632 | 22,252 | 473,278 | Native American |
| Salkhit_all_deam | Pima | Tianyuan | Mbuti | 0.0466 | 6.96 | 0.0066 | 24,579 | 22,390 | 473,257 | Native American |
| Salkhit_all_deam | Clovis | Tianyuan | Mbuti | 0.0435 | 5.72 | 0.0076 | 24,348 | 22,317 | 472,460 | Native American |
| Salkhit_all_deam | Cree | Tianyuan | Mbuti | 0.0582 | 9.16 | 0.0063 | 24,959 | 22,215 | 473,275 | Native American |
| Salkhit_all_deam | Mixtec | Tianyuan | Mbuti | 0.0638 | 9.82 | 0.0064 | 25,101 | 22,089 | 473,277 | Native American |
| Salkhit_all_deam | Botocudo | Tianyuan | Mbuti | 0.0372 | 5.15 | 0.0072 | 23,401 | 21,722 | 455,920 | Native American |

**Table S13** *D*-statistics of the form *D(Salkhit, X, Tianyuan, Mbuti)* where X is an ancient individual or present-day population (Pop 3). Significant *D*-statistics (|Z| > 2) are in red.

| Pop1 | Pop2 | Pop3 | Pop4 | D-Stat | Z | std.err | BABA | ABBA | #SNPs | Pop3 - origin |
|---|---|---|---|---|---|---|---|---|---|---|
| Tianyuan | Yana1 | Salkhit_all_deam | Mbuti | 0.0092 | 1.18 | 0.0078 | 22,863 | 22,447 | 454,951 | ancient |
| Tianyuan | Yana2 | Salkhit_all_deam | Mbuti | 0.0159 | 2.04 | 0.0078 | 22,906 | 22,189 | 454,623 | ancient |
| Tianyuan | Kolyma1 | Salkhit_all_deam | Mbuti | 0.0341 | 4.84 | 0.0070 | 22,969 | 21,452 | 454,916 | ancient |
| Tianyuan | Malta1 | Salkhit_all_deam | Mbuti | 0.0491 | 6.48 | 0.0075 | 18,331 | 16,614 | 342,502 | ancient |
| Tianyuan | Sunghir3 | Salkhit_all_deam | Mbuti | 0.0621 | 8.2 | 0.0075 | 24,460 | 21,602 | 454,900 | ancient |
| Tianyuan | GoyetQ116 | Salkhit_all_deam | Mbuti | 0.0466 | 5.95 | 0.0078 | 16,236 | 14,790 | 283,946 | ancient |
| Tianyuan | UstIshim | Salkhit_all_deam | Mbuti | 0.0926 | 12.63 | 0.0073 | 25,081 | 20,828 | 454,500 | ancient |
| Tianyuan | Kostenki14 | Salkhit_all_deam | Mbuti | 0.0679 | 8.45 | 0.0080 | 24,346 | 21,249 | 443,299 | ancient |
| Tianyuan | Vestonice16 | Salkhit_all_deam | Mbuti | 0.0613 | 7.71 | 0.0079 | 18,084 | 15,994 | 322,349 | ancient |
| Tianyuan | ElMiron | Salkhit_all_deam | Mbuti | 0.0594 | 7.46 | 0.0079 | 15,881 | 14,101 | 285,670 | ancient |
| Tianyuan | Saami | Salkhit_all_deam | Mbuti | 0.0668 | 10.42 | 0.0064 | 25,714 | 22,494 | 473,287 | West Eurasia |
| Tianyuan | French | Salkhit_all_deam | Mbuti | 0.0841 | 13.53 | 0.0062 | 26,585 | 22,460 | 473,297 | West Eurasia |
| Tianyuan | Russian | Salkhit_all_deam | Mbuti | 0.0811 | 13.05 | 0.0062 | 26,273 | 22,331 | 473,271 | West Eurasia |
| Tianyuan | Estonian | Salkhit_all_deam | Mbuti | 0.0789 | 12.65 | 0.0062 | 26,408 | 22,545 | 473,279 | West Eurasia |
| Tianyuan | Icelandic | Salkhit_all_deam | Mbuti | 0.0809 | 12.73 | 0.0063 | 26,426 | 22,471 | 473,276 | West Eurasia |
| Tianyuan | Finnish | Salkhit_all_deam | Mbuti | 0.0817 | 13.03 | 0.0062 | 26,520 | 22,513 | 473,292 | West Eurasia |
| Tianyuan | Czech | Salkhit_all_deam | Mbuti | 0.0832 | 12.01 | 0.0069 | 26,627 | 22,539 | 473,148 | West Eurasia |
| Tianyuan | Basque | Salkhit_all_deam | Mbuti | 0.0888 | 14.13 | 0.0062 | 26,747 | 22,384 | 473,273 | West Eurasia |
| Tianyuan | Spanish | Salkhit_all_deam | Mbuti | 0.0939 | 14.57 | 0.0064 | 26,990 | 22,356 | 473,273 | West Eurasia |
| Tianyuan | Sardinian | Salkhit_all_deam | Mbuti | 0.0977 | 15.60 | 0.0062 | 27,033 | 22,223 | 473,291 | West Eurasia |
| Tianyuan | English | Salkhit_all_deam | Mbuti | 0.0888 | 14.03 | 0.0063 | 26,766 | 22,402 | 473,267 | West Eurasia |
| Tianyuan | Anatolia_N | Salkhit_all_deam | Mbuti | 0.0918 | 15.04 | 0.0061 | 25,650 | 21,337 | 447,451 | Middle east |
| Tianyuan | Iraqi_Jew | Salkhit_all_deam | Mbuti | 0.1045 | 16.58 | 0.0063 | 27,251 | 22,093 | 473,283 | Middle east |
| Tianyuan | Palestinian | Salkhit_all_deam | Mbuti | 0.1207 | 20.25 | 0.0059 | 27,892 | 21,886 | 473,286 | Middle east |
| Tianyuan | Yi | Salkhit_all_deam | Mbuti | 0.0351 | 5.70 | 0.0061 | 23,862 | 22,245 | 473,276 | East Eurasia |
| Tianyuan | She | Salkhit_all_deam | Mbuti | 0.0357 | 5.81 | 0.0061 | 23,966 | 22,312 | 473,281 | East Eurasia |
| Tianyuan | Mongola | Salkhit_all_deam | Mbuti | 0.0356 | 5.95 | 0.0059 | 24,059 | 22,406 | 473,279 | East Eurasia |
| Tianyuan | Han | Salkhit_all_deam | Mbuti | 0.038 | 6.32 | 0.0060 | 24,009 | 22,252 | 473,298 | East Eurasia |
| Tianyuan | Cambodian | Salkhit_all_deam | Mbuti | 0.0482 | 7.91 | 0.0061 | 24,446 | 22,199 | 473,280 | East Eurasia |
| Tianyuan | Korean | Salkhit_all_deam | Mbuti | 0.0381 | 6.17 | 0.0061 | 24,052 | 22,285 | 473,275 | East Eurasia |
| Tianyuan | Japanese | Salkhit_all_deam | Mbuti | 0.0396 | 6.51 | 0.0061 | 24,038 | 22,208 | 473,291 | East Eurasia |
| Tianyuan | Thai | Salkhit_all_deam | Mbuti | 0.0434 | 7.08 | 0.0061 | 24,240 | 22,225 | 473,274 | East Eurasia |
| Tianyuan | Uygur | Salkhit_all_deam | Mbuti | 0.067 | 11.23 | 0.0059 | 25,411 | 22,220 | 473,286 | East Eurasia |
| Tianyuan | Tibetan | Salkhit_all_deam | Mbuti | 0.0442 | 7.17 | 0.0062 | 24,191 | 22,141 | 473,270 | East Eurasia |
| Tianyuan | Dusun | Salkhit_all_deam | Mbuti | 0.0419 | 6.68 | 0.0063 | 24,211 | 22,264 | 473,272 | East Eurasia |
| Tianyuan | Igorot | Salkhit_all_deam | Mbuti | 0.044 | 6.9 | 0.0064 | 24,070 | 22,041 | 473,251 | East Eurasia |
| Tianyuan | Hawaiian | Salkhit_all_deam | Mbuti | 0.0486 | 7.25 | 0.0067 | 24,551 | 22,275 | 473,091 | Oceania |
| Tianyuan | Papuan | Salkhit_all_deam | Mbuti | 0.0623 | 9.71 | 0.0064 | 25,004 | 22,072 | 473,297 | Oceania |
| Tianyuan | Bougainville | Salkhit_all_deam | Mbuti | 0.0619 | 9.37 | 0.0066 | 24,964 | 22,052 | 473,259 | Oceania |
| Tianyuan | Australian | Salkhit_all_deam | Mbuti | 0.0691 | 10.29 | 0.0067 | 25,093 | 21,851 | 473,287 | Oceania |
| Tianyuan | Maori | Salkhit_all_deam | Mbuti | 0.0756 | 11.57 | 0.0065 | 25,892 | 22,251 | 473,083 | Oceania |
| Tianyuan | Mayan | Salkhit_all_deam | Mbuti | 0.0345 | 5.22 | 0.0066 | 24,311 | 22,689 | 473,266 | Native American |
| Tianyuan | Piapoco | Salkhit_all_deam | Mbuti | 0.0378 | 5.94 | 0.0064 | 24,477 | 22,696 | 473262 | Native American |
| Tianyuan | Chipewyan | Salkhit_all_deam | Mbuti | 0.0388 | 6.08 | 0.0064 | 24,315 | 22,498 | 473,263 | Native American |
| Tianyuan | Surui | Salkhit_all_deam | Mbuti | 0.039 | 5.95 | 0.0065 | 24,216 | 22,397 | 473,249 | Native American |
| Tianyuan | Zapotec | Salkhit_all_deam | Mbuti | 0.0393 | 6.25 | 0.0063 | 24,437 | 22,588 | 473,271 | Native American |
| Tianyuan | Quechua | Salkhit_all_deam | Mbuti | 0.0401 | 6.32 | 0.0063 | 24,474 | 22,585 | 473,292 | Native American |
| Tianyuan | Karitiana | Salkhit_all_deam | Mbuti | 0.0408 | 6.13 | 0.0066 | 24,434 | 22,516 | 473,282 | Native American |
| Tianyuan | Chane | Salkhit_all_deam | Mbuti | 0.0425 | 6.24 | 0.0068 | 24,315 | 22,331 | 473,172 | Native American |
| Tianyuan | Nahua | Salkhit_all_deam | Mbuti | 0.0452 | 6.53 | 0.0069 | 24,615 | 22,488 | 473,103 | Native American |
| Tianyuan | Mixe | Salkhit_all_deam | Mbuti | 0.043 | 6.68 | 0.0064 | 24,632 | 22,602 | 473,278 | Native American |
| Tianyuan | Pima | Salkhit_all_deam | Mbuti | 0.0466 | 7.11 | 0.0065 | 24,579 | 22,389 | 473,257 | Native American |
| Tianyuan | Clovis | Salkhit_all_deam | Mbuti | 0.045 | 6.22 | 0.0072 | 24,348 | 22,253 | 472,460 | Native American |
| Tianyuan | Cree | Salkhit_all_deam | Mbuti | 0.0557 | 8.96 | 0.0062 | 24,959 | 22,326 | 473,275 | Native American |
| Tianyuan | Mixtec | Salkhit_all_deam | Mbuti | 0.0559 | 8.87 | 0.0063 | 25,101 | 22,444 | 473,277 | Native American |
| Tianyuan | Botocudo | Salkhit_all_deam | Mbuti | 0.0463 | 7.02 | 0.0066 | 23,401 | 21,331 | 455,920 | Native American |

**Table S14** *D*-statistics of the form *D(Tianyuan, X, Salkhit, Mbuti)* where X is an ancient individual or present-day population (Pop 3). Significant *D*-statistics (|Z| > 2) are red.

| Pop1 | Pop2 | Pop3 | Pop4 | D-Stat | Z | std.err | BABA | ABBA | #SNPs | Pop3 - origin |
|---|---|---|---|---|---|---|---|---|---|---|
| GoyetQ116-1 | Yana1 | Salkhit_all_deam | Mbuti | -0.0401 | -4.83 | 0.0083 | 14,632 | 15,853 | 289,874 | ancient |
| GoyetQ116-1 | Yana2 | Salkhit_all_deam | Mbuti | -0.0295 | -3.72 | 0.0079 | 14,741 | 15,636 | 289,697 | ancient |
| GoyetQ116-1 | Kolyma1 | Salkhit_all_deam | Mbuti | -0.01 | -1.34 | 0.0074 | 15,727 | 16,046 | 289,860 | ancient |
| GoyetQ116-1 | Malta1 | Salkhit_all_deam | Mbuti | 0.0065 | 0.84 | 0.0078 | 11,842 | 11,690 | 222,837 | ancient |
| GoyetQ116-1 | Sunghir3 | Salkhit_all_deam | Mbuti | 0.0197 | 2.51 | 0.0078 | 15,295 | 14,703 | 289,849 | ancient |
| GoyetQ116-1 | Tianyuan | Salkhit_all_deam | Mbuti | -0.0466 | -5.95 | 0.0078 | 14,790 | 16,236 | 283,946 | ancient |
| GoyetQ116-1 | UstIshim | Salkhit_all_deam | Mbuti | 0.0484 | 6.37 | 0.0076 | 16,513 | 14,989 | 289,546 | ancient |
| GoyetQ116-1 | Kostenki14 | Salkhit_all_deam | Mbuti | 0.0285 | 3.74 | 0.0076 | 15,318 | 14,471 | 287,616 | ancient |
| GoyetQ116-1 | Vestonice16 | Salkhit_all_deam | Mbuti | 0.0211 | 2.56 | 0.0082 | 13,026 | 12,488 | 249,877 | ancient |
| GoyetQ116-1 | ElMiron | Salkhit_all_deam | Mbuti | 0.0146 | 1.84 | 0.0079 | 10,852 | 10,540 | 224,131 | ancient |
| GoyetQ116-1 | Saami | Salkhit_all_deam | Mbuti | 0.0249 | 3.86 | 0.0064 | 16,963 | 16,140 | 305,481 | West Eurasia |
| GoyetQ116-1 | French | Salkhit_all_deam | Mbuti | 0.0437 | 7.28 | 0.0059 | 17,130 | 15,696 | 305,490 | West Eurasia |
| GoyetQ116-1 | Russian | Salkhit_all_deam | Mbuti | 0.039 | 6.18 | 0.0063 | 17,091 | 15,809 | 305,480 | West Eurasia |
| GoyetQ116-1 | Estonian | Salkhit_all_deam | Mbuti | 0.0387 | 6.22 | 0.0062 | 17,043 | 15,773 | 305,482 | West Eurasia |
| GoyetQ116-1 | Icelandic | Salkhit_all_deam | Mbuti | 0.0408 | 6.43 | 0.0063 | 16,995 | 15,663 | 305,479 | West Eurasia |
| GoyetQ116-1 | Finnish | Salkhit_all_deam | Mbuti | 0.0396 | 6.41 | 0.0061 | 17,111 | 15,808 | 305,484 | West Eurasia |
| GoyetQ116-1 | Czech | Salkhit_all_deam | Mbuti | 0.0424 | 6.04 | 0.0070 | 17,129 | 15,735 | 305,435 | West Eurasia |
| GoyetQ116-1 | Basque | Salkhit_all_deam | Mbuti | 0.0465 | 7.40 | 0.0062 | 17,153 | 15,630 | 305,479 | West Eurasia |
| GoyetQ116-1 | Spanish | Salkhit_all_deam | Mbuti | 0.054 | 8.63 | 0.0062 | 17,373 | 15,592 | 305,479 | West Eurasia |
| GoyetQ116-1 | Sardinian | Salkhit_all_deam | Mbuti | 0.0566 | 9.45 | 0.0059 | 17,357 | 15,498 | 305,484 | West Eurasia |
| GoyetQ116-1 | English | Salkhit_all_deam | Mbuti | 0.0464 | 7.43 | 0.0062 | 17,193 | 15,668 | 305,482 | West Eurasia |
| GoyetQ116-1 | Anatolia_N | Salkhit_all_deam | Mbuti | 0.0525 | 8.87 | 0.0059 | 17,237 | 15,518 | 305,057 | Middle east |
| GoyetQ116-1 | Iraqi_Jew | Salkhit_all_deam | Mbuti | 0.0633 | 9.98 | 0.0063 | 17,742 | 15,631 | 305,479 | Middle east |
| GoyetQ116-1 | Palestinian | Salkhit_all_deam | Mbuti | 0.0805 | 13.6 | 0.0059 | 18,129 | 15,429 | 305,482 | Middle east |
| GoyetQ116-1 | Yi | Salkhit_all_deam | Mbuti | -0.0118 | -1.82 | 0.0065 | 16,741 | 17,140 | 305,481 | East Eurasia |
| GoyetQ116-1 | She | Salkhit_all_deam | Mbuti | -0.0105 | -1.57 | 0.0067 | 16,832 | 17,189 | 305,483 | East Eurasia |
| GoyetQ116-1 | Mongola | Salkhit_all_deam | Mbuti | -0.0109 | -1.66 | 0.0066 | 16,781 | 17,151 | 305,479 | East Eurasia |
| GoyetQ116-1 | Han | Salkhit_all_deam | Mbuti | -0.0103 | -1.59 | 0.0065 | 16,834 | 17,183 | 305,488 | East Eurasia |
| GoyetQ116-1 | Cambodian | Salkhit_all_deam | Mbuti | 0.0029 | 0.45 | 0.0064 | 17,035 | 16,936 | 305,480 | East Eurasia |
| GoyetQ116-1 | Korean | Salkhit_all_deam | Mbuti | -0.009 | -1.33 | 0.0067 | 16,870 | 17,175 | 305,482 | East Eurasia |
| GoyetQ116-1 | Japanese | Salkhit_all_deam | Mbuti | -0.0071 | -1.10 | 0.0064 | 16,824 | 17,063 | 305,484 | East Eurasia |
| GoyetQ116-1 | Thai | Salkhit_all_deam | Mbuti | -0.0025 | -0.38 | 0.0066 | 16,988 | 17,075 | 305,480 | East Eurasia |
| GoyetQ116-1 | Uygur | Salkhit_all_deam | Mbuti | 0.0203 | 3.29 | 0.0068 | 17,132 | 16,449 | 305,484 | East Eurasia |
| GoyetQ116-1 | Tibetan | Salkhit_all_deam | Mbuti | -0.0012 | -0.18 | 0.0065 | 16,949 | 16,990 | 305,477 | East Eurasia |
| GoyetQ116-1 | Dusun | Salkhit_all_deam | Mbuti | -0.0057 | -0.85 | 0.0067 | 16,928 | 17,122 | 305,478 | East Eurasia |
| GoyetQ116-1 | Igorot | Salkhit_all_deam | Mbuti | -0.0021 | -0.31 | 0.0067 | 16,950 | 17,021 | 305,476 | East Eurasia |
| GoyetQ116-1 | Hawaiian | Salkhit_all_deam | Mbuti | 0 | -0.00 | 0 | 17,060 | 17,061 | 305,410 | Oceania |
| GoyetQ116-1 | Papuan | Salkhit_all_deam | Mbuti | 0.0168 | 2.57 | 0.0065 | 17,369 | 16,796 | 305,489 | Oceania |
| GoyetQ116-1 | Bougainville | Salkhit_all_deam | Mbuti | 0.0155 | 2.26 | 0.0069 | 17,306 | 16,778 | 305,468 | Oceania |
| GoyetQ116-1 | Australian | Salkhit_all_deam | Mbuti | 0.0222 | 3.4 | 0.0065 | 17,356 | 16,604 | 305,483 | Oceania |
| GoyetQ116-1 | Maori | Salkhit_all_deam | Mbuti | 0.0313 | 4.52 | 0.0069 | 17,222 | 16,178 | 305,419 | Oceania |
| GoyetQ116-1 | Mayan | Salkhit_all_deam | Mbuti | -0.0132 | -2.06 | 0.0064 | 16,577 | 17,020 | 305,473 | Native American |
| GoyetQ116-1 | Piapoco | Salkhit_all_deam | Mbuti | -0.0091 | -1.36 | 0.0067 | 16,662 | 16,967 | 305,468 | Native American |
| GoyetQ116-1 | Chipewyan | Salkhit_all_deam | Mbuti | -0.0094 | -1.46 | 0.0064 | 16,589 | 16,904 | 305,472 | Native American |
| GoyetQ116-1 | Surui | Salkhit_all_deam | Mbuti | -0.0101 | -1.47 | 0.0069 | 16,572 | 16,911 | 305,461 | Native American |
| GoyetQ116-1 | Zapotec | Salkhit_all_deam | Mbuti | -0.0091 | -1.39 | 0.0065 | 16,592 | 16,897 | 305,474 | Native American |
| GoyetQ116-1 | Quechua | Salkhit_all_deam | Mbuti | -0.0085 | -1.33 | 0.0064 | 16,694 | 16,980 | 305,484 | Native American |
| GoyetQ116-1 | Karitiana | Salkhit_all_deam | Mbuti | -0.0065 | -0.99 | 0.0065 | 16,724 | 16,944 | 305,481 | Native American |
| GoyetQ116-1 | Chane | Salkhit_all_deam | Mbuti | -0.0069 | -0.97 | 0.0071 | 16,679 | 16,910 | 305,446 | Native American |
| GoyetQ116-1 | Nahua | Salkhit_all_deam | Mbuti | -0.0028 | -0.4 | 0.0070 | 16,791 | 16,884 | 305,416 | Native American |
| GoyetQ116-1 | Mixe | Salkhit_all_deam | Mbuti | -0.0043 | -0.65 | 0.0066 | 16,754 | 16,898 | 305,478 | Native American |
| GoyetQ116-1 | Pima | Salkhit_all_deam | Mbuti | -0.0017 | -0.25 | 0.0067 | 16,878 | 16,935 | 305,468 | Native American |
| GoyetQ116-1 | Clovis | Salkhit_all_deam | Mbuti | -0.0059 | -0.78 | 0.0075 | 16,631 | 16,829 | 305,152 | Native American |
| GoyetQ116-1 | Cree | Salkhit_all_deam | Mbuti | 0.0083 | 1.32 | 0.0063 | 16,798 | 16,520 | 305,482 | Native American |
| GoyetQ116-1 | Mixtec | Salkhit_all_deam | Mbuti | 0.0081 | 1.28 | 0.0063 | 16,989 | 16,716 | 305,477 | Native American |
| GoyetQ116-1 | Botocudo | Salkhit_all_deam | Mbuti | 0.0005 | 0.07 | 0.0073 | 16,434 | 16,418 | 295,932 | Native American |

**Table S15** *D*-statistics of the form *D(Goyet Q116-1, X, Salkhit, Mbuti)* where X is an ancient individual or present-day population (Pop 3). Significant *D*-statistics (|Z| > 2) are in red.

| Pop1 | Pop2 | Pop3 | Pop4 | D-Stat | Z | std.err | BABA | ABBA | #SNPs | Pop3 - origin |
|---|---|---|---|---|---|---|---|---|---|---|
| GoyetQ116-1 | Salkhit_all_deam | Tianyuan | Mbuti | -0.0703 | -8.81 | 0.0079 | 14,103 | 16,236 | 283,946 | ancient |
| GoyetQ116-1 | Yana1 | Tianyuan | Mbuti | -0.0126 | -1.74 | 0.0072 | 35,169 | 36,067 | 684,511 | ancient |
| GoyetQ116-1 | Yana2 | Tianyuan | Mbuti | -0.0088 | -1.23 | 0.0071 | 35,233 | 35,856 | 684,069 | ancient |
| GoyetQ116-1 | Malta1 | Tianyuan | Mbuti | 0.0037 | 0.49 | 0.0075 | 27,539 | 27,337 | 524,482 | ancient |
| GoyetQ116-1 | Sunghir3 | Tianyuan | Mbuti | 0.0146 | 2.06 | 0.0070 | 35,612 | 34,586 | 684,452 | ancient |
| GoyetQ116-1 | Kolyma1 | Tianyuan | Mbuti | -0.0315 | -4.64 | 0.0067 | 36,196 | 38,549 | 684,463 | ancient |
| GoyetQ116-1 | UstIshim | Tianyuan | Mbuti | 0.0197 | 2.71 | 0.0073 | 37,868 | 36,404 | 683,616 | ancient |
| GoyetQ116-1 | Kostenki14 | Tianyuan | Mbuti | 0.0353 | 5 | 0.0070 | 35,908 | 33,462 | 676,281 | ancient |
| GoyetQ116-1 | Vestonice16 | Tianyuan | Mbuti | 0.0221 | 3.11 | 0.0071 | 30,169 | 28,862 | 580,818 | ancient |
| GoyetQ116-1 | ElMiron | Tianyuan | Mbuti | 0.0181 | 2.59 | 0.0069 | 24,998 | 24,109 | 516,068 | ancient |
| GoyetQ116-1 | Saami | Tianyuan | Mbuti | 0.0082 | 1.43 | 0.0057 | 39,278 | 38,637 | 721,328 | West Eurasia |
| GoyetQ116-1 | French | Tianyuan | Mbuti | 0.0371 | 6.47 | 0.0057 | 40,010 | 37,147 | 721,347 | West Eurasia |
| GoyetQ116-1 | Russian | Tianyuan | Mbuti | 0.0254 | 4.46 | 0.0057 | 39,631 | 37,665 | 721,329 | West Eurasia |
| GoyetQ116-1 | Estonian | Tianyuan | Mbuti | 0.0336 | 5.54 | 0.0061 | 39,854 | 37,265 | 721,330 | West Eurasia |
| GoyetQ116-1 | Icelandic | Tianyuan | Mbuti | 0.0332 | 5.71 | 0.0058 | 39,608 | 37,061 | 721,325 | West Eurasia |
| GoyetQ116-1 | Finnish | Tianyuan | Mbuti | 0.0333 | 5.69 | 0.0058 | 39,913 | 37,344 | 721,342 | West Eurasia |
| GoyetQ116-1 | Czech | Tianyuan | Mbuti | 0.0384 | 6.13 | 0.0063 | 40,240 | 37,262 | 721,216 | West Eurasia |
| GoyetQ116-1 | Basque | Tianyuan | Mbuti | 0.0424 | 7.30 | 0.0058 | 40,069 | 36,806 | 721,321 | West Eurasia |
| GoyetQ116-1 | Spanish | Tianyuan | Mbuti | 0.0479 | 8.33 | 0.0057 | 40,436 | 36,740 | 721,328 | West Eurasia |
| GoyetQ116-1 | Sardinian | Tianyuan | Mbuti | 0.0451 | 8.09 | 0.0056 | 40,274 | 36,795 | 721,346 | West Eurasia |
| GoyetQ116-1 | English | Tianyuan | Mbuti | 0.0382 | 6.34 | 0.0060 | 40,081 | 37,130 | 721,326 | West Eurasia |
| GoyetQ116-1 | Anatolia_N | Tianyuan | Mbuti | 0.0433 | 7.97 | 0.0054 | 40,201 | 36,863 | 720,034 | Middle east |
| GoyetQ116-1 | Iraqi_Jew | Tianyuan | Mbuti | 0.0515 | 8.71 | 0.0059 | 41,294 | 37,252 | 721,325 | Middle east |
| GoyetQ116-1 | Palestinian | Tianyuan | Mbuti | 0.0662 | 11.83 | 0.0056 | 41,931 | 36,728 | 721,337 | Middle east |
| GoyetQ116-1 | Yi | Tianyuan | Mbuti | -0.0474 | -7.75 | 0.0061 | 38,132 | 41,925 | 721,327 | East Eurasia |
| GoyetQ116-1 | She | Tianyuan | Mbuti | -0.0417 | -6.82 | 0.0061 | 38,420 | 41,763 | 721,330 | East Eurasia |
| GoyetQ116-1 | Mongola | Tianyuan | Mbuti | -0.0391 | -6.21 | 0.0063 | 38,374 | 41,494 | 721,326 | East Eurasia |
| GoyetQ116-1 | Han | Tianyuan | Mbuti | -0.0444 | -7.62 | 0.0058 | 38,369 | 41,935 | 721,348 | East Eurasia |
| GoyetQ116-1 | Cambodian | Tianyuan | Mbuti | -0.0316 | -5.19 | 0.0061 | 38,812 | 41,345 | 721,320 | East Eurasia |
| GoyetQ116-1 | Korean | Tianyuan | Mbuti | -0.0411 | -6.67 | 0.0062 | 38,472 | 41,768 | 721,322 | East Eurasia |
| GoyetQ116-1 | Japanese | Tianyuan | Mbuti | -0.0421 | -7.11 | 0.0059 | 38,363 | 41,738 | 721,338 | East Eurasia |
| GoyetQ116-1 | Thai | Tianyuan | Mbuti | -0.034 | -5.61 | 0.0060 | 38,833 | 41,565 | 721,322 | East Eurasia |
| GoyetQ116-1 | Uygur | Tianyuan | Mbuti | -0.0024 | -0.4 | 0.0060 | 39,530 | 39,723 | 721,329 | East Eurasia |
| GoyetQ116-1 | Tibetan | Tianyuan | Mbuti | -0.0369 | -6.26 | 0.0059 | 38,488 | 41,440 | 721,320 | East Eurasia |
| GoyetQ116-1 | Dusun | Tianyuan | Mbuti | -0.0359 | -5.75 | 0.0062 | 38,760 | 41,650 | 721,314 | East Eurasia |
| GoyetQ116-1 | Igorot | Tianyuan | Mbuti | -0.0407 | -6.50 | 0.0062 | 38,540 | 41,811 | 721,302 | East Eurasia |
| GoyetQ116-1 | Hawaiian | Tianyuan | Mbuti | -0.0265 | -3.94 | 0.0067 | 39,232 | 41,370 | 721,149 | Oceania |
| GoyetQ116-1 | Papuan | Tianyuan | Mbuti | -0.0121 | -1.87 | 0.0064 | 39,712 | 40,686 | 721,351 | Oceania |
| GoyetQ116-1 | Bougainville | Tianyuan | Mbuti | -0.0116 | -1.79 | 0.0065 | 39,617 | 40,545 | 721,296 | Oceania |
| GoyetQ116-1 | Australian | Tianyuan | Mbuti | -0.012 | -1.83 | 0.0066 | 39,568 | 40,531 | 721,337 | Oceania |
| GoyetQ116-1 | Maori | Tianyuan | Mbuti | 0.0106 | 1.65 | 0.0064 | 39,746 | 38,912 | 721,162 | Oceania |
| GoyetQ116-1 | Mayan | Tianyuan | Mbuti | -0.0266 | -4.14 | 0.0064 | 38,544 | 40,653 | 721,310 | Native American |
| GoyetQ116-1 | Piapoco | Tianyuan | Mbuti | -0.0246 | -3.84 | 0.0064 | 38,756 | 40,712 | 721,307 | Native American |
| GoyetQ116-1 | Chipewyan | Tianyuan | Mbuti | -0.0336 | -5.34 | 0.0063 | 38,281 | 40,941 | 721,302 | Native American |
| GoyetQ116-1 | Surui | Tianyuan | Mbuti | -0.0354 | -5.41 | 0.0065 | 38,103 | 40,900 | 721,290 | Native American |
| GoyetQ116-1 | Zapotec | Tianyuan | Mbuti | -0.0264 | -4.12 | 0.0064 | 38,559 | 40,653 | 721,325 | Native American |
| GoyetQ116-1 | Quechua | Tianyuan | Mbuti | -0.024 | -3.89 | 0.0061 | 38,725 | 40,626 | 721,336 | Native American |
| GoyetQ116-1 | Karitiana | Tianyuan | Mbuti | -0.0295 | -4.68 | 0.0063 | 38,577 | 40,919 | 721,332 | Native American |
| GoyetQ116-1 | Chane | Tianyuan | Mbuti | -0.034 | -5.05 | 0.0067 | 38,265 | 40,960 | 721,224 | Native American |
| GoyetQ116-1 | Nahua | Tianyuan | Mbuti | -0.0213 | -3.06 | 0.0069 | 38,864 | 40,552 | 721,175 | Native American |
| GoyetQ116-1 | Mixe | Tianyuan | Mbuti | -0.0211 | -3.24 | 0.00651 | 38,963 | 40,639 | 721,327 | Native American |
| GoyetQ116-1 | Pima | Tianyuan | Mbuti | -0.0262 | -4.17 | 0.0063 | 38,781 | 40,865 | 721,301 | Native American |
| GoyetQ116-1 | Clovis | Tianyuan | Mbuti | -0.0256 | -3.7 | 0.0069 | 38,636 | 40,666 | 720,572 | Native American |
| GoyetQ116-1 | Cree | Tianyuan | Mbuti | -0.0126 | -2.12 | 0.0059 | 38,732 | 39,719 | 721,328 | Native American |
| GoyetQ116-1 | Mixtec | Tianyuan | Mbuti | -0.0052 | -0.84 | 0.0062 | 39,531 | 39,941 | 721,324 | Native American |
| GoyetQ116-1 | Botocudo | Tianyuan | Mbuti | -0.0315 | -5 | 0.0063 | 37,546 | 39,988 | 697,513 | Native American |

**Table S16** *D*-statistics of the form *D(Goyet Q116-1, X, Tianyuan, Mbuti)* where X is an ancient individual or present-day population (Pop 3). Significant *D*-statistics (|Z| > 2) are in red.

| Neandertal 1 | Outgroup1 | Target - modern human | Outgroup2 | Neandertal 1 | Outgroup1 | Neandertal2 | Outgroup2 | Neandertal ancestry (%) | std.err | Z score |
|---|---|---|---|---|---|---|---|---|---|---|
| Altai | Chimp | **Ust'Ishim_snpAD** | Mbuti | Altai | Chimp | Vindija33.19 | Mbuti | **2.11%** | 0.0041 | 5.18 |
| Altai | Chimp | **Yana1** | Mbuti | Altai | Chimp | Vindija33.19 | Mbuti | **2.34%** | 0.0041 | 5.74 |
| Altai | Chimp | **Yana2** | Mbuti | Altai | Chimp | Vindija33.19 | Mbuti | **2.72%** | 0.0039 | 6.92 |
| Altai | Chimp | **Kolyma1** | Mbuti | Altai | Chimp | Vindija33.19 | Mbuti | **1.97%** | 0.0037 | 5.23 |
| Altai | Chimp | **Salkhit_all_deam** | Mbuti | Altai | Chimp | Vindija33.19 | Mbuti | **1.73%** | 0.0048 | 3.63 |
| Altai | Chimp | **Tianyuan** | Mbuti | Altai | Chimp | Vindija33.19 | Mbuti | **1.71%** | 0.0043 | 3.99 |
| Altai | Chimp | **Oase1** | Mbuti | Altai | Chimp | Vindija33.19 | Mbuti | **6.36%** | 0.0077 | 8.20 |
| Altai | Chimp | **Kostenki14** | Mbuti | Altai | Chimp | Vindija33.19 | Mbuti | **1.75%** | 0.004 | 4.38 |
| Altai | Chimp | **GoyetQ116-1** | Mbuti | Altai | Chimp | Vindija33.19 | Mbuti | **2.60%** | 0.0046 | 5.64 |
| Altai | Chimp | **Muierii2** | Mbuti | Altai | Chimp | Vindija33.19 | Mbuti | **1.32%** | 0.0096 | 1.37 |
| Altai | Chimp | **KremsWA3** | Mbuti | Altai | Chimp | Vindija33.19 | Mbuti | **1.60%** | 0.0068 | 2.35 |
| Altai | Chimp | **Vestonice13** | Mbuti | Altai | Chimp | Vindija33.19 | Mbuti | **2.61%** | 0.008 | 3.27 |
| Altai | Chimp | **Vestonice16** | Mbuti | Altai | Chimp | Vindija33.19 | Mbuti | **1.45%** | 0.0045 | 3.26 |
| Altai | Chimp | **Malta1** | Mbuti | Altai | Chimp | Vindija33.19 | Mbuti | **2.24%** | 0.0041 | 5.45 |
| Altai | Chimp | **ElMiron** | Mbuti | Altai | Chimp | Vindija33.19 | Mbuti | **1.87%** | 0.0046 | 4.04 |
| Altai | Chimp | **Dai** | Mbuti | Altai | Chimp | Vindija33.19 | Mbuti | **1.90%** | 0.003 | 6.30 |
| Altai | Chimp | **Han** | Mbuti | Altai | Chimp | Vindija33.19 | Mbuti | **1.96%** | 0.0031 | 6.23 |
| Altai | Chimp | **Papuan** | Mbuti | Altai | Chimp | Vindija33.19 | Mbuti | **2.83%** | 0.0032 | 8.72 |
| Altai | Chimp | **French** | Mbuti | Altai | Chimp | Vindija33.19 | Mbuti | **1.49%** | 0.0029 | 5.15 |
| Altai | Chimp | **Bougainville** | Mbuti | Altai | Chimp | Vindija33.19 | Mbuti | **2.94%** | 0.0036 | 7.95 |

**Table S17** Proportion of Neandertal ancestry in some ancient and present-day human genomes estimated by *f4-ratio* statistics.

| Sample | #Tracks assigned significantly to Denisovan ancestry (tracks > 0.2 cM) |
|---|---|
| Salkhit | **18** |
| Tianyuan | **20** |
| Yana1 | **3** |
| Yana2 | **6** |
| Malta1 | **4** |
| Kolyma1 | **4** |
| Vestonice16 | **0** |
| Sunghir3 | **0** |
| Kostenki14 | **0** |
| Oase1 | **0** |
| UstIshim | **0** |

**Table S18** Number of genomic tracks assigned to Denisovans ancestry in the genomes of 12 ancient modern humans.

| Present-day populations | nbr of individual | #Denisovan Tracks > 0.05 cM | Overlap to Denisovan tracks in the Salkhit genome | | | | Overlap to Denisovan tracks in the Tianyuan genome | | | | Country | Geographic Region | Latitude coord. | Longitude coord. |
|---|---|---|---|---|---|---|---|---|---|---|---|---|---|---|
| | | | #Denisovan Tracks Overlapping | Correlation Coefficient of the overlap | CI Bootstrap | Significant Overlap | #Denisovan Tracks Overlapping | Correlation Coefficient of the overlap | CI Bootstrap | Significant Overlap | | | | |
| Abkhasian | 2 | 5 | 1 | 0.0352 | 0.0008 | Yes | 0 | -0.001 | 0.0127 | No | Russia | MiddleEastCaucasus | 43.003 | 41.015 |
| Adygei | 2 | 14 | 0 | -0.0008 | 0.0118 | No | 0 | -0.0011 | 0.0121 | No | Russia(Caucasus) | MiddleEastCaucasus | 44 | 39 |
| Albanian | 1 | 3 | 0 | -0.0004 | 0.0205 | No | 0 | -0.0006 | 0.0149 | No | Albania | WestEurasia | 41.3 | 19.8 |
| Aleut | 2 | 22 | 0 | -0.001 | 0.0057 | No | 0 | -0.0013 | 0.0161 | No | Russia | CentralAsiaSiberia | 55.18 | 166 |
| Altaian | 1 | 22 | 2 | 0.0946 | 0.0179 | Yes | 1 | 0.0744 | 0.0168 | Yes | Russia | CentralAsiaSiberia | 50.84 | 85.654 |
| Ami | 2 | 29 | 2 | 0.0372 | 0.0172 | Yes | 2 | 0.0363 | 0.0102 | Yes | Taiwan | EastAsia | 22.843 | 121.185 |
| Armenian | 2 | 3 | 0 | -0.0004 | 0.0172 | No | 0 | -0.0005 | 0.0161 | No | Armenia | MiddleEastCaucasus | 40.605 | 43.101 |
| Atayal | 1 | 17 | 0 | -0.0009 | 0.0153 | No | 0 | -0.0012 | 0.0127 | No | Taiwan | EastAsia | 24.611 | 121.296 |
| Australian | 5 | 550 | 0 | -0.0055 | 0.0175 | No | 1 | -0.006 | 0.0167 | No | Australia | Oceania | -13 | 143 |
| Balochi | 2 | 17 | 0 | -0.001 | 0.0208 | No | 0 | -0.0013 | 0.0148 | No | Pakistan | SouthAsia | 30.498 | 66.5 |
| Basque | 2 | 5 | 0 | -0.0004 | 0.0031 | No | 0 | -0.0005 | 0.0161 | No | France | WestEurasia | 43 | 0 |
| BedouinB | 2 | 5 | 0 | -0.0003 | 0.013 | No | 0 | -0.0004 | 0.0051 | No | Israel(Negev) | MiddleEastCaucasus | 31 | 35 |
| Bengali | 2 | 28 | 0 | -0.0012 | 0.007 | No | 1 | 0.0026 | 0.0144 | No | Bangladesh | SouthAsia | 23.7 | 90.4 |
| Bergamo | 2 | 5 | 0 | -0.0004 | 0.0008 | No | 0 | -0.0005 | 0.0019 | No | Italy(Bergamo) | WestEurasia | 46 | 10 |
| Bougainville | 2 | 491 | 2 | 0.0087 | 0.0156 | No | 1 | -0.0022 | 0.0161 | No | PapuaNewGuinea | Oceania | -6 | 155 |
| Brahmin | 2 | 24 | 0 | -0.0011 | 0.0081 | No | 0 | -0.0014 | 0.019 | No | India | SouthAsia | 17.7 | 83.3 |
| Brahui | 2 | 14 | 0 | -0.0009 | 0.0113 | No | 0 | -0.0012 | 0.0182 | No | Pakistan | SouthAsia | 30.5 | 66.5 |
| Bulgarian | 2 | 4 | 0 | -0.0004 | 0.0193 | No | 0 | -0.0005 | 0.0037 | No | Bulgaria | WestEurasia | 42.163 | 24.741 |
| Burmese | 2 | 36 | 1 | 0.0198 | 0.0115 | Yes | 1 | 0.004 | 0.0174 | No | Myanmar | EastAsia | 17 | 96.7 |
| Burusho | 2 | 24 | 0 | -0.001 | 0.0057 | No | 1 | 0.0845 | 0.0144 | Yes | Pakistan | SouthAsia | 36.5 | 74 |
| Cambodian | 2 | 38 | 2 | 0.118 | 0.0101 | Yes | 0 | -0.0018 | 0.0133 | No | Cambodia | EastAsia | 12 | 105 |
| Chane | 1 | 3 | 0 | -0.0004 | 0.0105 | No | 0 | -0.0005 | 0.0142 | No | Argentina | America | -22.53 | -63.82 |
| Chechen | 1 | 9 | 0 | -0.0007 | 0.0051 | No | 0 | -0.0009 | 0.0107 | No | Russia | MiddleEastCaucasus | 43.333 | 45.65 |
| Chukchi | 1 | 9 | 0 | -0.0007 | 0.0216 | Yes | 1 | 0.0105 | 0.0077 | Yes | Russia | CentralAsiaSiberia | 69 | 169 |
| Crete | 2 | 9 | 0 | -0.0004 | 0.004 | No | 0 | -0.0005 | 0.0144 | No | Greece | WestEurasia | 35.163 | 25.445 |
| Czech | 1 | 3 | 0 | -0.0005 | 0.0035 | No | 0 | -0.0006 | 0.0225 | No | Czechoslovia(pre1989) | WestEurasia | 50.1 | 14.4 |
| Dai | 4 | 69 | 3 | 0.0762 | 0.0134 | Yes | 3 | 0.0191 | 0.0147 | Yes | China | EastAsia | 21 | 100 |
| Daur | 1 | 19 | 0 | -0.0012 | 0.0149 | No | 0 | -0.0015 | 0.0177 | No | China | EastAsia | 48.5 | 124 |
| Druze | 2 | 5 | 0 | -0.0005 | 0.0 | No | 0 | -0.0007 | 0.0172 | No | Israel(Carmel) | MiddleEastCaucasus | 32 | 35 |
| Dusun | 2 | 47 | 3 | 0.109 | 0.0136 | Yes | 2 | 0.0228 | 0.014 | Yes | Brunei | SouthAsia | 4.7 | 114.7 |
| English | 2 | 2 | 0 | -0.0004 | 0.0005 | No | 0 | -0.0006 | 0.0005 | No | England | WestEurasia | 51.2 | 0.7 |
| Eskimo_Chaplin | 1 | 12 | 0 | -0.0007 | 0.0118 | No | 2 | 0.0265 | 0.0111 | Yes | Russia | CentralAsiaSiberia | 64.48 | 172.86 |
| Eskimo_Naukan | 2 | 30 | 0 | -0.0011 | 0.0138 | No | 1 | 0.0061 | 0.0157 | No | Russia | CentralAsiaSiberia | 66.02 | 169.71 |
| Eskimo_Sireniki | 2 | 27 | 0 | -0.0011 | 0.0085 | No | 1 | 0.0123 | 0.0142 | No | Russia | CentralAsiaSiberia | 64.4 | 173.9 |
| Estonian | 2 | 9 | 0 | -0.0005 | 0.0155 | No | 0 | -0.0007 | 0.004 | No | Estonia | WestEurasia | 58.985 | 26.864 |
| Even | 3 | 57 | 0 | -0.0014 | 0.0111 | No | 2 | 0.0399 | 0.0121 | Yes | Russia | CentralAsiaSiberia | 57.53 | 135.88 |
| Finnish | 3 | 14 | 0 | -0.0007 | 0.0082 | No | 0 | -0.0009 | 0.0149 | No | Finland | WestEurasia | 60.2 | 24.9 |
| French | 3 | 8 | 0 | -0.0004 | 0.0144 | No | 0 | -0.0005 | 0.0091 | No | France | WestEurasia | 46 | 2 |
| Georgian | 2 | 5 | 0 | -0.0005 | 0.0062 | No | 0 | -0.0006 | 0.0153 | No | Georgia | MiddleEastCaucasus | 42.5 | 41.85 |
| Greek | 2 | 2 | 0 | -0.0002 | 0.0011 | No | 0 | -0.0003 | 0.0012 | No | Greece | WestEurasia | 38 | 23.7 |
| Han | 3 | 62 | 1 | 0.0572 | 0.0175 | Yes | 1 | 0.0113 | 0.0164 | No | China | EastAsia | 32.3 | 114 |
| Hawaiian | 1 | 48 | 1 | 0.0408 | 0.0149 | Yes | 1 | 0.0149 | 0.0119 | Yes | USA | Oceania | 21.3 | -157.8 |
| Hazara | 2 | 25 | 0 | -0.0009 | 0.0088 | No | 0 | -0.0012 | 0.008 | No | Pakistan | SouthAsia | 33.5 | 70 |
| Hezhen | 2 | 33 | 2 | 0.0277 | 0.0085 | Yes | 2 | 0.0153 | 0.012 | Yes | China | EastAsia | 47.5 | 133.5 |
| Hungarian | 2 | 5 | 0 | -0.0005 | 0.004 | No | 0 | -0.0007 | 0.0147 | No | Hungary | WestEurasia | 47.5 | 19.1 |
| Icelandic | 2 | 9 | 0 | -0.0005 | 0.0043 | No | 0 | -0.0007 | 0.0128 | No | Iceland | WestEurasia | 64.1 | -21.9 |
| Igorot | 2 | 38 | 0 | -0.0015 | 0.012 | No | 1 | -0.0003 | 0.0142 | No | Philippines | Oceania | 17.1 | 121 |
| Iranian | 2 | 10 | 0 | -0.0005 | 0.0127 | No | 0 | -0.0007 | 0.0116 | No | Iran | MiddleEastCaucasus | 35.586 | 51.46 |
| Iraqi_Jew | 2 | 8 | 0 | -0.0007 | 0.0265 | No | 0 | -0.0009 | 0.0138 | No | Iraq | MiddleEastCaucasus | 33.3 | 44.4 |
| Irula | 2 | 49 | 0 | -0.0019 | 0.0162 | No | 0 | -0.0025 | 0.0139 | No | India | SouthAsia | 13.5 | 80 |
| Itelman | 1 | 18 | 1 | 0.0042 | 0.0113 | No | 0 | -0.0011 | 0.0175 | No | Russia | CentralAsiaSiberia | 57 | 157 |
| Japanese | 3 | 50 | 0 | -0.0017 | 0.0175 | No | 1 | 0.0075 | 0.0145 | No | Japan | EastAsia | 37.85 | 139 |
| Jordanian | 3 | 5 | 0 | -0.0002 | 0.0014 | No | 0 | -0.0003 | 0.0122 | No | Jordan | MiddleEastCaucasus | 32.055 | 35.914 |
| Kalash | 2 | 21 | 1 | 0.053 | 0.0122 | Yes | 1 | 0.0162 | 0.0135 | Yes | Pakistan | SouthAsia | 36 | 71.5 |
| Kapu | 2 | 25 | 0 | -0.0011 | 0.0086 | No | 0 | -0.0015 | 0.0147 | No | India | SouthAsia | 17.7 | 83.3 |
| Karitiana | 3 | 42 | 1 | 0.0024 | 0.0189 | No | 1 | 0.0058 | 0.0165 | No | Brazil | America | -10 | -63 |

**Table S19A** Overlap of Denisovan ancestry tracks between Early East Asians and present-day populations.

| Present-day populations | nbr of individual | #Denisovan Tracks > 0.05 cM | Overlap to Denisovan tracks in the Salkhit genome | | | | Overlap to Denisovan tracks in the Tianyuan genome | | | | Country | Geographic Region | Latitude coord. | Longitude coord. |
|---|---|---|---|---|---|---|---|---|---|---|---|---|---|---|
| | | | #Denisovan Tracks Overlapping | Correlation Coefficient of the overlap | CI Bootstrap | Significant Overlap | #Denisovan Tracks Overlapping | Correlation Coefficient of the overlap | CI Bootstrap | Significant Overlap | | | | |
| Khonda_Dora | 1 | 15 | 0 | -0.0008 | 0.0112 | No | 0 | -0.0011 | 0.0175 | No | India | SouthAsia | 18.3 | 82.9 |
| Kinh | 2 | 35 | 2 | 0.0215 | 0.0099 | Yes | 1 | 0.0023 | 0.0155 | No | Vietnam | EastAsia | 21 | 105.9 |
| Korean | 2 | 31 | 2 | 0.078 | 0.0111 | Yes | 1 | 0.0162 | 0.0126 | Yes | Korea | EastAsia | 37.6 | 127 |
| Kusunda | 2 | 41 | 1 | 0.001 | 0.0151 | No | 2 | 0.0253 | 0.0229 | Yes | Nepal | SouthAsia | 28.07 | 84.25 |
| Kyrgyz | 2 | 23 | 0 | -0.001 | 0.0167 | No | 0 | -0.0013 | 0.0112 | No | Kyrgyzystan | CentralAsiaSiberia | 42.9 | 74.6 |
| Lahu | 2 | 32 | 1 | 0.0248 | 0.0176 | Yes | 2 | 0.0127 | 0.0193 | No | China | EastAsia | 22 | 100 |
| Lezgin | 2 | 4 | 0 | -0.0006 | 0.0209 | No | 0 | -0.0009 | 0.0141 | No | Russia | MiddleEastCaucasus | 42.117 | 48.183 |
| Madiga | 2 | 40 | 0 | -0.0015 | 0.0165 | No | 0 | -0.002 | 0.0135 | No | India | SouthAsia | 17.7 | 83.3 |
| Makrani | 2 | 10 | 0 | -0.0006 | 0.0053 | No | 0 | -0.0008 | 0.0132 | No | Pakistan | SouthAsia | 26 | 64 |
| Mala | 2 | 36 | 0 | -0.0014 | 0.0187 | No | 0 | -0.0019 | 0.0132 | No | India | SouthAsia | 17.7 | 83.3 |
| Mansi | 2 | 20 | 0 | -0.0011 | 0.0158 | No | 2 | 0.0179 | 0.015 | Yes | Russia | CentralAsiaSiberia | 63.65 | 62.1 |
| Maori | 1 | 42 | 0 | -0.0015 | 0.0161 | No | 0 | -0.002 | 0.0137 | No | NewZealand | Oceania | -41.3 | 174.5 |
| Mayan | 2 | 34 | 0 | -0.0013 | 0.0183 | No | 0 | -0.0017 | 0.0163 | No | Mexico | America | 19 | -91 |
| Miao | 2 | 33 | 1 | 0.1367 | 0.0154 | Yes | 1 | 0.0015 | 0.0237 | No | China | EastAsia | 28 | 109 |
| Mixe | 3 | 53 | 0 | -0.0013 | 0.0153 | No | 0 | -0.0017 | 0.0149 | No | Mexico | America | 16.95 | -96.58 |
| Mixtec | 2 | 24 | 0 | -0.001 | 0.0123 | No | 0 | -0.0014 | 0.0128 | No | Mexico | America | 17 | -97 |
| Mongola | 2 | 46 | 3 | 0.095 | 0.0126 | Yes | 0 | -0.0021 | 0.0142 | No | China | EastAsia | 45 | 111 |
| Naxi | 3 | 65 | 0 | -0.0018 | 0.0152 | No | 3 | 0.011 | 0.0177 | No | China | EastAsia | 26 | 100 |
| North_Ossetian | 2 | 5 | 0 | -0.0004 | 0.0051 | No | 0 | -0.0005 | 0.0045 | No | Russia | MiddleEastCaucasus | 43.017 | 44.65 |
| Norwegian | 1 | 1 | 0 | -0.0001 | 0.0 | No | 0 | -0.0001 | 0.0047 | No | Norway | WestEurasia | 60.4 | 5.4 |
| Orcadian | 2 | 3 | 0 | -0.0002 | 0.0 | No | 0 | -0.0003 | 0.0097 | No | OrkneyIslands | WestEurasia | 59 | -3 |
| Oroqen | 2 | 43 | 1 | 0.0026 | 0.0107 | No | 2 | 0.0094 | 0.0125 | No | China | EastAsia | 50.4 | 126.5 |
| Palestinian | 3 | 6 | 0 | -0.0006 | 0.0107 | No | 0 | -0.0008 | 0.004 | No | Israel(Central) | MiddleEastCaucasus | 32 | 35 |
| Papuan | 15 | 3958 | 3 | -0.0046 | 0.0232 | No | 3 | -0.0064 | 0.0231 | No | PapuaNewGuinea | Oceania | -4 | 143 |
| Pathan | 2 | 27 | 0 | -0.0014 | 0.0134 | No | 0 | -0.0018 | 0.0165 | No | Pakistan | SouthAsia | 33.5 | 70.5 |
| Piapoco | 2 | 28 | 1 | 0.0007 | 0.0119 | No | 1 | -0.0012 | 0.0128 | No | Colombia | America | 3 | -68 |
| Pima | 2 | 28 | 0 | -0.001 | 0.021 | No | 1 | 0.0064 | 0.014 | No | Mexico | America | 29 | -108 |
| Polish | 1 | 3 | 0 | -0.0004 | 0.0009 | No | 0 | -0.0005 | 0.0122 | No | Poland | WestEurasia | 52.2 | 21 |
| Punjabi | 4 | 67 | 0 | -0.002 | 0.0168 | No | 1 | 0.0276 | 0.0126 | Yes | Pakistan | SouthAsia | 31.5 | 74.3 |
| Quechua | 3 | 42 | 0 | -0.0012 | 0.0116 | No | 0 | -0.0016 | 0.0136 | No | Peru | America | -13.5 | -72 |
| Relli | 2 | 27 | 2 | 0.0091 | 0.0242 | No | 0 | -0.0021 | 0.022 | No | India | SouthAsia | 17.7 | 83.3 |
| Russian | 2 | 10 | 0 | -0.0005 | 0.0124 | No | 0 | -0.0007 | 0.0161 | No | Russia | WestEurasia | 61 | 40 |
| Saami | 2 | 14 | 0 | -0.0007 | 0.01 | No | 0 | -0.001 | 0.0127 | No | Finland | WestEurasia | 69.9 | 27 |
| Samaritan | 1 | 3 | 0 | -0.0002 | 0.0069 | No | 0 | -0.0002 | 0.0057 | No | Israel | MiddleEastCaucasus | 32.203 | 35.27 |
| Sardinian | 3 | 5 | 0 | -0.0003 | 0.0011 | No | 0 | -0.0004 | 0.003 | No | Italy(Sardinia) | WestEurasia | 40 | 9 |
| She | 2 | 40 | 2 | 0.0282 | 0.0167 | Yes | 2 | 0.0258 | 0.0138 | Yes | China | EastAsia | 27 | 119 |
| Sindhi | 2 | 24 | 1 | 0.005 | 0.0141 | No | 0 | -0.0015 | 0.0217 | No | Pakistan | SouthAsia | 25.5 | 69 |
| Spanish | 2 | 7 | 0 | -0.0005 | 0.0167 | No | 0 | -0.0006 | 0.0099 | No | Spain | WestEurasia | 39.9 | -4 |
| Surui | 2 | 31 | 0 | -0.0009 | 0.0138 | No | 0 | -0.0012 | 0.0123 | No | Brazil | America | -11 | -62 |
| Tajik | 2 | 8 | 0 | -0.0007 | 0.0108 | No | 0 | -0.0009 | 0.0116 | No | Tajikistan | WestEurasia | 37.508 | 71.56 |
| Thai | 2 | 29 | 1 | 0.018 | 0.0107 | Yes | 1 | 0.0267 | 0.0162 | Yes | Thailand | EastAsia | 13.8 | 100.5 |
| Tlingit | 2 | 12 | 0 | -0.0008 | 0.0114 | No | 0 | -0.0011 | 0.0147 | No | Russia | CentralAsiaSiberia | 55.18 | 166 |
| Tu | 2 | 21 | 4 | 0.2219 | 0.0161 | Yes | 0 | -0.0016 | 0.0131 | No | China | EastAsia | 36 | 101 |
| Tubalar | 2 | 39 | 2 | 0.0579 | 0.0114 | Yes | 1 | -0.0005 | 0.01 | No | Russia | CentralAsiaSiberia | 51.13 | 87 |
| Tujia | 2 | 59 | 1 | 0.0157 | 0.0129 | No | 1 | 0.0186 | 0.0198 | No | China | EastAsia | 29 | 109 |
| Turkish | 2 | 8 | 0 | -0.0004 | 0.0033 | No | 0 | -0.0005 | 0.0094 | No | Turkey | MiddleEastCaucasus | 38.7 | 35.5 |
| Tuscan | 2 | 3 | 0 | -0.0005 | 0.0249 | No | 0 | -0.0006 | 0.019 | No | Italy(Tuscany) | WestEurasia | 43 | 11 |
| Ulchi | 2 | 44 | 1 | 0.0018 | 0.0141 | No | 1 | 0.0117 | 0.0151 | No | Russia | CentralAsiaSiberia | 52.37 | 140.45 |
| Uygur | 2 | 26 | 1 | 0.0005 | 0.0164 | No | 2 | 0.031 | 0.0156 | Yes | China | EastAsia | 44 | 81 |
| Xibo | 2 | 39 | 1 | 0.0123 | 0.0178 | No | 4 | 0.0329 | 0.0116 | Yes | China | EastAsia | 43.5 | 81.5 |
| Yadava | 2 | 28 | 1 | 0.0194 | 0.0084 | Yes | 0 | -0.0018 | 0.0153 | No | India | SouthAsia | 17.7 | 83.3 |
| Yakut | 2 | 36 | 0 | -0.0014 | 0.0208 | No | 1 | 0.0092 | 0.0136 | No | Russia | CentralAsiaSiberia | 63 | 129.5 |
| Yemenite_Jew | 2 | 5 | 0 | -0.0003 | 0.0165 | No | 0 | -0.0003 | 0.0024 | No | Yemen | WestEurasia | 15.4 | 44.2 |
| Yi | 2 | 47 | 0 | -0.0015 | 0.0114 | No | 1 | 0.0096 | 0.0121 | No | China | EastAsia | 28 | 103 |
| Zapotec | 2 | 22 | 0 | -0.001 | 0.0172 | No | 1 | 0.0064 | 0.0187 | No | Mexico | America | 16.5 | -97.2 |

**Table S19A** Overlap of Denisovan ancestry tracks between Early East Asians and present-day populations.
